# Pre-Settlement Forests around Puget Sound: Eyewitness Evidence

**DOI:** 10.1101/592733

**Authors:** Tom Schroeder

## Abstract

Witness trees from GLO surveys covering 6,300 square miles around Puget Sound (western Washington State) reveal, for the first time, the character and local diversity in the region’s mid-19^th^-century forest cover, before it was severely logged during the settlement period. Although only a few coniferous and hardwood species occurred overall, discrete geographical areas supported distinctive species compositions. Geo-climatic and developmental factors are explored to explain these local differences. Profiles of tree diameters reveal that most trees were small to medium in size, even though most areas also exhibited a minority of larger legacy trees. Approximate stand ages and stages of ecological succession are inferred from local tree sizes and site qualities. Despite current inclusion within the Western Hemlock Zone, major portions of the Puget lowlands (below 1000 feet) displayed extremely few western hemlock, the putative climax species, even stands of decidedly advanced age; in extreme cases “pioneer old growth” prevailed. Conversely, hemlock was strongly predominant in the region’s framing foothills (above 1000 feet), even when stand age there was relatively low. These problematic successional conditions question whether many early forests around Puget Sound deserve a categorical recognition separate from the Western Hemlock Zone.

## Introduction

Surprisingly little is known with certainty about the pre-settlement forests around western Washington’s Puget Sound. No direct scientific description had been undertaken before the original forests had been thoroughly cut over, which by about 1960 had consumed forests across the lowlands and into the flanking foothills. The present investigation offers a retrospective glimpse into the early vegetation through the lens of 19^th^-century land surveys. The surveys sampled and documented trees across the region; the uniformity and comprehensiveness of those records allows a broad, if sometimes sketchy, reconstruction of the region’s entire forested landscape.

In 1853, the inaugural year for Washington Territory, the U.S. General Land Office (GLO, now Bureau of Land Management) began imposing a rectangular grid on the land. While crisscrossing the region, GLO surveyors fixed the grid by determining and marking intersect corners with wooden posts. (For more procedural details about GLO surveys, see Appendix A.) They enlisted natural trees to “witness” their grid lines and corners; witness trees served as convenient way-finders for section corners and so-called quarter corners halfway between, even if the posts disintegrated. A contractual survey unit generally covered one township, a six-by-six-mile square of 36 sections, whose geographic location was identified by Township and Range (T/R) coordinates. Because field notes containing witness-tree data and other surveying details were compiled and archived by township, this study’s descriptions and analyses are usually unitized by township. Puget Sound country contains over two hundred townships.

GLO instructions specified that witness trees should be whichever trees happened to lie closest to the reference post: four trees per section corner (the nearest tree in each compass quadrant) and two per quarter corner (on either side of the section line). This procedure resulted in 24 witness trees around a section’s boundaries: eight trees *within* the section and the remaining 16 trees belonging to adjacent sections. Accordingly, a 36-square-mile township featured as many as 288 witness trees. Each recorded tree was identified by type (common name), diameter at chest height (in inches), and location relative to the post. Because of the manner of their selection, corner and quarter-corner witness trees are generally understood to be relatively free from surveyors’ biases and thus randomly representative of their surroundings, the rationale being that the artificial grid, as well as any proximate tree, bore no predictable relationship to the surrounding landscape. Even if randomly designated, however, witness trees sampled their forest surroundings at a very low rate (order of magnitude 0.01%), so the spatial resolution of any forest reconstruction based on witness trees will necessarily also be very low, just as all conclusions will be inferential.

The purpose of GLO surveys was to subdivide the land so that it could be sold or homesteaded, so it was important for the surveys to *precede* arriving settlers. Over a period of four decades, surveyors marked thousands of miles of straight lines through the largely trackless forests, beginning at Puget Sound’s shorelines and progressing inland into the lower foothills of the flanking Cascade and Olympic Mountains. For the purposes of the present study, those lower foothills define the geographical boundaries of Puget Sound country – to elevations of about 2000 feet in the north and 1000 feet in the south – and coincide with the limits of glacial soils deposited by a lobe of continental ice that intruded into Puget Basin ∼15,000 years ago. Defined in that way, the region encompasses about 6,300 sq miles of land.

## Region-wide Results

For this study, information from 49,746 witness trees was extracted from 14,000 handwritten pages of GLO field notes (archived and available at BLM online) and compiled into spreadsheets for later sorting and filtering. Collected information included tree species, tree diameters, locations within sections and townships, associated landscape observations, field-note source pages, and dates of surveys.

### Quantification by Species

Across the region, there were fewer conifer species than hardwoods in the pre-settlement forests (9 species versus 24, all identified in Appendix C), but the grand tally of coniferous specimens vastly outnumbered hardwoods (relative frequencies of occurrence 79% versus 21%). Abundances of the principal species around Puget Sound are displayed in Figure 1; one-third of all trees were Douglas-fir, and another one-half were either western redecedar or western hemlock. Together, the five principal species (the above plus red alder and bigleaf maple) accounted for 86% of all trees. Witness-tree diameters ranged from 2 to 180 inches, with an overall average diameter (linear mean) of 16.5 inches. The average diameter of all conifers (18.5 inches) was twice that of all hardwoods (9.1 inches).

**Figure 1.**
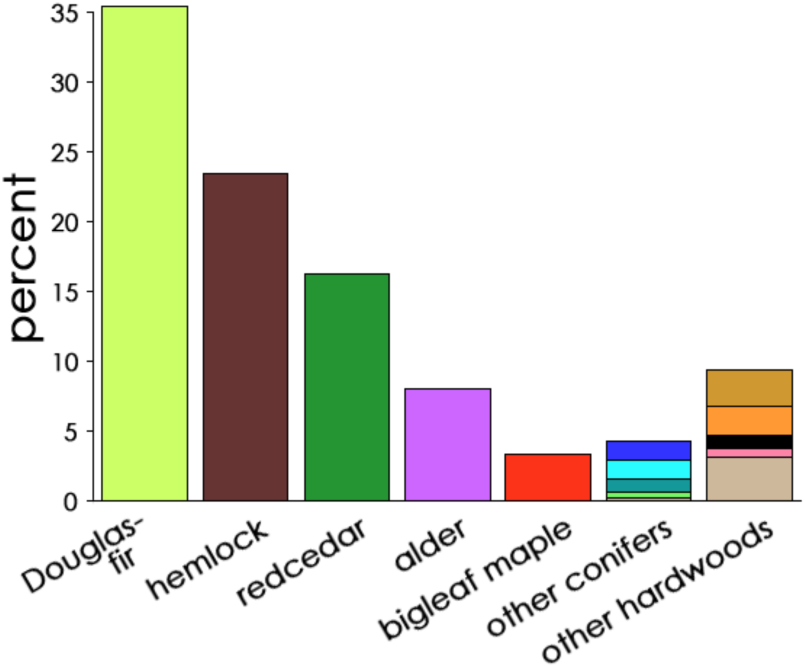
Relative frequencies of species in the total pool of witness trees for Puget Sound country. The top three conifers and two hardwoods are individually identified. Color codes are standard for this study.

### Diameter Profiles

Tree diameters within total populations of the five most abundant species are displayed in Figure 2. All diameter-distribution histograms are left-skewed, indicating that in each case more witness trees were smaller than the species’ average diameter rather than larger; even so, in each instance a few trees were exceptionally large, notably redcedar (to 180 inches, or 15 feet in diameter!) and Douglas-fir (to 100 inches, or 8 feet). Among the principal confers, redcedar were largest in average diameter, but they were least in abundance.

**Figure 2.**
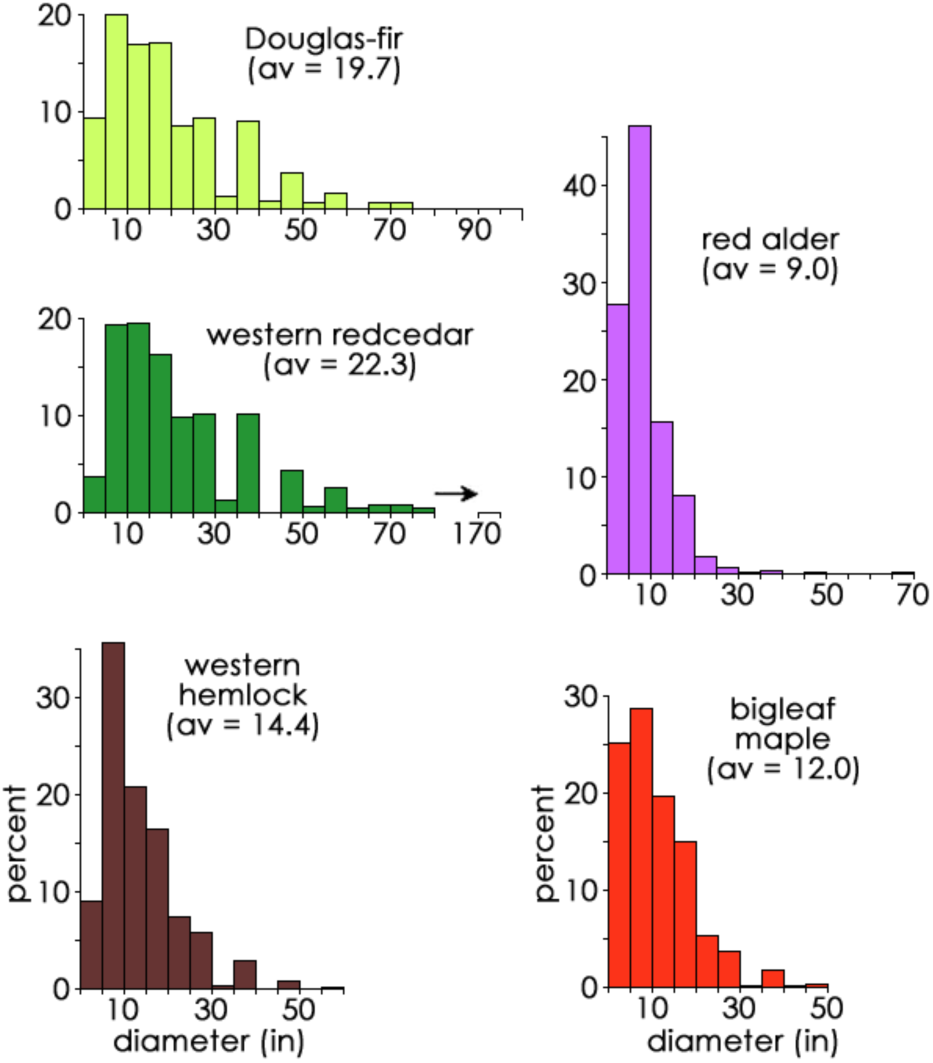
Histograms of diameter distributions for the top tree species. Values are binned in 5-inch intervals, e.g., 1-5, 6-10, etc. Histogram forms are all left-skewed, signifying that small specimens were more numerous than large specimens. av, region-wide linear average diameters, in inches.

Histograms in Figure 2 notably exhibit low-value gaps at the 25-29-inch and 41-45-inch bins. These gaps are artifactual and do not indicate size- or age-segregated cohorts of trees; it is likely that surveyors displayed “integer biases” while determining tree diameters, especially when estimating large diameters, by rounding up or down to 24, 30, 40, or 46 instead of less “popular” intervening integers.

### Witness Tree Map

Figure 3 displays the geographic distribution of 49,746 witness trees color-coded by species. Conifers are portrayed mostly in greens or dark brown, hardwood species in oranges and reds. By scanning the map for general color patterns, it is readily evident that early forests around Puget Sound were not compositionally uniform across the region. Species were unevenly distributed: Douglas-fir (chartreuse) was hyper-abundant in some places; hemlock (brown) was concentrated in others; and willow (orange) and alder (magenta) were arranged in various configurations. The entire area south of Lake Washington (T25/4E) contained very little hemlock, except in the foothills at the region’s margins; in contrast, pine (pale blue, likely signifying mostly lodgepole pine) occurred throughout the southern lowlands, as well as in the San Juans, yet pines were rare elsewhere. Big leaf maple (red) was common throughout the region’s eastern half, but scarce in the western half. Garry oak (black) was concentrated near the south end. Two minor hardwoods (coded tan among “other species” and therefore not separately visible in Figure 3) occupied mutually exclusive zones: paper birch north of T32 and Oregon ash only south of T27.

**Figure 3.**
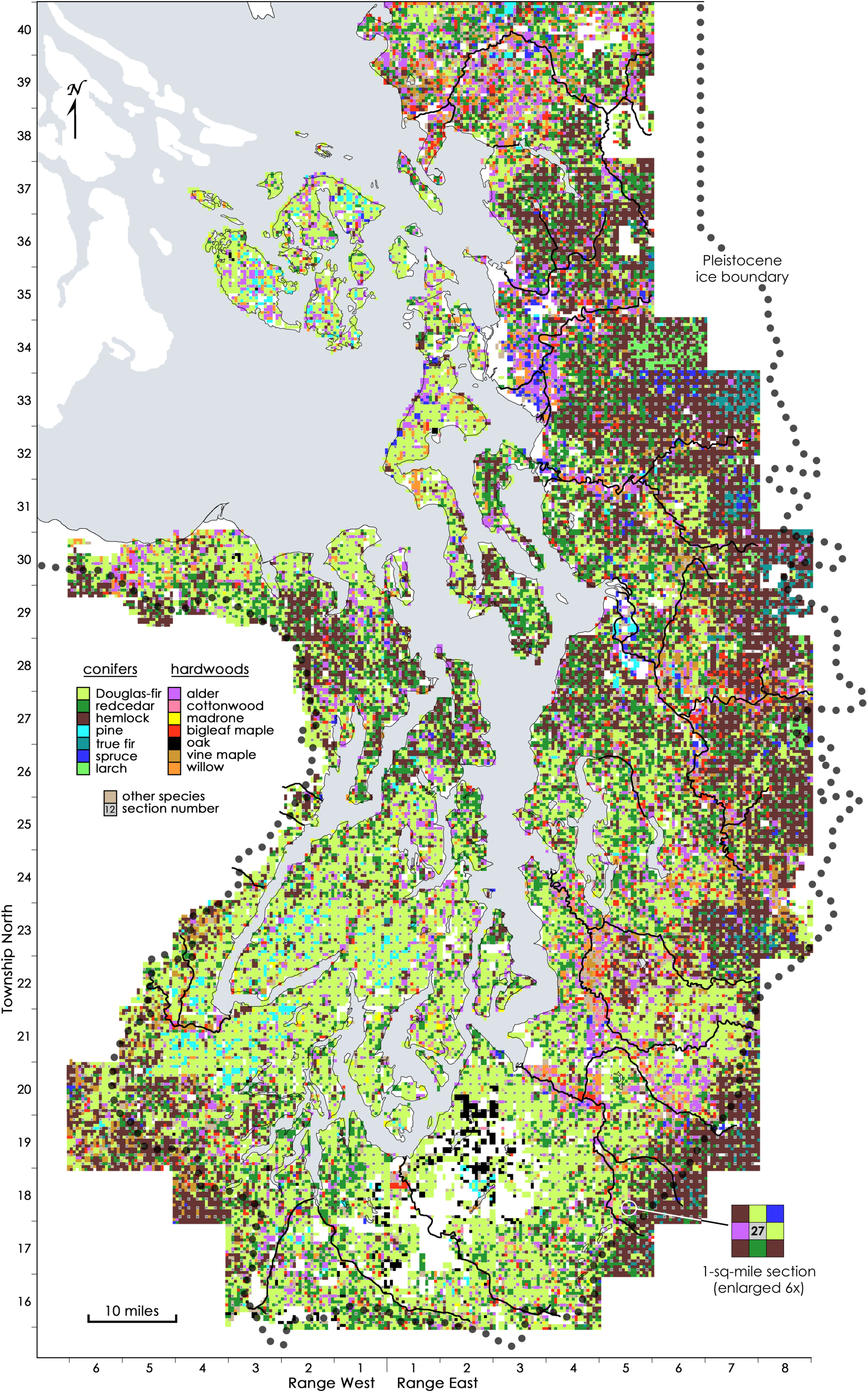
(This map withstands high magnification.) Species and locations of 49,746 GLO witness trees from the late-pre-settlement forests around Puget Sound. Inset, color-coded trees in a survey section.

The map in Figure 3 should be regarded as a thematic map on a rectilinear projection; it was not designed to conform geodetically. It was prepared by hand in a graphic program using colorized small squares to indicate the species of individual witness trees (see Appendix D for methodological details). For any mile-square survey section, such as that represented by the inset in Figure 3, eight witness trees (four each at corners and sides) are arranged around a central section identifier, all as directed by GLO field notes. The map was prepared in township units. When completed, townships units were tiled together according to their T/R coordinates to form the final display, which was then overlain by a standard shoreline outline of Puget Sound, including major rivers and bodies of freshwater.

## Distinctive Local Areas

Heterogeneity of forest composition is evident in Figure 3, and a few foci of particular differentiation stand out. Five locally specialized forest areas – assigned nicknames for reference – plus the less differentiated interstitial matrix are identified in Figure 4. Two areas, so-called South Prairies and SW Colonizers, are both coherent units, whereas Floodplains, Rainshadow, and Hemlock Heights are composites of separate portions with shared species makeups. The Matrix Forest was composed of transitional mixes of species.

**Figure 4.**
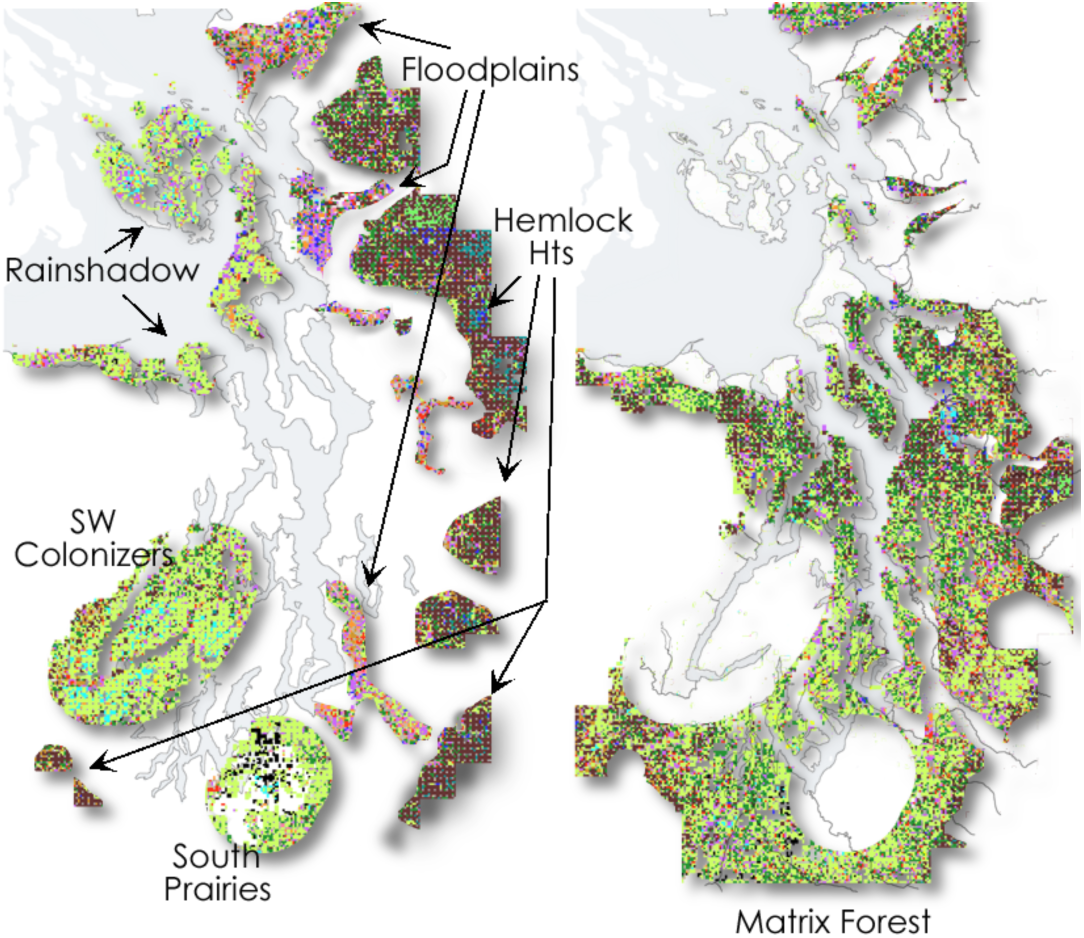
Dissected map that isolates and identifies distinctive forest areas. Appendix E lists and locates the diagnostic townships chosen to represent each area, from which tree data were pooled.

For quantitative characterizations of witness-tree populations within an area, data were pooled from a subset of core townships judged to be representative (specified in Appendix E). Graphic displays of quantitative witness-tree parameters from those area pools, excluding Floodplains, are shown together in Figure 5 to facilitate comparisons between areas.

**Figure 5.**
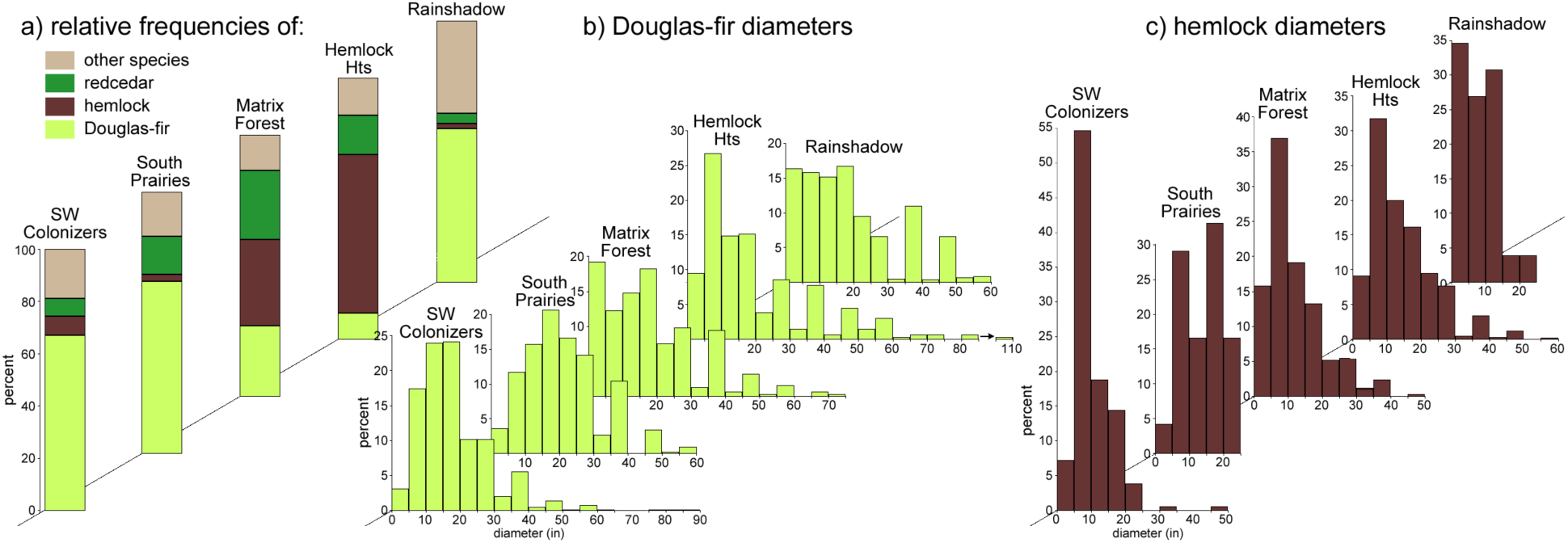
Quantitative attributes of area forests displayed along trending z-axes to aid comparisons (charted areas align horizontally). It can be argued that the mainland areas (the first four) follow a developmental gradient, whereas mostly insular Rainshadow does not conform to such a scheme.

### Floodplains

According to Figure 3, discrete ribbons of (mostly) hardwoods accompany the lower reaches of major rivers as they cross Puget lowlands while draining the Cascades. Moisture-tolerating vegetation occupied the broad alluvial accumulations deposited over millennia by silt-laden mountain torrents as they discharged onto and slowed crossed the coastal plains. Flood plains were generally more extensive in the region’s north, presumably because Cascade watersheds were larger there; the Nooksack and Skagit Rivers (T38-39/R2-4E and T33-35/R3-4E, respectively) developed very broad flood plains, in part due to ancient, mile-long, semi-permanent logjams that deflected and dispersed flow.

To analyze the vegetation in greater detail, a witness-tree database was compiled from a few townships of the Nooksack and Puyallup River flood plains that were more thoroughly filled with the distinctive mix of hardwoods. That database shows that half of the trees were fast-growing, short-lived hardwoods, primarily red alder, bigleaf maple, and willow, most of which were less than 10 inches in diameter; undoubtedly, Floodplains vegetation experienced high turnover caused by periodic destructive flooding and the sheer hydraulics of water flow. Some larger-diameter redcedar, Douglas-fir, and cottonwood also occurred and may have represented older trees persisting on more stable sub-sites, such as riverside benches, hummocks of elevated ground, and perhaps even on the old logjams. Medium-size Sitka spruce were uniquely common on Skagit River’s very low-lying alluvial fan, perhaps facilitated by elevated magnesium levels from nearby seawater.

Floodplains and their transitory vegetation accounted for less than 5% of land area around Puget Sound. Though undeniably present as part of the region’s diverse land cover, Floodplains vegetation is here considered to be an outlier of proto-forest distinct from more established forest types and thus are afforded little further attention in this study. Nevertheless, from the perspective of incoming settlers intent on farming, nutrient-rich bottom lands represented agricultural lands of high quality and ready accessibility; clearing away small hardwoods would not have been arduous, yet drainage projects sufficient to protect against the ravages of annual flooding required significant communal engineering.

### Rainshadow

The unusual forests of the region’s farthest northwest ironically shared a feature with vegetation of the flood plains of the Puget Lowlands – a preponderance of pioneers. Instead of the moisture-adapted species of flood plains, however, Rainshadow pioneers were drought-resistant species, well adapted to severe summer droughts in landscapes that lacked year-round rivers altogether.

This area occupies a zone of low precipitation and relative cloudlessness, the Olympic Rain Shadow (see Appendix A). This 40-mile-wide zone is centered over the marine straits, but it affects the north coast of the Olympic Peninsula, San Juan Archipelago, and the northern half of Whidbey Island (T33/ R2E), an aggregate land area of about 350 sq miles. Precipitation is only 20 inches on the north coast of the Olympic Peninsula, rising to 25-35 inches over north Whidbey and the San Juans, (higher on Orcas Island’s Mt. Constitution). The growing season is so severely shortened by intense summer drought that many tree species are unable to survive. As dry-land specialists, Douglas-fir and lodgepole pine can tolerate the drought stress, but even their rates of growth were likely reduced here, relative to other areas.

The database chosen to characterize Rainshadow forests is the pool of San Juan County’s witness trees, representing an irregular land area equivalent to about five townships (175 sq miles). Three pioneer species accounted for 83% of all stems in the database (Figure 5a): Douglas-fir (58.7%), lodgepole pine (9.1%), and red alder (15.9%); of these, only Douglas-fir is notably fire-resistant, thanks to its thicker bark. Redcedar and hemlock, so abundant elsewhere around Puget Sound, were uncommon (<4% and <2%, respectively). Some local differences are noted from other parts of Rainshadow; for example, north Whidbey Island had far fewer pine and alder than the San Juans.

The average diameter of Rainshadow Douglas-fir was 19.4 inches, similar to the regional average, despite their depressed rate of growth. Pine and alder diameters averaged only 6.9 and 8.2 inches; these two species are short-lived and fire-sensitive, so their very presence in 1874, the year of the survey, suggests that less than a century had elapsed, on average, since a tree-killing fire disturbance, at least locally where those species occurred.

Douglas-fir’s diameter histogram (Figure 5b) reveals equal populations of small- and mid-sized trees, as well as several larger specimens, indicating either a) that younger cohorts were intimately intermingled with considerably older specimens, or b) that the landscape was a geographic mosaic of stands of diverse ages. This distribution of diameters is unlike Douglas-fir populations in other distinct areas, and it may signify that local parts of Rainshadow experienced episodic opportunities for Douglas-fir establishment, such as relatively frequent, small-scale wildfires, as recently reported by Bakker *et al*. (2019). Diameters of the uncommon hemlock (Figure 5c) were concentrated in the 1-5-inch class, again suggesting either that wildfire had intervened in the recent past or, alternatively, that hemlock growth was older than it seemed but severely stunted by low soil moisture.

Besides its unique forest composition, the Rainshadow area was also distinguished by the presence of several small and a few medium-sized treeless prairies. Early settlers were strongly attracted to such sites in north Whidbey, Sequim Prairie (T30/R2W), and San Juan Islands by providing immediate pasturage for livestock and prospects for grain production. For untold centuries prior to settlement, these prairies and numerous beach landings were used by competing groups of Native people, and it is presumed that anthropogenic wildfires fairly frequently swept through these dry landscapes, periodically and locally suppressing fire-sensitive tree species (Bakker *et al*., 2019). In some cases the absence of trees is traceable to excessively drained soil types, which exacerbate already harsh droughtiness.

### Hemlock Hts

Land above about 1000 feet at the margins of Puget Sound country displayed heavier concentrations of hemlock than anywhere in the lowlands. Hemlock predominance covered about 800 sq miles (13% of the region) and included foothills of the North Cascades (T30/R8E), Chuckanut Mountains (T33/R5E), northeast Olympics (T28/R1W), and the somewhat lower foothills of the south-central Cascades and Black Hills (T18/R4W). Based on twelve widely separated townships chosen as the database, hemlock’s relative frequency was 60.6% (Figure 5a), whereas redcedar and Douglas-fir abundance was only 15.3% and 10.0%, respectively. Though not shown, hemlock was undoubtedly also concentrated on the east-facing slopes of the Olympic Mountains (which rose too abruptly to be surveyed) and higher into the west-facing slopes of the Cascades, beyond the bounds of this study.

The average diameter of hemlock specimens in the database was 15.4 inches, exceeded by redcedar and Douglas-fir (19.9 and 19.1 inches). Histograms of both hemlock and Douglas-fir diameters (Figure 5a and b) are both left-skewed and broad, indicating a wide range of diameters, mostly small but including minor sub-populations of large specimens (up to 60 inches for hemlock and 110 inches for Douglas-fir). Redcedar’s diameter range (not shown) was similarly broad and included a few trees 100-180 inches in diameter (8-15 feet). Although all of these long-lived species are potentially capable of very great diameter growth, given enough disturbance-free time, their reportedly modest average dimensions suggest that the major components of Hemlock Hts stands were decidedly not ancient when surveyed, despite having some large legacy specimens that were likely centuries older.

Conditions for tree growth for the species found in Hemlock Hts must have been well-suited to the principal conifers. The area’s rising topography stimulates heavy precipitation (70-100 inches annually) by re-intercepting humid oceanic air after crossing the lowlands. Likewise, increased cloudiness and moderated summer temperatures would have encouraged moist soil conditions, prolonged the growing season, and reduced the area’s susceptibility to wildfire. Such conditions would have benefited drought-intolerant hemlock and supported ecological succession, all of which may justify hemlock’s high concentrations; however, if succession had indeed advanced significantly, then hemlock’s average diameter should have been greater than the meager 15.4 inches.

### South Prairies

Vegetation of this ∼300-sq-mile area straddling the Nisqually River was, and to a reduced extent remains, a patchwork of treeless bunch-grass prairie, sparsely treed oak-savanna, and conifer-encroached prairie. It also featured seasonal streams, shallow ponds, and small mucky swamps, many of which are vividly depicted in a GLO-commissioned map from 1855 (WSOS online; Anderson, 2012). Treeless prairies covered about 75 sq miles at the area’s center, appearing in Figure 3 (*e.g.*, T18/R2E) as irregular blank expanses that signify the absence of witness trees. Another 50 sq miles of oak-savanna occurred north of the more open prairies (T19&20/R2&3E). When surveyors entered open prairies or savannas they were obliged to mark corners with “a mound of stone around post as no tree was convenient.” Peripheral coniferous forests around the margins were dominated by Douglas-fir.

The pre-settlement vegetation in South Prairies differed starkly from that of both Hemlock Hts and Floodplains, even though portions of each area were only a few miles away. Precipitation in South Prairies would appear to be sufficient for trees (40-50 inches, equivalent to that of much more heavily treed areas), yet the relative absence of trees suggests that something else impeded establishment. A long history of anthropogenic burning for prairie management has been implicated (Leopold and Boyd, 1999; Peter and Harrington, 2014), but highly pervious gravelly outwash soils likely also slowed tree establishment. The paradoxical proximity of small ponds and excessively drained outwash may indicate subsoil heterogeneity: topographic depressions may retain water if underlain by lenses of impervious clay or hardpan, that could easily have resulted from glacial kettles or other post-glacial processes. Whatever the causes, early and current vegetation of South Prairies underscores that tree success does not always correlate perfectly with precipitation alone: excessive or problematic drainage can be inhospitable to tree germination and survival, especially where the fire-return interval is also short.

Fire-resistant Garry oak found in South Prairies’ savanna-like settings averaged about 20 inches in diameter, ranging as high as 54 inches. Oak was numerous toward the area’s north, becoming scattered in the more open south. A roughly equal number of similar-sized Douglas-fir co-occurred with the scattered oak; they too increased in concentration toward the north and east. The dynamics of South Prairies vegetation in the centuries and decades before GLO surveying is not fully understood. In 1839, well after a demographic decline in Native populations (Boyd, 1999), Hudson’s Bay Company established the Puget Sound Agricultural Company on South Prairies to raise livestock on preexisting grassland. Grazing by cattle and sheep must have had consequences for tree germination, but details of the extent and locations remain unclear. By the time of the GLO surveys three decades later, areas immediately outside treeless bunch grass prairies already supported dense stands of Douglas-fir, apparently representing pioneers that had begun to encroach from prairie margins after intentional burning diminished with the demographic decline.

The heart of South Prairies cannot be characterized by witness-tree data because it lacked trees. On the other hand, an examination of adjacent coniferous forests can document tree encroachment upon the grasslands; for that purpose, a database pooled witness-tree information from four forested townships in the north, east, and southeast peripheries of the prairie/savanna complex (identified in Appendix E). In those peripheral stands Douglas-fir were very abundant (65.7%) and averaged 22.1 inches in diameter, somewhat exceeding the regional average; their diameter histogram (Figure 5b) is symmetrically domed, perhaps indicative of multiple age cohorts, though less emphatically than the histogram for Rainshadow’s Douglas-fir. Extremely few hemlock were present (2.8%), and they averaged 14.3 inches in diameter, only slightly less than in Hemlock Hts. Redcedar represented 14.5% of all stems and occurred in scattered aggregations throughout the peripheral forests. In the encroached peripheral zone 10% of trees were oak.

Peter and Harrington (2014) analyzed Douglas-fir encroachment of these prairies during *post*-settlement times, noting that extensive invasion began in the 1890s, that is after the Agricultural Company had departed and after the GLO surveys. The authors also comment, more obliquely, that an earlier invasion may have germinated in the early 1700s, presumably accounting for many of the Douglas-fir witness trees of the present study.

### SW Colonizers

Pre-settlement vegetation of this 500-sq-mile oblong area in the region’s southwest exhibited an unusually uniform species composition that superficially resembles the forest composition of Rainshadow much farther north. Centered on the southern reach of Hood Canal, SW Colonizers includes both low-elevation terrain farther south and the 600-feet-high domed Tahuya Peninsula, located within the crook of the Great Bend. Douglas-fir and pine were almost the only species present, except for narrow riverside bands of hardwoods along the Skokomish River and a creek at the head of Hood Canal. A database of witness trees was collected from ten townships (∼350 sq miles) in the area’s distinctive core.

Douglas-fir, with an average diameter was 19.1 inches, accounted for 67.1% of pooled witness trees. The form of their diameter-distribution histogram in Figure 5b is symmetric (unimodal) and rather compressed, suggesting that the age range in the population was also narrow; the few larger specimens in the histogram presumably represent sparse legacies of an older cohort. “Pine” trees amounted to 6.8% of stems, averaged 8.4 inches in diameter, and occurred in small aggregations; most were probably lodgepole pine, although some on Tahuya Peninsula might have been western white pine, judging from that species’ occurrence there today. A few scattered hemlock with an average diameter of 10.8 inches also occurred, and their histogram was again unimodal and narrow (Figure 5c). There were virtually no redcedar or alder.

The extensive, quasi-monoculture of Douglas-fir in SW Colonizers may be partially attributed to its excessively drained substrate. Most soils in SW Colonizers are deep deposits of coarse gravelly outwash with very little moisture-retaining ability. Even though precipitation exceeds 70 inches in the area’s western and northern parts, declining to about 60 inches toward the east, water rapidly drains away from the surface. A recent soil-moisture map characterizes soil conditions for the entire SW Colonizers area as “aquic-xeric” (Peterman, 2012), meaning that soils may saturate in winter with static, anoxic water (implying a deeper impervious layer) and then desiccate severely in summer. (See similar evidence in Appendix A, Figures A.1b and c.) Alternating extremes of soil moisture impede germination and seedling survival, leading to delayed aforestation after a disturbance and slowed ecological succession.

As well-known colonizers, Douglas-fir and lodgepole pine are also often considered “fire followers.” The preponderance of those species in SW Colonizers may have resulted from a serious wildfire that effectively obliterated all traces of an antecedent forest stand – but there are other possibilities. Although it is conventional to refer to forestland conflagrations as “stand-replacing” events, that term contains a potentially misleading presumption: it presupposes that the pre-fire land cover was necessarily a closed forest, rather than a more open or even treeless landscape, say, like a savanna.

Given its soil conditions and proximity to Native habitation, it is possible that SW Colonizers’ long-standing prehistoric land cover was similar to South Prairies, that is, a patchwork of treeless prairie, savanna (prairie with scattered open-grown trees), and parkland intermixed with prairie and small stands of trees. Present-day evidence of relict prairie vegetation actually supports those possibilities: a) vestige patches of camas and other signature lowland prairie-associated plants are abundant even today in and around the town of Shelton at the southern margin of SW Colonizers (Stanley Graham, personal communication); and b) bear grass, a long-lived herbaceous perennial more commonly associated with wetter highland savannas, persists today at numerous locations at intermediate elevations southwest of Hood Canal’s Great Bend and on the Tahuya Peninsula (Peter and Shebitz, 2006). Thus, SW Colonizers, rather than being a continuously forested area, may in antiquity have been a phasic mosaic of transitory forest stands intermixed with fire-susceptible savannas, perhaps dominated by bear grass in wetter portions and bunch grasses where drier; such a scenario may gain plausibility by considering climatic extremes that characterized the Little Ice Age (1300 to 1870). If so, the low-diversity pine-Douglas-fir quasi-monoculture documented by GLO surveys may have established after a scorching savanna fire, not a fire that consumed a closed forest.

### Matrix Forest

Together, the five distinctive forest areas described above occupy one-third of the land area of Puget Sound country; the remaining ∼4000 sq miles of interstitial Matrix Forest presented compositional intermediates between the extremes of the specialized areas. Matrix Forest, however, is so diverse that no single part can be considered representative. Accordingly, a witness-tree database pooled from five townships near the region’s center captures a “balanced” composition in which no singular species predominated; the sampled pool is an evenly divided mix of Douglas-fir, redcedar, hemlock, and representatives of all other species (Figure 5a). Tree diameters were likewise mostly intermediate, except for a few larger legacy specimens (Figures 5b and c).

A band of Matrix Forest rich in hemlock crossed the region’s midsection (*e.g*., T27/R4E in Figure 3) from the mainland north of Lake Washington across southern Whidbey Island, northern Kitsap Peninsula, to Quimper Peninsula. These terrains are neither high in elevation nor influenced by rivers, so the source of elevated soil moisture generally required for hemlock success is not immediately evident. One possibility is that the stripe may have benefited from increased rainfall from the Puget Sound Convergence Zone, especially if that meteorological anomaly was stronger during the late Little Ice Age; in modern times the Convergence Zone causes a modest increase in local rainfall where marine airflow recombines in the lowlands after splitting around the Olympic Range.

Further south, where redcedar and hemlock were less abundant, Matrix Forest seems to unify SW Colonizers and South Prairies areas into a larger zone of Douglas-fir dominance that embraces South Sound’s array of finger inlets. The intimate proximity of pioneer vegetation and inlets may not be wholly coincidental; both may be attributed to the very loose substrate – gravelly outwash – dumped by the receding glacial terminus: a) on the plains unable to retain moisture, hence supportive only of colonizers; and b) deeply eroded at the end-phase of recession by retrograde subglacial drainage (*i.e*., northward toward Strait of Juan de Fuca) and subsequently flooded by sea-level rise post-glacial.

### Summary of Distinctive Areas

Tree parameters of specialized forest areas (omitting Floodplains) are displayed together in Figure 5 to allow comparisons. Curiously, tree diameters varied rather little between areas; principal differences were in species composition. Graphic elements in Figure 5 are arranged to provoke two synthesizing considerations: a) the first four areas (SW Colonizers to Hemlock Hts) roughly align sequentially along bio-geographic gradients; whereas, b) the uppermost area, Rainshadow, is best excluded from that scheme and considered separately, for reasons that will emerge below. The first four areas progress geographically from southwest to northeast, topographically from sea level to foothills, by precipitation from moderate to high, and successionally from pioneer (dominated by shade intolerants) to advanced (shade tolerants). It will be shown that Rainshadow was crucially different from mainland areas in pre-settlement times, despite its superficial compositional resemblance to SW Colonizers, largely because growing conditions there were so much poorer.

## Exploratory Exercises

### Very Large Trees

Popular literature and imagery sometimes imply that trees in the early forests around Puget Sound were uniformly immense in girth and height. Frequent claims that Douglas-fir, western redcedar and western hemlock, as species, can attain very large size and great longevity are often misconstrued as evidence that pre-settlement forests were primarily populated by gigantic, ancient trees. With such over-stimulated expectations, it must be difficult to accept that GLO surveyors found the average tree during pre-settlement times to be merely 16.9 inches in diameter (or 18.5 inches, if hardwoods are excluded). Indeed, the majority of trees were smaller than these averages, as illustrated by diameter histograms, even if a few were very large.

Of the 49,746 witness trees for this study, 1,940 (3.9%) were 4 feet or more across – larger than two people with linked arms can embrace. One-third of those very large trees was at least 5 feet across. Across Puget Sound country as a whole, more than half of very large trees in pre-settlement times were Douglas-fir and most of the remainder were redcedar (Figure 6, right column).

**Figure 6.**
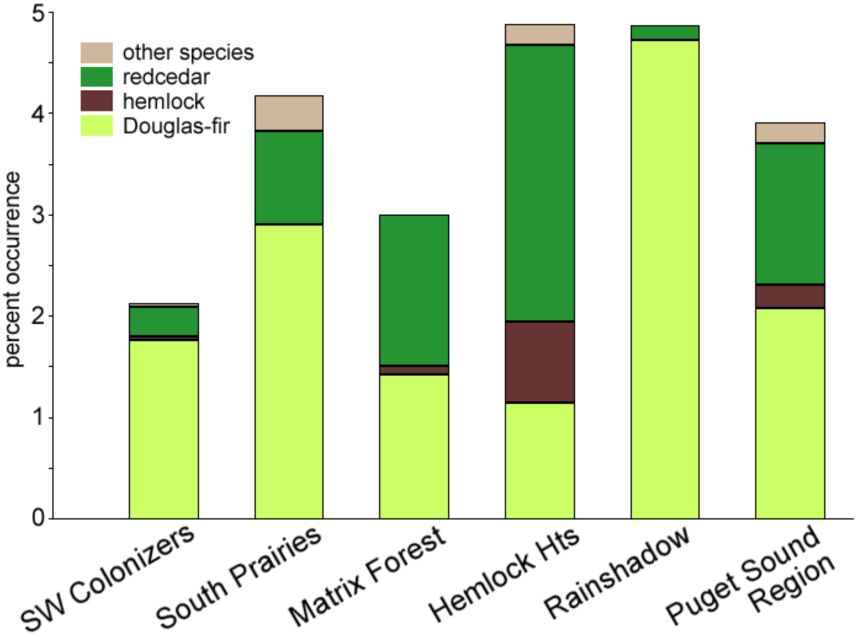
Species proportions of very large trees (≥48 inches in diameter) in area databases.

A proportion of 3.9% means that, on average, one out of every 26 pre-settlement trees was at least 4 feet across. At that rate, if evenly distributed, at least one very large tree would have been visible from every vantage point in the region; at a postulated tree density of 100 stems per acre, 26 trees would fit into a circle with a radius of 60 feet, thus all would be visible (at a density of 200, the radius would be 42 feet). Of course, the distribution of very large trees was not uniform. Figure 6 indicates that very large trees were more common in Hemlock Hts and Rainshadow areas (one of every 21 trees, mostly redcedar in the former and Douglas-fir in the latter) and least common in SW Colonizers (one in 49 trees, mostly Douglas-fir). Interestingly, of the twelve townships whose data were pooled for the Hemlock Hts database, nearly half of the area’s very large trees were clustered in only three townships around T24/R8E in the southern Mt Baker-Snoqualmie National Forest due east of Lakes Washington and Sammamish.

Beyond tree size, another recurring question about early forests is: How many trees were there? If, as previously postulated, the average tree density in the original forests around Puget Sound was either 100 or 200 trees per acre, then the entire region (6,300 sq miles) would have had 0.4 or 0.8 billion trees, respectively (64,000 or 128,000 per sq mile). Reflecting back to the 3.9% abundance of very large trees, the region-wide total would therefore have been 15,600 or 31,200, or, on average, about 2,500 or 5,000 very large trees for every square mile of forest.

### Successional Status

In broad terms, the process of ecological succession in the Western Hemlock Zone commences, after a stand-replacing disturbance, with newly exposed land being invaded by drought-tolerant, shade-intolerant colonizers such as Douglas-fir, red alder, and sometimes lodgepole pine, depending on seed availability. After 50-100 years of pioneer growth, increased shade on the forest floor discourages further germination by colonizers, whereas accumulated litter improves moisture retention enough that shade-tolerant redcedar and hemlock begin to appear. With further aging, attrition gradually reduces the number of original pioneers; short-lived alder and pine are the first to die off, whereas attrition among longer-lived Douglas-fir may continue for centuries. Meanwhile, shade-tolerant species progressively gain prominence and eventually ascend into the canopy. The transition into late-successional status, commonly known as old growth, occurs after 150 to 200 years when shade-tolerant species become dominant in the canopy. Ultimately, ecologists characterize old growth by a suite of structural and functional attributes, not just “time served”; typical diagnostic features include canopy layering, distributions of tree ages and sizes, species representation, decadence, quantities of down logs or snags, etc. Experienced foresters can recognize definitive old growth by these features, but *threshold criteria* for the transition from mature growth into old growth are more elusive, and ultimately arbitrary.

In the 1980s, in response to concerns about dwindling quantities of old-growth stands in Pacific Northwest forests, a team of scientists and managers enunciated entry-level or minimum threshold criteria that a stand must meet to qualify as old growth. The Old Growth Definition Task Group (OG Task Group, 1987) specified threshold parameters for old growth in the Western Hemlock Zone based, in part, on the metrics of large-tree abundance. Specifically, their “Interim Definition” states that a forest stand would be considered old growth if – on a per-acre basis – it contains: a) at least eight Douglas-fir specimens 32 inches or more in diameter; and/or b) at least 12 redcedar or hemlock specimens at least 16 inches in diameter. The Interim Definition also noted that if either component (a) or (b) is poorly represented in a stand, regardless of tree size, it could be sub-typed according to its predominant component, namely as “Douglas-fir old growth” or “hemlock old growth.”

The present exercise will gauge township stands of pre-settlement forests against the Interim Definition’s threshold criteria. To do so, however, a means must first be invented to harmonize the absolute numbers of large trees per acre (Interim Definition’s metric) with relative frequencies (percentage proportions among GLO witness-tree populations). An absolute-to-relative conversion requires that the two systems share a common tree density (trees per acre) for which, regrettably, there is not better recourse than reasonable guesswork. Over a large area of unmanaged forest, tree density realistically ranges between 50 and 300 trees per acre; for the purposes of this exercise, an across-the-board stand density of 100 trees per acre is assumed as a point of reference. This arbitrary value now allows the Interim Definition’s absolute tree numbers to be relativized and thus offered as a yardstick against which witness-tree datasets can be compared. For example, in a stand with 100 trees per acre, the requirement of eight Douglas-fir ≥32 inches in diameter can also be expressed as 8%; therefore, upon screening a township’s witness trees, if at least 8% of its trees were Douglas-fir of the requisite diameter, or similarly if 12% were shade-tolerant redcedar or hemlock of sufficient size, then the township forest passes the qualifying test as old growth.

Witness-tree databases for all townships were filtered according to the Interim Definition’s criteria, and the results are illustrated in Figure 7. Some townships failed the test entirely, mostly south of Lake Washington. On the other hand, about three-fourths of the land area around Puget Sound did qualify as old growth in one of three subtypes: Douglas-fir old growth, where only that species passed; hemlock old growth, where hemlock and/or redcedar passed but the Douglas-fir component failed; and hemlock + Douglas-fir old growth, where both components met or exceeded Interim Definition quotas.

**Figure 7.**
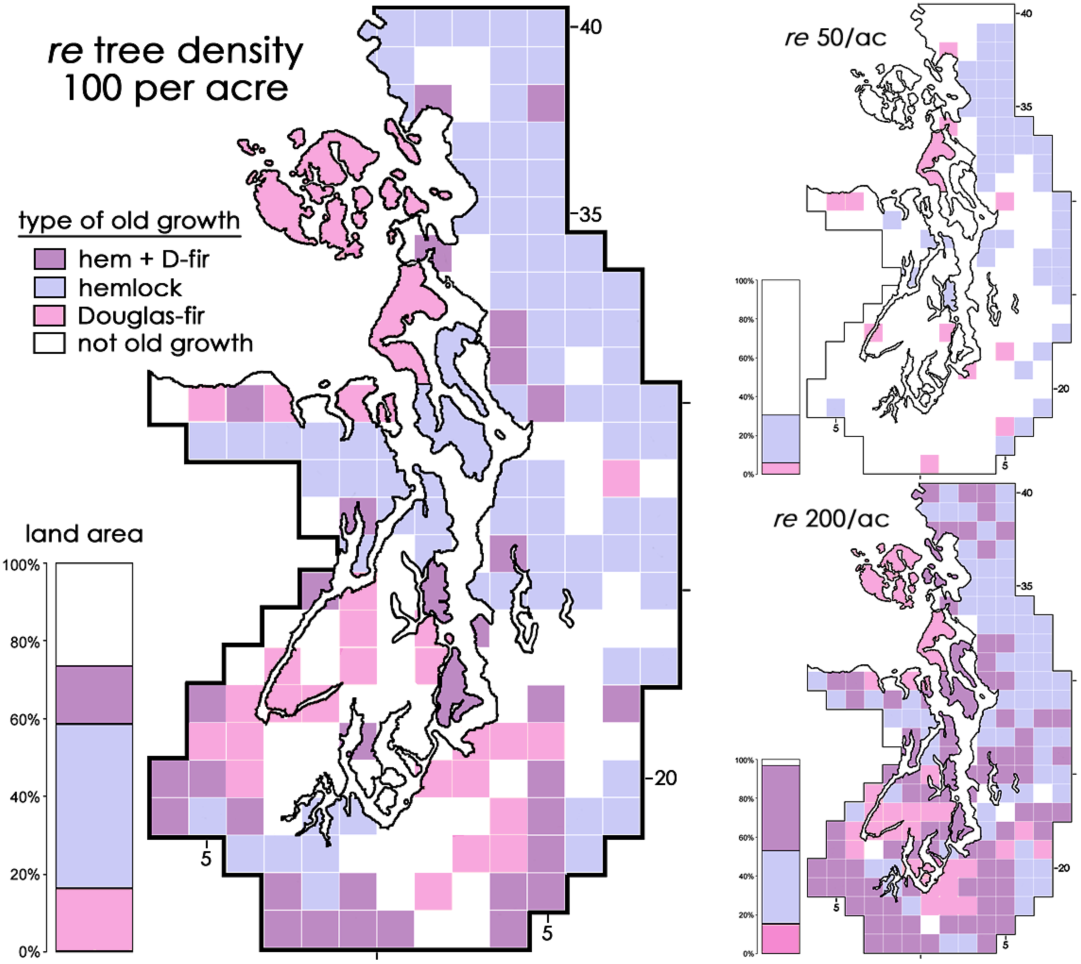
Successional status by township according to threshold abundances of large trees specified by Interim Definition, calculated at three assumed forest densities. Types of old growth based on species of qualifying large trees: “hemlock” implies an admixture with redcedar; hem + D-fir signifies that both Douglas-fir and hemlock/redcedar met their thresholds.

Figure 7 also illustrates the problematic consequences of assumed forest-density values. By choosing a density of 50, instead of 100, the amount of old growth drops to one-third of the region’s land area. By raising the density to 200, the entry-bar is lowered so that nearly all township forests qualify as old growth. Regardless of selected density, certain areas were sufficiently robust in their old-growth status to retain the same classification, notably: many townships in the Cascade foothills and Chuckanuts remain hemlock old growth; and some parts of Rainshadow, SW Colonizers, and South Prairies remain Douglas-fir old growth regardless of density assumptions.

Hemlock old growth was generally more widespread in the region’s northern half than south of Lake Washington. Conversely, despite having more land area whose forests failed to qualify as old growth altogether, the southern half displayed more Douglas-fir old growth or hemlock + Douglas-fir old growth than hemlock old growth. Old-growth designations for the region’s major islands (San Juans, Whidbey, Camano, Bainbridge and Vashon) seemed more robustly independent of density than near-shore mainland townships; perhaps intervals between *severe* disturbances were longer on islands.

In the end, the Interim Definition provides a test of oldgrowthness for stands defined by GLO witness trees, but the results are not definitive. An alternative approach, also imperfect, is offered in Appendix F. A preferred methodology would have to account for many variables not yet considered, including site quality, locally tailored forest-density values, climatic conditions during and since the Little Ice Age, and whether a township’s set of witness trees is a fair unit for evaluation.

### Estimated Stand Ages

Empirical studies in the 20^th^ century determined that, in a stand of normal density on a site of known quality, trees of a species grow at a predictable rate. When graphed, the growth rate traces a characteristic curve, rising steeply in youth and gradually approaching a plateau with age. Forest scientists have developed competing growth models (“base 50” versus “base 100”) from which families of growth tables and growth curves have been formulated. An individual curve in a family is specific to a species and a site-quality rating – its site index – which is expressed as a size measurement of an average tree at the target age (either 50 or 100 years, depending on the model). Thus, a growth curve identified as SI 110 (Douglas-fir; base 50) will show that at 50 years of age (either above breast height or total, according to the way the model was composed) the height of the average Douglas-fir canopy tree will be 110 feet; some versions also show that the tree would be about 13 inches in diameter. For a different species or a different site quality some other growth curve will apply.

The present exercise estimates ages of pre-settlement stands in townships and, separately, of specialized forest areas by reverse-reading empirical growth models. That is, once the site quality is known, the size of the average witness tree at that site will indicate the age of the pre-settlement stand at the time it was surveyed. Of course, GLO information includes neither local site indexes (another 20^th^-century construction) nor tree height; rather, the survey record provides only tree diameter and location; thus, to estimate stand ages from witness trees it will be necessary to consult growth tables and curves that employ *diameter* values, and equally necessary to discover site-index information from an outside source that can reasonably be applied to townships and groups of townships. Tools that satisfy these requirements and that will be used for this exercise are reproduced in Appendix G, which also elaborates how they were employed to estimate stand ages.

Stand ages were estimated at two geographical scales: individual townships (for Douglas-fir only) and distinctive areas (for Douglas-fir and hemlock separately), the latter based on tree data pooled from the representative townships listed in Appendix E. Results for townships are illustrated in Appendix G.2, and forest ages of special areas are tabulated in Table 1, which also documents the key measures employed in the process: diameters of average trees and site-quality ratings. The diameter metric for tree growth deserves special explanation: growth curves rely on so-called quadratic mean diameters (QMDs), rather than the more conventional linear average diameters (arithmetic means) featured so far in this report. QMD is defined as the diameter equivalent of the average basal area of a group of trees – basal area being the group’s aggregate cross-sectional area at chest height. Tree growth is a volumetric process, so QMDs offer closer proxies to volumetric increase than one-dimensional average diameters because derivations of QMDs include a square function (Curtis and Marshall, 2000). As illustrated in Table 1, the QMD of a group of trees is always somewhat larger than its linear mean because QMD weights larger-diameter specimens over small ones.

**Table 1.**
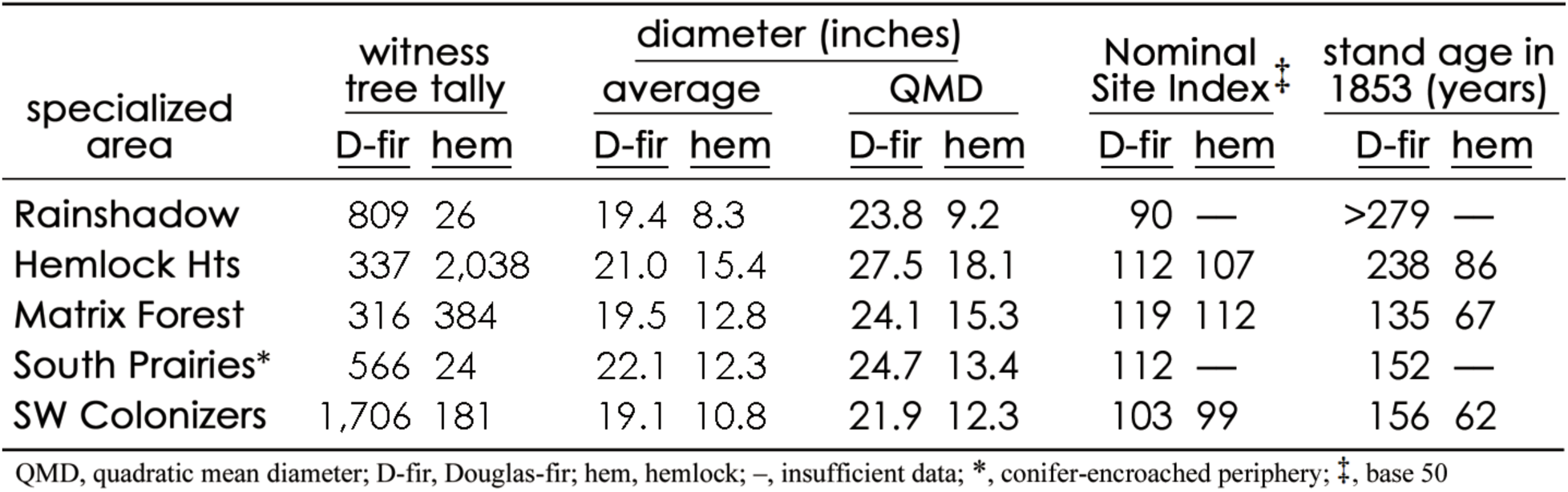
Stand ages, normalized to 1853, as derived from witness-tree size and site quality

Site quality is expressed by a numeric site index that is linked to a particular empirical growth model, here always base 50; the index number declares the average tree height (convertible with QMD in the chosen models) that a stand will attain after 50 years. A site index generally applies to a small polygon of landscape, perhaps only an acre in area, as in a soil survey; a so-called *site class* is a larger polygon (several square miles) that aggregates neighboring site-index polygons with a range of site-index values. For this study, the midpoint of the range of site indexes is taken as the Nominal Site Index for a site-class polygon. Appendix G.1(a) reproduces a published base-50, Douglas-fir-specific Site Class Webmap for Puget Sound country, customized to include Nominal Site Indexes. As the map shows, a township generally overlaps more than one site-class polygon, so its singular Nominal Site Index is derived by area-weight averaging its site classes. Nominal Site Indexes for mainland locations ranged between 103 and 119; for Rainshadow, the index was decidedly lower: 90.

Douglas-fir stand ages *for townships* (see Appendix G.2) mostly fell into 50-149 or over-250 year classes, the latter probably indicating the presence of ancient legacy trees. Douglas-fir and hemlock ages for distinctive areas are shown in Table 1, above; they were normalized to 1853, the year that GLO surveys in the region began. Stand ages varied according to species and areas. Douglas-fir stands in the lowland areas of SW Colonizers, South Prairies, and Matrix Forest, where that species was common or predominant, ranged from 135 to 156 years, indicating that they germinated between the years 1697 and 1718. Hemlock populations in the same areas were several decades younger (62 to 67 years of age, therefore germinating between 1786 and 1791). Thus, ages of the two species differed by about 80 years.

In Hemlock Hts both Douglas-fir and hemlock components were older than their counterparts in other mainland areas, but they differed from each other by 150 years, longer than expected for normal successional processes. The diminutive population of Douglas-fir (one-sixth the abundance of hemlock) contained a number of large, putatively legacy specimens, consistent with its stand age of 238 years (germinated on average in 1615); the predominant hemlock population, however, at 86 years of age (germinated in 1767), was only slightly older than hemlock stands in the lowlands.

The impression gained from these age estimations of Puget Sound’s *mainland* areas is that the successsional clock of their forests was reset around the year 1700, implicating a catastrophic disturbance that spared only a scattered minority of legacy Douglas-fir specimens from earlier cohorts. If ecological succession was restarted around 1700, young Douglas-fir colonized most parts of the region without much competition for the first 80 years, or so, at which time hemlock (and redcedar) appeared. Events in Hemlock Hts, however, evidently took a somewhat different course in which hemlock reestablishment somehow occurred earlier and without much competition from Douglas-fir.

Stands of Douglas-fir in Rainshadow proved to be far older than stands on the mainland, perhaps even older than many legacy Douglas-fir in the foothills. The estimated age, however, was indeterminate, that is literally “off the charts”: definitely older than 300 years because the QMD in 1874 fails to intersect the relevant growth curve. One may reasonably conclude, therefore, that Rainshadow stands were unaffected by the suspected catastrophic event that reset the successional clock for most mainland forests. All of these aspects of Rainshadow’s forest are what differentiates it so dramatically from the superficially similar growth in SW Colonizers and that caused it to be displaced in Figure 5, outside of the developmental continuum proposed for its mainland counterparts. Conceivably, *insularity* itself protected Rainshadow from the clock-resetting event; in this regard, it is noteworthy that Douglas-fir components of *all* major islands in Puget Sound were older than 250 years, even islands not part of Rainshadow and with higher Nominal Site Indexes (Appendix Figure G.2).

### Destructive Disturbances

The fourteen thousand pages of GLO field notes that underpin this study contain observations made along 12,600 lineal miles of section lines; the records include 1,108 instances of recognized forest disturbance – about one comment per 10 miles surveyed: 781 citations for fires of some kind (usually only tersely cited as “fire,” “burnt,” or “dead”) and 337 citations indicating windthrow (“fallen”). Most comments simply announced and fail to elaborate on possible causation, timeframe, or spatial extent of the disturbance.

Of all reports of windthrow two-thirds are clustered in the region’s south-central Cascade foothills (T22-25/R7-8E in Figure 9, the same area that was most heavily endowed with very large trees); not coincidentally, that area is directly upslope from the Enumclaw plateau, a Native-American place-name that refers to strong winds. In one township (T25/8E) 90% of section-line summaries conclude with the phrase “heavy fallen timber,” almost always in association with thorny devil’s club in the underbrush; curiously, a different surveyor working in the adjacent township to the north scarcely mentioned down logs, yet even further north, in T27/8e in the year 1883, the township’s final summary again declares “devil’s club, which has grown up among the fallen timber, making it very difficult to get through … timber is nearly all blown down … immense windfall.” Outside of this particular portion of the Cascade foothills, references to windthrow were incidental and unremarkable.

Disturbance by fire was more frequently reported, yet only one comment referred to a fire actually in progress: “fire now burning” (T32/R4E in 1872). Some remarks were especially evocative: “burnt over in the last 20 or 30 years” (T40/R5E); “not one tree in twenty living” (T39/1E); “timber … killed by fire, and has since fallen, covering the ground … to a depth of 10 to 15 feet” and “a fire five years ago destroyed most of the timber” (T21/R6E). Survey notes for Nooksack River’s flood plain were especially graphic (and ironic, considering the area frequently flooded): “the soil is greatly deteriorated by the fires, which have burned the alluvial soil in places to the depth of one foot” (T39/R2E) and “the timber is about all killed by fires, there not being more than a section in the township unburned” (T40/R4E in 1874).

Many fire references are ambiguous or even contradictory. For example, the very common entry “timber all killed by fire” is often followed, without explanation, by nearby living witness trees, thereby ostensibly negating the assertion that *all* trees were dead. Unambiguous references to the extent of fire destruction are rare, but in one location a linked pair of citations stands out as exceptionally convincing: the 1856 survey for T23/R2E explicitly declared that, due to fire, witness trees were unavailable for several square miles on either side of a marine channel (evident in Figure 3 as treeless blanks on Vashon Island and on Kitsap Peninsula to its immediate west, separated by Colvos Passage). The geographic proximity and parallel language of those entries suggest that a single fire may have crossed 0.9 miles of open water!

Figure 8 illustrates the townships in Puget Sound country that exhibited the highest instances of fire citations (exceeding 20 per 36-sq-mile township). The southern lowlands and San Juan County received few fire citations, despite traditions of anthropogenic burning for both locations. Conversely, fire evidence was abundant in Island County and three townships in or around Nooksack River’s large flood plain; both areas were heavily trafficked by Native people.

**Figure 8.**
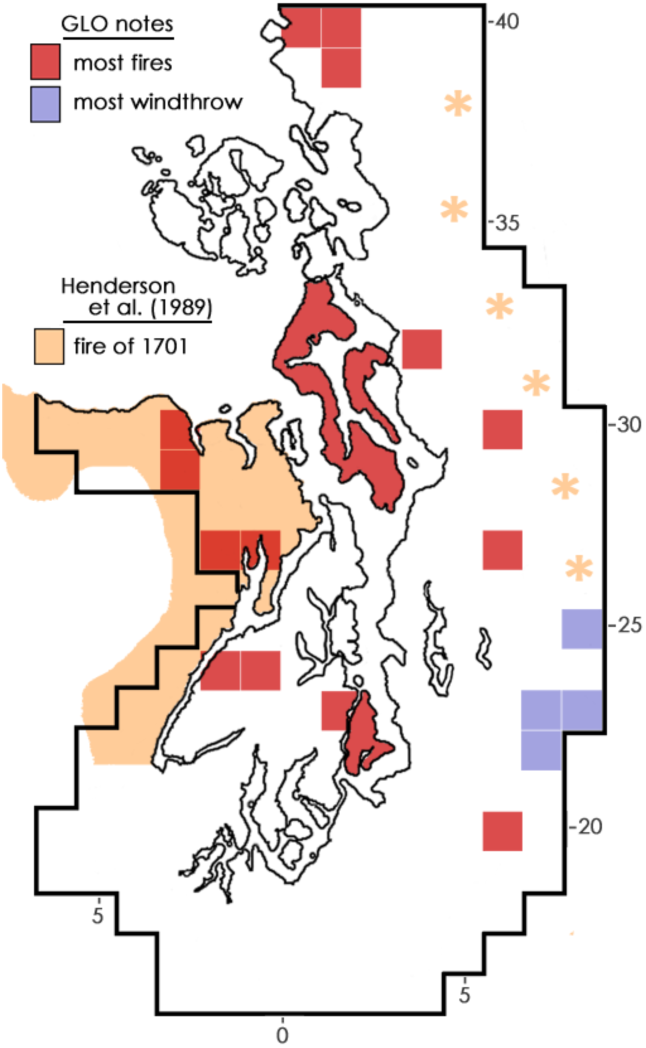
Forest disturbances. Heaviest concentrations of windthrow (blue) and fire (red), and published extent of prehistoric fire (orange; asterisks, unpublished evidence of same in Cascades).

As mentioned in the context of ecological succession, forests can be reconfigured by destructive disturbances and subsequent recovery processes. By themselves, GLO survey records proved frustratingly vague and unenlightening about the role that wildfire may have played in shaping early forests. On the other hand, modern investigations may have discovered evidence that is dramatically more pertinent to the late pre-settlement landscape.

Recent research provides evidence that a great fire (or a family of roughly contemporaneous fires) may have passed through the entirety of Puget Sound country in the year 1701. If correct, such a large-scale destructive event could explain much about the age structure and successional status of the region’s diverse forests at the time of the GLO surveys. In a monumental ecological study of the Olympic National Forest, Henderson *et al*. (1989) discovered abundant and widespread fire scars on trunks and stumps that all dated to 1701. They concluded that a great fire burned more than 1,500 sq miles of forest along the northern and eastern sides of Olympic Peninsula, including all lands west and north of Hood Canal’s Great Bend in the pattern tinted orange in Figure 8. Subsequent paleo-environmental and palynological studies by Gavin *et al*. (2013) concur with that interpretation. Both published studies, however, were jurisdictionally confined to the bounds of federal National Forest lands, which explains why the burn pattern in Figure 8 ends so abruptly at its southern extreme. Circumstantially, the 1701 fire almost certainly extended into the present study’s SW Colonizers area. The original investigators further imply that the fire may have affected most of the region, even crossing 60 miles of lowlands into (or out from) the Cascade Range.

Henderson *et al*. (1989, p. 33) summarize the effects of the great fire of 1701 rather obliquely, yet provocatively: “This fire or series of fires apparently burned more than one million acres on the Olympic Peninsula, and 3 to 10 million acres in western Washington. Much of the valuable Douglas-fir old-growth, that has formed the basis for the local timber industry, is the result of this great fire.” As explained by co-author David Peter (US Forest Service, personal communication), this statement alludes to additional widespread fire-scar evidence, again dated to 1701, from Mt. Baker-Snoqualmie National Forest in the Cascades, collected by the same team but not yet published; those observations are suggested in Figure 8 by orange-colored asterisks.

If two large fires were underway simultaneously on both sides of Puget Sound, then they possibly were united as a single, immense burn that continued right across the region; that is understood to be the significance of the aforementioned “3 to 10 million acres” (or 4,700 to 15,600 sq miles). In their analyses of Olympic Peninsula’s fire history, both Henderson *et al*. (1989) and Gavin *et al*. (2013) speculate that increased climatic variability during the 500-year-long Little Ice Age could have introduced extended episodes of severe regional drought, high-wind events, and serious winter tree damage, all conditions that would have increased fuels, ignition susceptibility, and the spread of a great fire across the land divide between the Olympics and Cascades. Unfortunately, the continuity of a single great fire encompassing both the Olympics and Cascades cannot be confirmed by fire-scar data from Puget lowlands, because that area was too thoroughly altered by timber extraction, stump removal, and land conversion.

## Conclusions

The present study provides evidence that late pre-settlement forests throughout the southern half of Puget Sound country were dominated by colonizing pioneers that were mostly about 150 years old and not quite old growth, unlike slightly younger, but successionally more advanced, hemlock-dominated forests of the foothills (and presumably continuing further inland beyond the study area). Much of the pattern of forest development around low-lying Puget Sound mainland is conceivably explained by a destructive great fire around 1701 that reset their successional clocks before being extinguished in the moister foothills. Major islands in Puget Sound may have escaped the great fire, including insular forests in the Olympic Rain Shadow that were able to achieve “pioneer old growth” status.

In their foundational monograph on forests of the Pacific Northwest, Franklin and Dyrness (1979) sweepingly included Puget Sound country into the expansive Western Hemlock Zone, where the trajectory of ecological succession trends toward hemlock dominance. The broad scope of their proposal may have suited the vast, still-intact forests above 1000 feet of elevation, at a time when lowland forests were no longer available to pose persuasive, contradictory challenges. However, the present study shows that hemlock played a distinctly minor role in forests of all ages at elevations below 1000 feet – certainly in late pre-settlement times and probably also for preceding centuries and millennia.

This study identifies such distinct qualities of early forests in the Puget lowlands that it may be appropriate to dissociate them from the Western Hemlock Zone, whose categorical expectations for the role of hemlock do not seem to apply. Arguably, it seems more informative to acknowledge that ecological succession of many lowland forests persistently and predictably concluded with terminal dominance of pioneer species, thanks to convergent effects of three undeniable lowland realities: poorly developed glacial soils, especially outwash; the austerities of growth in the Olympic Rain Shadow; and millennia of Native occupation and land management. Indeed, Franklin and Dyrness (1979, p. 89) anticipated precisely such a dissociation: “eventually, the Puget lowlands may be recognized as a separate vegetative zone.” Present evidence suggests that such a separation is now warranted.

## Appendices

### A. GLO Surveys

For Puget Sound country, surveys progressed geometrically from a monument in northern Oregon. They created a grid of perpendicular lines one mile apart running in cardinal directions that enclosed squares of land (sections), each consisting of up to 640 acres. A township’s 36 sections were numbered serially from the its northeast corner in a boustrophedon ending at its southeast corner.

**Figure A.1.**
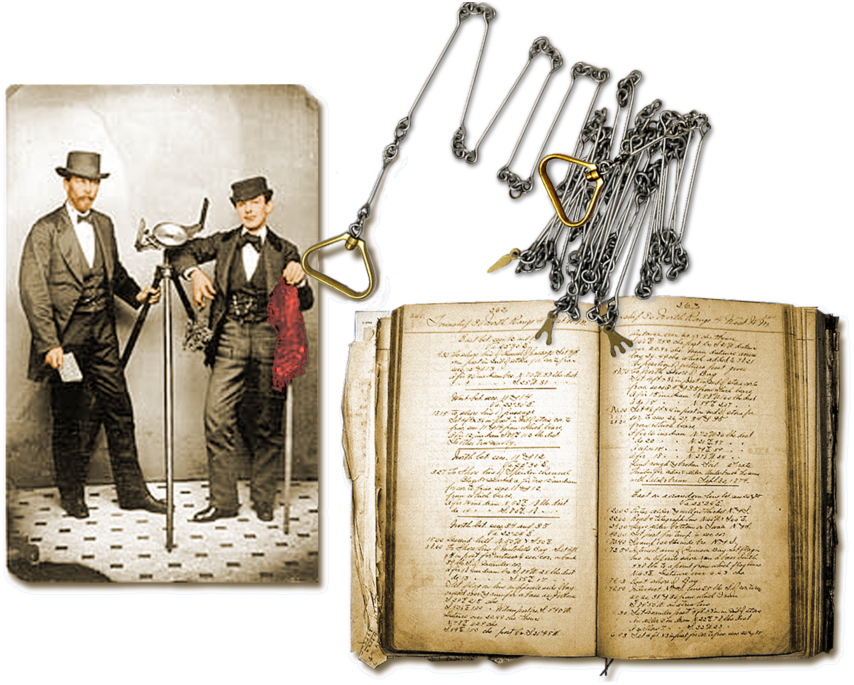
Studio photo of a GLO surveyor, with solar compass, and a flagman, 1871, colorized by the author. Steel “half-chain” 33 feet in length with 50 links. Bound field notes for a county in Puget Sound were found abandoned in a decommissioned public-works shed.

Surveyors with solar compasses and measuring chains (Figure A.1) defined section lines on the ground by installing wooden posts at section corners and points midway between (so-called quarter section corners). Witness trees, also called bearing trees, and landscape features along section lines were recorded in field notes in a prescribed, formulaic manner. Data and notes were bundled by township, copied, and bound into volumes to be used by county and state agencies. Although several subsets of witness trees appear in the surveys, only corner and quarter-corner trees were employed for this study; rejected subsets included: non-corner or line trees, which favored larger specimens; and fractional corner trees at shorelines or other impassable barriers, which represented edges but not interior forests. This study relied only on initial surveys, not resurveys. For additional details of the U.S. Public Land Survey System see Bourdo (1956), Galatowitsch (1990), Hutchison (1988), White (1983), and Whitney and DeCant (2001).

### B. Geographic Resources

The interplay of weather, soils, and topography drives forest growth in Puget Sound country. Treatises by Kruckeberg (1995), Van Pelt (2007), and Maas (2008) bring important environmental factors into focus. Figure B.1 illustrates some of these influences in map format, against which descriptions of the early forests can be assessed. The quantitative and seasonal pattern of soil moisture is most significant; wherever moisture demand exceeds moisture availability, plant growth will slow or cease altogether. Soil moisture also affects fire susceptibility, both the likelihood of ignition as well as the geographic pattern and intensity of a burn.

**Figure B.1.**
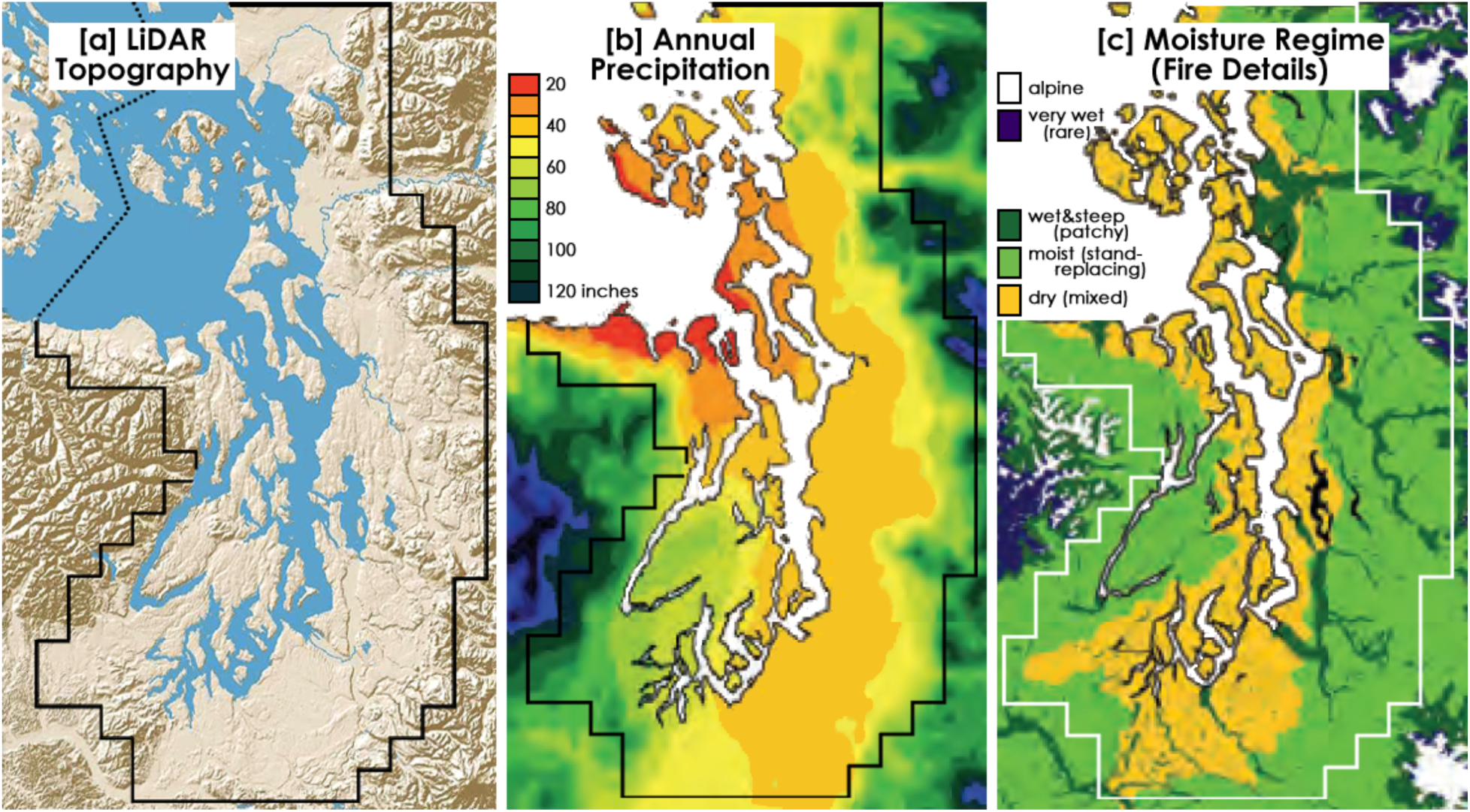
Geo-climatic circumstances, which exert defining influences over local forests, vary in complex ways around Puget Sound. Low-elevation Puget lowlands (a) cover most of the region and their axis roughly coincides with the Olympic Rain Shadow, whose locus of lowest precipitation is in the north. Droughty soils occur throughout the rain shadow, but excessively drained outwash in the south exacerbates low moisture (c); major rivers counteract climatic droughtiness along their flood plains. Figures (a) and (b) taken from Van Pelt (2007), with slight alteration.

Weather in western Washington is driven by the Aleutian Low that seasonally migrates up and down the west coast. Humid air from the Pacific Ocean brings heavy precipitation to the western slopes of the Olympic Range. Precipitation then declines east of the Olympic crest to create a rain shadow throughout the Puget lowlands, most profoundly in the north. As airflow continues up the windward slopes of the Cascades precipitation is again stimulated. If onshore winds slacken, an air inversion can occur; reversal of airflow can admit arctic air and damaging ice storms in winter, or, in summer, allow hot, dry air from eastern Washington to pour into the basin. The endemic summer drought predictably interrupts vegetation growth on a seasonal basis, but a protracted incursion of hot continental air into the lowlands can profoundly exacerbate fire susceptibility.

### C. Summary of Witness Trees

All species of witness trees included in this study are listed in Table C.1, but some names may require clarification: “spruce” means Sitka spruce; “yew” Pacific yew; “oak” Oregon white or Garry oak; “ash” Oregon ash; etc. Some common names express dual or multiple identities: “pine” meant either lodgepole pine or western white pine, depending upon situation; “white fir” meant any true fir and mostly grand fir; “juniper” could have been either common or maritime juniper; “cherry” and “willow” could have been any of a few possible species. “Schittemwood” and “laurel” are understood to mean cascara and madrone (but when mentioned in the underbrush layer “laurel” probably signified rhododendron). “Larch,” cited in a few places in the Cascades, was probably an outright misidentification. Some negligibly minor species, such as “smokewood” and “pigeonwood,” remain unidentified.

**Table C.1.** corner and quqrter-corner witness trees.

| WT species | number | rel freq.(%) | av diam.(in) |  |  |  |  |
| --- | --- | --- | --- | --- | --- | --- | --- |
| 1 Douglas-fir | 17660 | 35.50 | 19.7 | 17 birch | 166 | 0.33 | 7.3 |
| 2 hemlock | 11579 | 23.28 | 14.4 | 18 dogwood | 150 | 0.30 | 6.9 |
| 3 redcedar | 8069 | 16.22 | 22.3 | 19 barberry | 125 | 0.25 | 6.3 |
| 4 alder | 3982 | 8.00 | 9 | 20 madrone | 116 | 0.23 | 8.6 |
| 5 bl maple | 1652 | 3.32 | 12 | 21 yew | 83 | 0.17 | 9 |
| 6 vine maple | 1300 | 2.61 | 4.7 | 22 cascara | 69 | 0.14 | 5.3 |
| 7 willow | 1105 | 2.22 | 5.6 | 23 hazel | 60 | 0.12 | 6.7 |
| 8 spruce | 707 | 1.42 | 20.4 | 24 juniper | 18 | 0.04 | 10.5 |
| 9 pine | 676 | 1.36 | 9.5 | 25 aspen | 15 | 0.03 | 5.4 |
| 10 true fir | 428 | 0.86 | 17.1 | 26 skunkwood | 13 | 0.03 | 8.9 |
| 11 oak | 385 | 0.77 | 20.7 | 27 thornwood | 11 | 0.02 | 7.9 |
| 12 cottonwood | 310 | 0.62 | 17.1 | 28 elder | 6 | 0.01 | 4.3 |
| 13 ash | 303 | 0.61 | 11.4 | 29 pigeonwood | 3 | 0.01 | 5.3 |
| 14 crabapple | 279 | 0.56 | 6.8 | 30 arrowwood | 1 | 0.00 | 4 |
| 15 cherry | 259 | 0.52 | 6.6 | 31 serviceberry | 1 | 0.00 | 12 |
| 16 larch (misID?) | 213 | 0.43 | 20.2 | 32 smokewood | 1 | 0.00 | 6 |
|  |  |  |  | 33 sweetwood | 1 | 0.00 | 3 |
|  |  |  |  | **All WT =** | **49,746** | **100.00** | **16.5** |

### D. Database and Map Construction

Spreadsheets of raw data were assembled in Microsoft Excel while referring to downloaded images of handwritten field-note pages. Individual witness trees were encoded according to species, diameter, section and township, location within a section, and originating page number. Preliminary analyses of township databases were secondarily rearranged and filtered for analytical explorations. Data-reduction summaries were collated as “township digests” (two examples in Figure D.1). All 200-plus digests may be viewed by opening *supplement: Township Digests* on this article’s bioRxiv webpage by activating *Supplementary Material* beneath the “Download PDF” button.

**Figure D.1.**
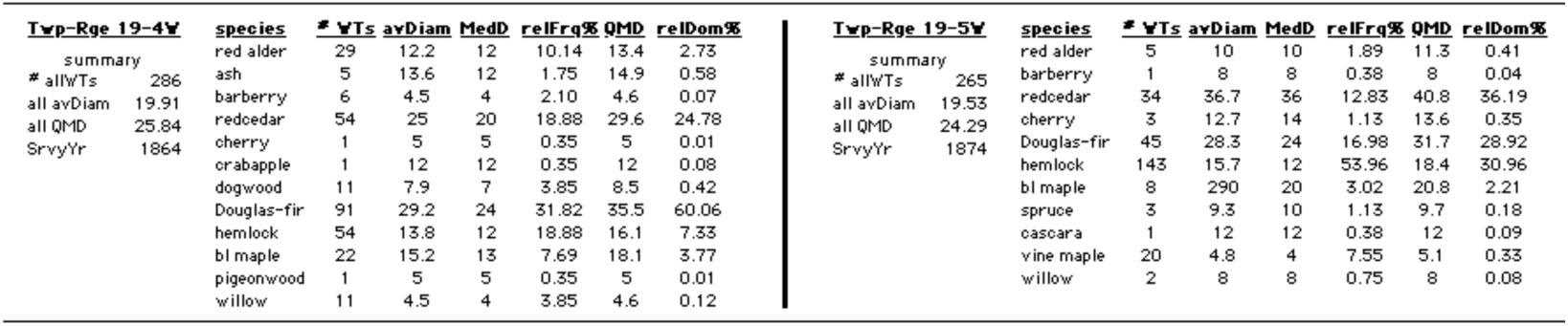
Samples of township data digests. All digests are available in an accompanying supplement.

While consulting the primary databases, the witness-tree map (text Figure 3) was assembled, one township at a time, in graphic layers in Adobe Photoshop 5.0, as schematized in Figure D.2. Composition of a generic township started with a removable template grid as a graphic layer; the template consisted of 324 cells in an 18×18 array to accommodate a township’s 288 color-coded witness trees and its 36 section numbers. Nine cells comprised each section, as illustrated in the inset of text Figure 3; the section’s number occupied the central cell and color-coded witness trees occupied the surrounding eight cells: four for corner trees and four for quarter-corner trees. Coding colors were added manually to template cells, as directed by location information in database spreadsheets. Wherever a witness tree was missing in the record, its cell was left unfilled (blank). When a township was complete, the template grid was removed and squares were fitted seamlessly together. To complete the map of the Puget Sound region, adjacent colorized townships were tiled together according to their T/R coordinates. A standard sketch of the Puget Sound’s shorelines and rivers was superimposed manually upon the array of townships by “best fit.”

This graphic methodology fills 89% of landscape space with colors (except for section numbers) that represent species locations and distributions in the pre-settlement forests. These benefits are achieved at the cost of grossly inflating the spatial scale of individual witness trees to a real-world equivalent of one-ninth of a square mile, or 1760 feet on a side.

**Figure D.2.**
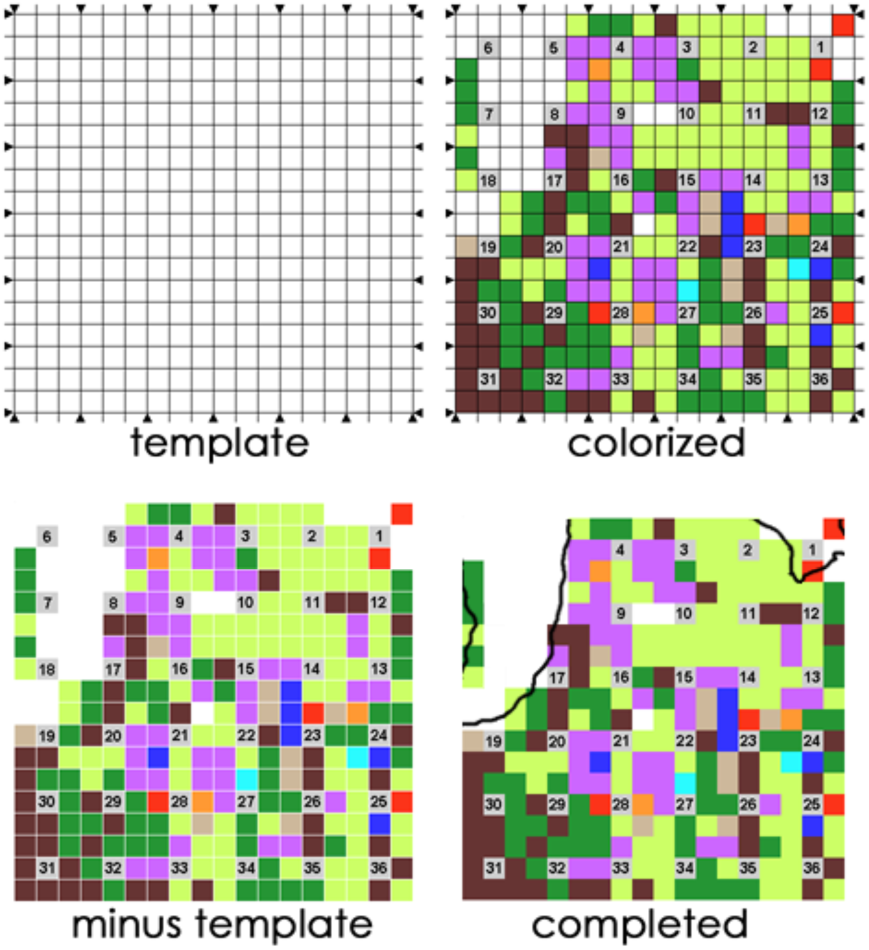
Schematized production steps of a township (T29/R1W) of the species map.

### E. Representative Townships

Specialized forest areas around Puget Sound were characterized by pooling data from representative townships, whose locations appear in Figure E.1 and whose T/R identities are itemized below:

- **Floodplains** (F): T20/4E, T39/2E, and T39/3E (pool of 836 witness trees)
- **Rainshadow** (R): San Juan County (175 sq miles in land area, 1,378 witness trees)
- **Hemlock Hts** (H): T18/6E, T18/4W, T19/6E, T23/7E, T25/8E, T26/8E, T29/7E, T30/7E, T33/5E, T33/6E, T36/4E, and T37/4E (3,362 witness trees)
- **South Prairies** (SP): T17/2E, T18/4E, T19/3E, and T20/2E (862 witness trees)
- **SW Colonizers** (SW): T20/3W, T21/3W, T21/2W, T22/2W, T22/3W, T22/1W, T22/1E, T23/2W, T23/1W, and T23/1E (2,544 witness trees)
- **Matrix Forest** (M): T24/5E, T27/2E, T27/5E, T28/5E, and T30/5E (1,266 witness trees).

**Figure E.1.**
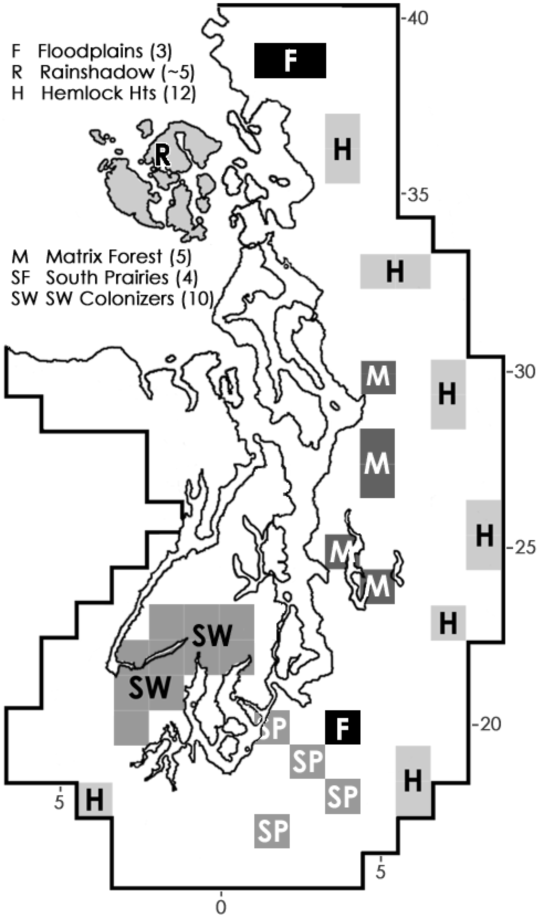
Representative townships (and tallies) for data pooling.

### F. Alternate Old-Growth Estimation

The Interim Definition distinguishes old growth from younger forests by proposing a threshold or pass/fail “entrance exam.” That may seem to be a definitive yardstick, but other approaches can be imagined. Instead of Interim Definition criteria, it might be more straightforward to choose a general size-cutoff criterion, where old growth is assigned wherever a certain proportion of trees in a forest attains a particular diameter. A general size-cutoff criterion may have sufficed for early 20^th^-C foresters who relied on “self-evident” standards for declaring old growth versus forest stands of younger growth, but its accuracy may be questioned. For example, working in Douglas-fir stands in coastal Oregon, Tappeiner *et al*. (1997, see their Table 1, p. 641) found that ten old-growth stands (status assumed *a priori*) had an average density of 106 trees per acre and that their proportions of their stems larger than 20 inches in diameter ranged between 43% and 86%; by contrast, young stands had densities of about 152 trees per acre and essentially all trees were under 10 inches in diameter. Can this standard be applied elsewhere?

If, in the above Oregon study, the stand with the lowest proportion of large trees (43% ≥20 inches in diameter) is considered closest to the transition threshold into old growth, then that parameter could be applied as an “oldgrowthness filter” for screening pre-settlement forests in townships around Puget Sound. As indicated in Figure F.1, twenty-three townships (10.2% of the region’s land area) had 43% or more of all witness trees at least 20 inches in diameter: Douglas-fir predominated in fifteen of the townships (nearly all in the region’s southern half), hemlock predominated in only two townships (in the Cascade foothills), and redcedar co-dominated in most of the six remaining townships. Thus, according to this test, perhaps 10% of pre-settlement forests were late-successional; however, neither San Juan County’s Douglas-fir forests nor most of the hemlock-dominated forests of the Cascade foothills were correctly detected as old growth, even though stand ages for both areas have here been estimated at significantly greater than 200 years. This shortcoming remains even when the “oldgrowthness” filter is dropped from 43 to 40%, as also shown in Figure F.1; that filter extension added twenty more townships and 9.8% more land area for a grand total of forty-three and 20% of the land area. Accordingly, it can be concluded that a simple size-cutoff criterion does not adequately detect old growth, especially considering that the criterion is based only on Douglas-fir and that the putative climax species, western hemlock, is inherently disadvantaged by being smaller in diameter.

**Figure F.1.**
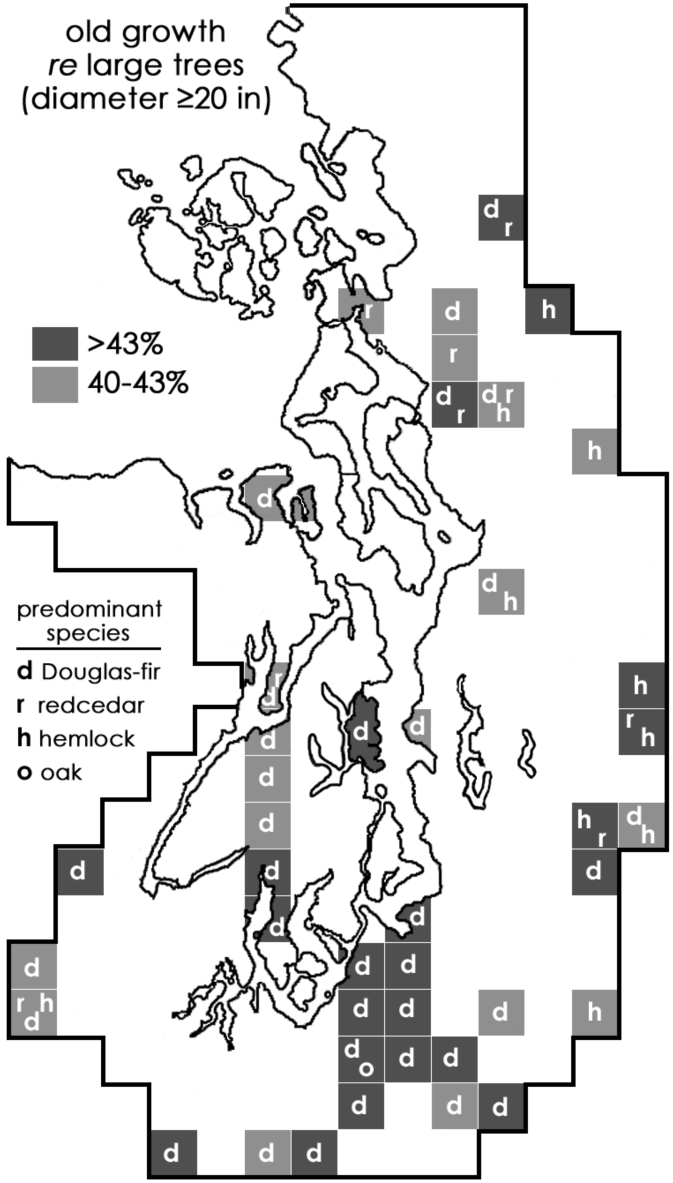
Townships exhibiting old-growth forest characteristics, according to size-cutoff criteria adopted from Tappeiner et al. (1997). Results do not appear to be reasonable.

### G. Age estimations

Tools for estimating stand ages are illustrated in Figure G.1(a-c). The published Site Class Webmap was amended by adding a township grid and by reducing the site-index range of site classes to the midpoint numeric value, identified here as the Nominal Site Index. The map and its site-index values are specific to Douglas-fir; without a separate site-class map specific to hemlock, site-index values for Douglas-fir were adapted as applicable to an area’s *hemlock* component by deducting 5%; thus, a location with a Nominal Site Index of 120 (for Douglas-fir) was judged to have a Nominal Site Index for hemlock of 114. (Hemlock-specific site-quality assessments are far less common in soil surveys and other publications, but where both ratings are reported, commonly for foothill locations, hemlock values are generally about 95% of Douglas-fir counterpart values.)

A township’s singular Nominal Site Index was resolved by inspecting its contents in the Webmap (Figure G.1a) and averaging the relative areas of the constituent Nominal Site Class values, *e.g*., if two-thirds of the township’s area displays a Nominal Site Class value of 107 and the remaining one-third a value of 127, then the township’s assigned Nominal Site Index is 114; ((2×107)+(1×127)) ÷ 3 = 114. This rating process was repeated for all townships for which site-class information is provided in the Site Class map (many urbanized areas were not). This value dictated which specific growth curve in Figure G.1 (b or c) was applicable to the township. By consulting the growth curve, the stand age is the *x*-coordinate value of the point on the curve whose *y*-coordinate has the value of the QMD. Interpolations and extrapolations are applied, as needed.

**Figure G.1.**
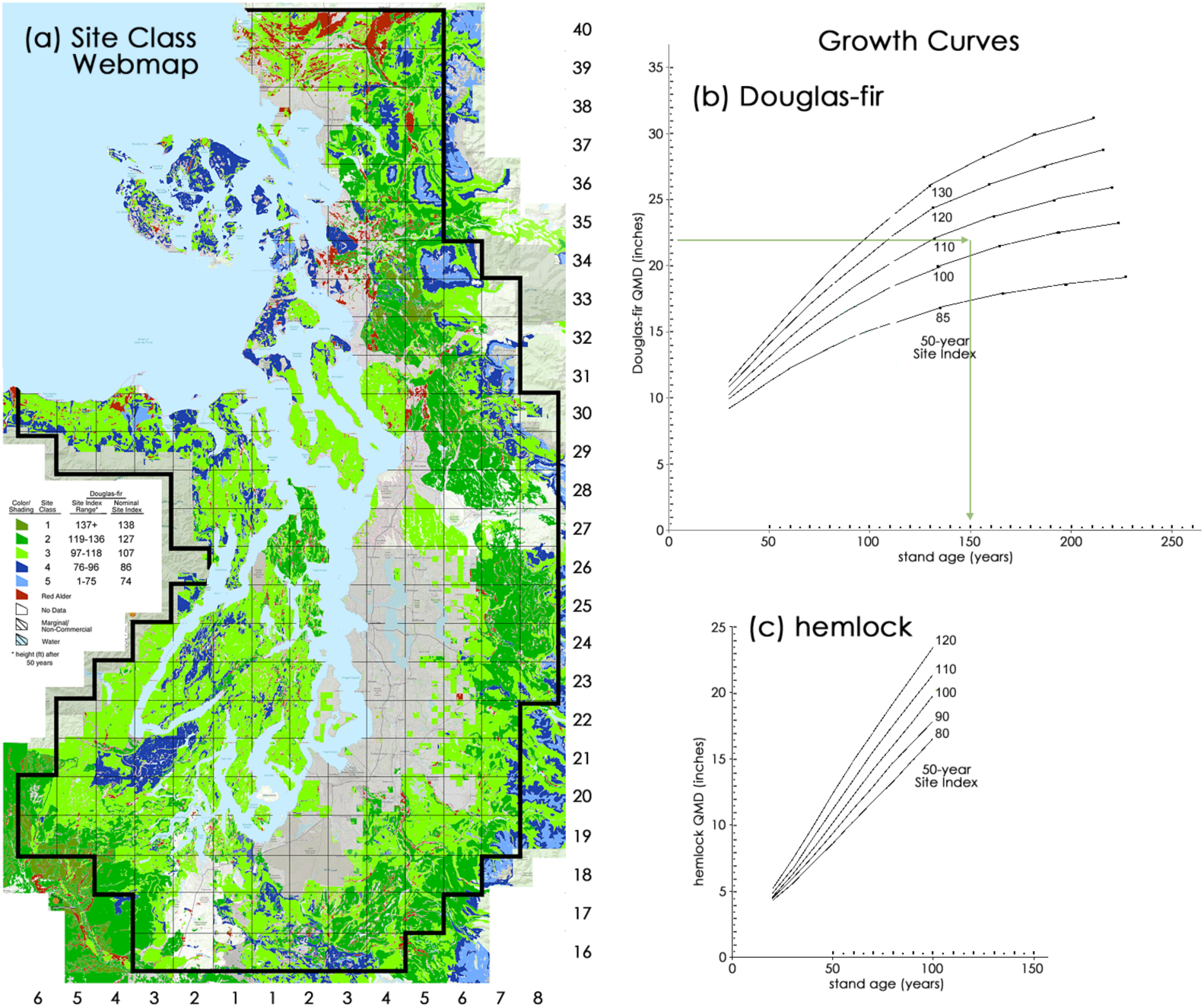
Tools for estimating stand age of a township’s Douglas-fir or hemlock population. (a) Site Class Webmap published online by WA DNR, customized with a township grid and Nominal Site Index values. Strictly speaking, the map is specific to Douglas-fir only; values reduced to 95% adjust it to hemlock. (b) Douglas-fir growth curves, plotted from tables in Chambers (1980); beyond 120 years curves were extrapolated. (c) Hemlock growth curves, plotted from tables in Wiley and Chambers (1981). A township’s Nominal Site Index inferred from map (a) dictates which curve in (b) or (c) is applicable; stand age is gauged by carrying the species’ QMD value (y-axis) across to the selected curve and then down to the x-axis, e.g., green lines in curve (b), where QMD 22 & SI 105 indicates an age of 150 years.

Estimating stand ages retrospectively is an inexact exercise. Procedures and tools include a problematic amount of generalizing, averaging, and interpretation; consequently, the accuracy of results cannot be satisfactorily ascertained. Other issues are even more confounding and unresolved, for example:

1. Is it meaningful to average growth characteristics across very large land areas, such as multiple townships? Unfortunately, GLO witness-tree information is spatially too dilute to provide an alternative.
2. Are mid-20^th^ century tools for describing site qualities and tree growth valid for conditions in the Little Ice Age, when pre-settlement stands developed? If those conditions resulted in slower tree growth, then estimated stand ages should be *increased* by some unknown amount; Gavin *et al*. (2013, p. 32) assert that, indeed, the late Little Ice Age “from about 1750 to 1830, was apparently a period of poor growth for most tree species.”
3. Is it appropriate in age-estimation analyses to pool tree data composed of populations of significantly differing age cohorts? Site indexes, after all, were originally developed from and designed for even-age forest stands, so excluding all large legacy trees (supposedly ancient, if above a certain diameter) would compress the age range of the stand’s remainder and *reduce* its estimated age.

**Figure G.2.**
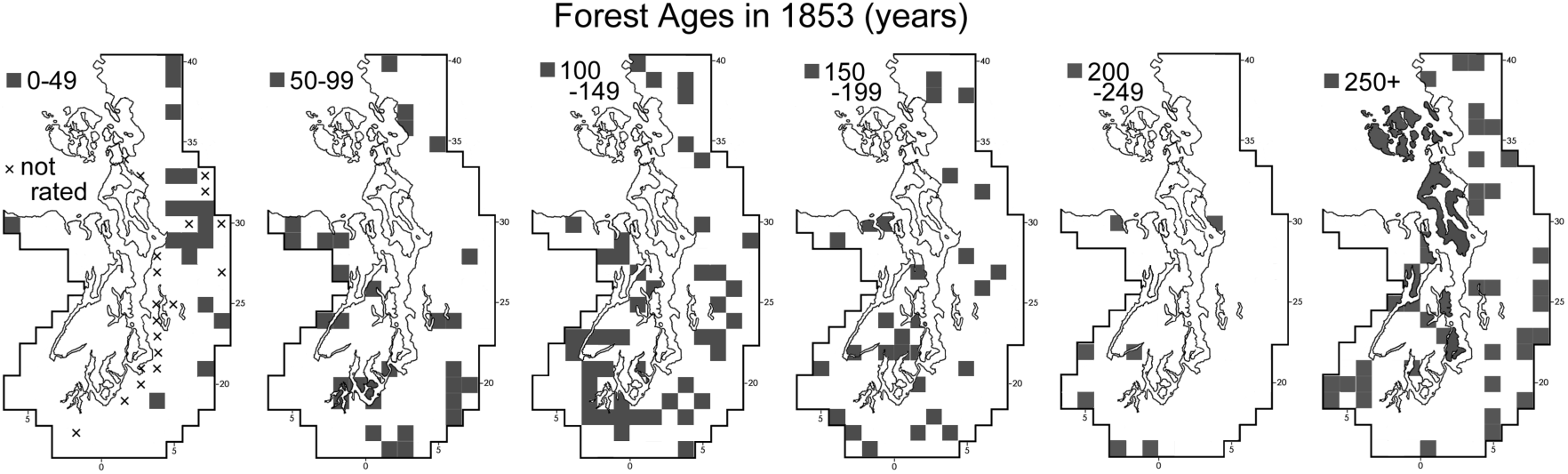
Stand ages for Douglas-fir by township in half-century steps, normalized to 1853. Ages clustered in the 50-149 year century and the 250+ year age classes.

