## Supplementary Materials for "Pre-Settlement Forests around Puget Sound: Eyewitness Evidence"

by

**Tom Schroeder**

This document provides summaries of witness-tree data, compiled for each of the 203 townships, as supplementary information to accompany the parent article, cited above. Provided below is a gridded area map (Figure 1) that readers may use as a worksheet for inserting data for projects. It indicates township locations around Puget Sound and a LiDAR-derived representation of general topography.

The body of this supplement tabulates digested witness-trees data, by township, from their initial GLO surveys. Column headings for the digests are abbreviated as follows:

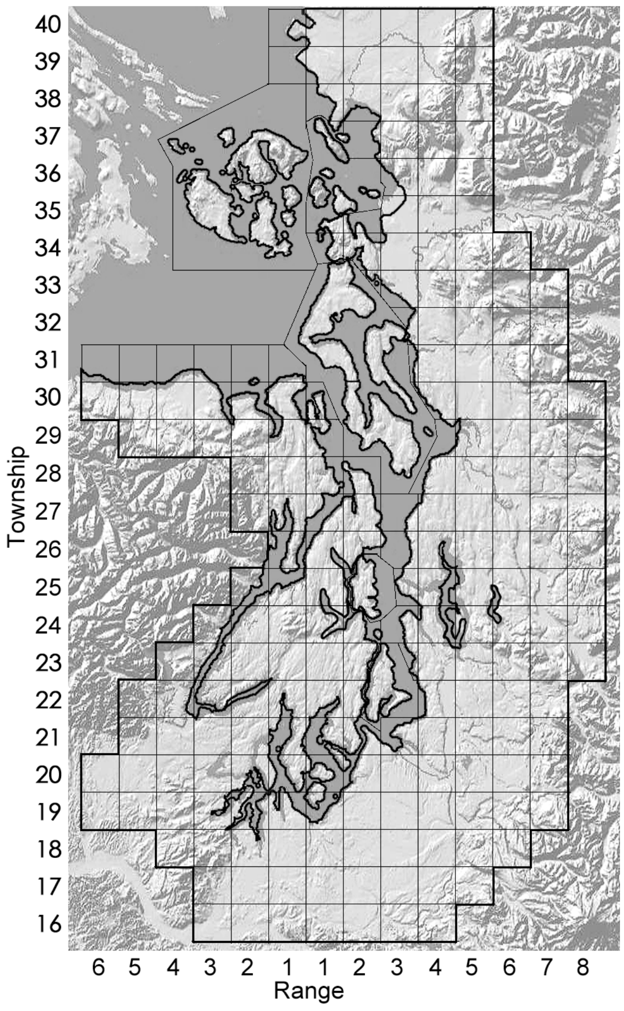


*Figure 1. Template of for entering project data.*

• **Twp/Range**: location coordinates by Township (North) and Range West or East)

• **allWTs**: total tally of all witness trees in the township

• **all avDiam**: average linear diameter of all witness trees, regardless of species, in inches

• **ttl QMD**: quadratic mean diameter (QMD, defined below) of all witness trees in the township

• **SrvyYr**: year date of the GLO survey
•#WTs: number of witness trees of a given species

• **DiamRange**: diameter range, from smallest to largest, within species, in inches

• **avDiam**: average linear diameter in inches

• **MedianDiam**: median diameter, of which an equal number of witness trees was smaller and larger

• **relFreq%**: relative frequency, meaning the percent fraction of a species among "allWTs"

• **QMD**: quadratic mean diameter of witness trees of the species

• **relDOM%**: relative dominance of the species, *i.e.*, the derived diameter in inches of the abstract tree whose cross-sectional area is the average cross-sectional area of all witness trees in a set (of a species or of all all township witness trees, "ttl QMD" above); multiple QMDs cannot be averaged, rather for any new set its QMD must be recomputed from linear diameters of all members of the set).

| Twp-Range | 40-1w &1e | species | # WTs | Diam Range | av Diam | Median Diam | rel Freq% | QMD | rel DOM% |
|---|---|---|---|---|---|---|---|---|---|
| # allWTs | 330 | red alder | 56 | 3to22 | 8.4 | 7 | 16.97 | 9.4 | 3.18 |
| all avDiam | 16.1 | birch | 6 | 4to8 | 5.7 | 5.5 | 1.82 | 5.9 | 0.13 |
| ttl QMD | 21.7 | redcedar | 97 | 5to60 | 27.2 | 24 | 29.39 | 30.8 | 59.13 |
| SrvyYr | 1859 | cherry | 1 | 5 | 5 | 5 | 0.30 | 5 | 0.02 |
| | &72 | crabapple | 2 | 6 | 6 | 6 | 0.61 | 6 | 0.05 |
| | | cottonwood | 4 | 5to12 | 8.5 | 8 | 1.21 | 8.8 | 0.20 |
| | | Douglas-fir | 105 | 4to60 | 15.1 | 7 | 31.82 | 21.8 | 32.06 |
| | | grand fir | 1 | 15 | 15 | 15 | 0.30 | 15 | 0.14 |
| | | hemlock | 7 | 6to20 | 11.1 | 10 | 2.12 | 12 | 0.65 |
| | | bl maple | 23 | 3to30 | 13.2 | 12 | 6.97 | 16.3 | 3.93 |
| | | pine | 1 | 6 | 6 | 6 | 0.30 | 6 | 0.02 |
| | | spruce | 2 | 6to12 | 9 | 9 | 0.61 | 9.5 | 0.12 |
| | | smokewood | 1 | 6 | 6 | 6 | 0.30 | 6 | 0.02 |
| | | vine maple | 1 | 5 | 5 | 5 | 0.30 | 5 | 0.02 |
| | | willow | 23 | 3to10 | 5.1 | 5 | 6.97 | 5.5 | 0.45 |

| Twp-Range | 40-2e | species | # WTs | Diam Range | av Diam | Median Diam | rel Freq% | QMD | rel DOM% |
|---|---|---|---|---|---|---|---|---|---|
| # allWTs | 260 | red alder | 19 | 3to24 | 10.6 | 10 | 7.31 | 11.9 | 2.16 |
| all avDiam | 16.2 | birch | 21 | 3to24 | 8.1 | 8 | 8.08 | 9.2 | 1.43 |
| all QMD | 21.9 | redcedar | 86 | 2to100 | 25 | 20 | 33.08 | 31 | 66.27 |
| SrvyYr | 1871 | cherry | 6 | 2to8 | 3.5 | 2.5 | 2.31 | 4.1 | 0.08 |
| | | crabapple | 4 | 4to14 | 8.8 | 8.5 | 1.54 | 9.4 | 0.28 |
| | | cottonwood | 5 | 4to33 | 14.4 | 10 | 1.92 | 18.1 | 1.31 |
| | | dogwood | 3 | 6to33 | 16.3 | 10 | 1.15 | 20.2 | 0.98 |
| | | Douglas-fir | 35 | 4to36 | 13 | 9 | 13.46 | 16.3 | 7.46 |
| | | hemlock | 14 | 5to24 | 11.2 | 8 | 5.38 | 13 | 1.90 |
| | | hazel | 12 | 3to62 | 18 | 15.5 | 4.62 | 23.7 | 5.40 |
| | | bl maple | 9 | 6to24 | 12.6 | 8 | 3.46 | 14.1 | 1.43 |
| | | pine | 6 | 6to18 | 10 | 9 | 2.31 | 11.2 | 0.60 |
| | | spruce | 16 | 6to70 | 22.6 | 22 | 6.15 | 27.1 | 9.42 |
| | | vine maple | 12 | 4to31 | 7.1 | 5 | 4.62 | 10.2 | 1.00 |
| | | willow | 12 | 3to10 | 5.5 | 6 | 4.62 | 5.8 | 0.32 |

| Twp-Range | 40-3e | species | # WTs | Diam Range | av Diam | Median Diam | rel Freq% | QMD | rel DOM% |
|---|---|---|---|---|---|---|---|---|---|
| # allWTs | 261 | red alder | 39 | 3to30 | 12.7 | 10 | 14.94 | 14.3 | 6.15 |
| all avDiam | 17.6 | birch | 10 | 3to18 | 10.1 | 11 | 3.83 | 11.1 | 0.95 |
| all QMD | 22.3 | redcedar | 61 | 3to72 | 24.5 | 20 | 23.37 | 29.4 | 40.64 |
| SrvyYr | 1873 | cherry | 1 | 15 | 15 | 15 | 0.38 | 15 | 0.17 |
| | | crabapple | 5 | 5to6 | 5.8 | 6 | 1.92 | 5.8 | 0.13 |
| | | cottonwood | 9 | 7to60 | 31.6 | 24 | 3.45 | 38 | 10.02 |
| | | dogwood | 1 | 10 | 10 | 10 | 0.38 | 10 | 0.08 |
| | | Douglas-fir | 22 | 5to60 | 33 | 36 | 8.43 | 36 | 21.97 |
| | | grand fir | 1 | 18 | 18 | 18 | 0.38 | 18 | 0.25 |
| | | hemlock | 25 | 7to40 | 15.2 | 12 | 9.58 | 17.7 | 6.04 |
| | | hazel | 1 | 3 | 3 | 3 | 0.38 | 3 | 0.01 |
| | | bl maple | 18 | 8to40 | 17.3 | 16 | 6.90 | 19.2 | 5.11 |
| | | pine | 21 | 4to12 | 7.5 | 6 | 8.05 | 7.8 | 0.98 |
| | | spruce | 25 | 3to36 | 15.4 | 14 | 9.58 | 17.7 | 6.04 |
| | | vine maple | 12 | 5to31 | 8.4 | 6.5 | 4.60 | 10.9 | 1.10 |
| | | willow | 10 | 3to12 | 7.2 | 6.5 | 3.83 | 7.7 | 0.46 |

| Twp-Range | 40-4e | species | # WTs | Diam Range | av Diam | Median Diam | rel Freq% | QMD | rel DOM% |
|---|---|---|---|---|---|---|---|---|---|
| # allWTs | 226 | red alder | 31 | 4to20 | 9.8 | 10 | 13.72 | 10.7 | 3.65 |
| all avDiam | 16.5 | birch | 5 | 6to8 | 6.4 | 6 | 2.21 | 6.4 | 0.21 |
| all QMD | 20.7 | barberry | 1 | 12 | 12 | 12 | 0.44 | 12 | 0.15 |
| SrvyYr | 1874 | redcedar | 67 | 4to60 | 22.1 | 18 | 29.65 | 25.4 | 44.44 |
| | | cherry | 3 | 5to8 | 6.3 | 6 | 1.33 | 6.5 | 0.13 |
| | | crabapple | 2 | 6to8 | 7 | 7 | 0.88 | 7.1 | 0.10 |
| | | cottonwood | 3 | 10to12 | 11.3 | 12 | 1.33 | 11.4 | 0.40 |
| | | dogwood | 1 | 10 | 10 | 10 | 0.44 | 10 | 0.10 |
| | | Douglas-fir | 41 | 4to72 | 26.4 | 24 | 18.14 | 30.9 | 40.25 |
| | | grand fir | 3 | 12to24 | 16 | 12 | 1.33 | 16.9 | 0.88 |
| | | hemlock | 10 | 10to30 | 16.8 | 14 | 4.42 | 17.8 | 3.26 |
| | | bl maple | 19 | 4to16 | 10.1 | 12 | 8.41 | 10.6 | 2.19 |
| | | spruce | 7 | 6to30 | 17.4 | 16 | 3.10 | 18.7 | 2.52 |
| | | vine maple | 19 | 4to8 | 6 | 6 | 8.41 | 6.2 | 0.75 |
| | | willow | 14 | 3to14 | 6.4 | 4.5 | 6.19 | 7.2 | 0.75 |

| Twp-Range | 40-5e | species | # WTs | Diam Range | av Diam | Median Diam | rel Freq% | QMD | rel DOM% |
|---|---|---|---|---|---|---|---|---|---|
| # allWTs | 261 | red alder | 39 | 4to22 | 10.7 | 10 | 14.94 | 11.6 | 5.40 |
| all avDiam | 13.2 | birch | 10 | 4to14 | 7.8 | 8 | 3.83 | 8.6 | 0.76 |
| all QMD | 19.3 | redcedar | 32 | 6to120 | 33.6 | 22 | 12.26 | 44.2 | 64.33 |
| SrvyYr | 1891 | cherry | 4 | 8to10 | 8.5 | 8 | 1.53 | 8.5 | 0.30 |
| | | dogwood | 1 | 4 | 4 | 4 | 0.38 | 4 | 0.02 |
| | | Douglas-fir | 96 | 4to40 | 9.6 | 8 | 36.78 | 11.9 | 13.99 |
| | | grand fir | 2 | 6 | 6 | 6 | 0.77 | 6 | 0.07 |
| | | hemlock | 40 | 4to36 | 14.9 | 12 | 15.33 | 17.2 | 12.18 |
| | | bl maple | 18 | 4to18 | 10.2 | 10 | 6.90 | 11.2 | 2.32 |
| | | pine | 4 | 2to8 | 6 | 7 | 1.53 | 6.5 | 0.17 |
| | | vine maple | 9 | 4to8 | 4.7 | 4 | 3.45 | 4.9 | 0.22 |
| | | willow | 6 | 4to10 | 7 | 7 | 2.30 | 7.3 | 0.33 |

| Twp-Range | 39-1w | species | # WTs | Diam Range | av Diam | Median Diam | rel Freq% | QMD | rel DOM% |
|---|---|---|---|---|---|---|---|---|---|
| # allWTs | 29 | red alder | 6 | 4to12 | 11.2 | 8.5 | 20.69 | 14.3 | 16.56 |
| all avDiam | 12.4 | redcedar | 3 | 12to36 | 21.3 | 16 | 10.34 | 23.8 | 22.94 |
| all QMD | 16.0 | Douglas-fir | 6 | 4to24 | 11.5 | 9 | 20.69 | 13.6 | 14.98 |
| SrvyYr | 1859 | hemlock | 3 | 6to30 | 18 | 18 | 10.34 | 20.5 | 17.02 |
| | | bl maple | 4 | 2to24 | 10 | 7 | 13.79 | 13 | 9.13 |
| | | spruce | 1 | 36 | 36 | 36 | 3.45 | 36 | 17.50 |
| | | vine maple | 6 | 3to6 | 4.8 | 5 | 20.69 | 5 | 2.03 |

| Twp-Range | 39-1e | species | # WTs | Diam Range | av Diam | Median Diam | rel Freq% | QMD | rel DOM% |
|---|---|---|---|---|---|---|---|---|---|
| # allWTs | 242 | red alder | 37 | 3to24 | 8.5 | 6 | 15.29 | 10.1 | 4.70 |
| all avDiam | 13.2 | birch | 17 | 2to12 | 4.2 | 4 | 7.02 | 4.7 | 0.47 |
| all QMD | 18.2 | redcedar | 43 | 2to80 | 24.5 | 24 | 17.77 | 28.4 | 43.19 |
| SrvyYr | 1859 | cherry | 1 | 3 | 3 | 3 | 0.41 | 3 | 0.01 |
| | | crabapple | 11 | 2to36 | 12.9 | 5 | 4.55 | 17.8 | 4.34 |
| | | cottonwood | 5 | 3to40 | 16.6 | 5 | 2.07 | 22.6 | 3.18 |
| | | Douglas-fir | 11 | 4to72 | 25.5 | 30 | 4.55 | 33.1 | 15.01 |
| | | hemlock | 34 | 5to40 | 18.3 | 20 | 14.05 | 20.4 | 17.62 |
| | | bl maple | 21 | 2to24 | 11 | 8 | 8.68 | 13.3 | 4.63 |
| | | spruce | 5 | 12to48 | 24.4 | 14 | 2.07 | 28.6 | 5.09 |
| | | vine maple | 46 | 3to8 | 4.5 | 4 | 19.01 | 4.6 | 1.21 |
| | | willow | 11 | 2to12 | 5.3 | 4 | 4.55 | 6.1 | 0.51 |

| Twp-Range | 39-2e | species | # WTs | Diam Range | av Diam | Median Diam | rel Freq% | QMD | rel DOM% |
|---|---|---|---|---|---|---|---|---|---|
| # allWTs | 270 | red alder | 60 | 2to20 | 9 | 8.5 | 22.22 | 9.7 | 8.57 |
| all avDiam | 11.5 | ash | 4 | 5to12 | 7.3 | 6 | 1.48 | 7.8 | 0.37 |
| all QMD | 15.6 | birch | 23 | 5to18 | 8.7 | 7 | 8.52 | 9.4 | 3.08 |
| SrvyYr | 1873 | redcedar | 37 | 2to60 | 22.2 | 20 | 13.70 | 26.1 | 38.26 |
| | | cherry | 6 | 3to8 | 4.5 | 3.5 | 2.22 | 4.8 | 0.21 |
| | | crabapple | 3 | 4to10 | 8.5 | 8.5 | 1.11 | 8.5 | 0.33 |
| | | cottonwood | 3 | 6to10 | 7 | 10 | 1.11 | 8.9 | 0.36 |
| | | dogwood | 1 | 8 | 8 | 8 | 0.37 | 8 | 0.10 |
| | | Douglas-fir | 37 | 3to60 | 15.1 | 8 | 13.70 | 21.9 | 26.93 |
| | | grand fir | 4 | 18to60 | 38.5 | 38 | 1.48 | 41.3 | 10.36 |
| | | hemlock | 14 | 4to24 | 10.6 | 8 | 5.19 | 11.9 | 3.01 |
| | | bl maple | 28 | 2to24 | 9.1 | 6 | 10.37 | 11 | 5.14 |
| | | spruce | 6 | 3to18 | 8.8 | 8 | 2.22 | 10 | 0.91 |
| | | skunkwood | 2 | 4to6 | 5 | 5 | 0.74 | 5.1 | 0.08 |
| | | vine maple | 15 | 3to8 | 4.9 | 5 | 5.56 | 5.1 | 0.59 |
| | | willow | 27 | 3to18 | 6.3 | 6 | 10.00 | 7 | 2.01 |

| Twp-Range | 39-3e | species | # WTs | Diam Range | av Diam | Median Diam | rel Freq% | QMD | rel DOM% |
|---|---|---|---|---|---|---|---|---|---|
| # allWTs | 282 | red alder | 57 | 2to14 | 6.9 | 6 | 20.21 | 7.3 | 4.72 |
| all avDiam | 11.4 | birch | 21 | 4to10 | 6.4 | 6 | 7.45 | 6.6 | 1.42 |
| all QMD | 15.1 | barberry | 2 | 6to8 | 7 | 7 | 0.71 | 7.1 | 0.16 |
| SrvyYr | 1874 | redcedar | 46 | 3to40 | 17 | 14 | 16.31 | 18.9 | 25.51 |
| | | cherry | 9 | 2to50 | 13.9 | 5 | 3.19 | 21.3 | 6.34 |
| | | crabapple | 5 | 4to8 | 6 | 6 | 1.77 | 6.1 | 0.29 |
| | | cottonwood | 2 | 8 | 8 | 8 | 0.71 | 8 | 0.20 |
| | | dogwood | 1 | 4 | 4 | 4 | 0.35 | 4 | 0.02 |
| | | Douglas-fir | 52 | 2to60 | 19.2 | 15.5 | 18.44 | 24.4 | 48.07 |
| | | grand fir | 1 | 5 | 5 | 5 | 0.35 | 5 | 0.04 |
| | | hemlock | 16 | 5to15 | 7.6 | 6 | 5.67 | 8.3 | 1.71 |
| | | bl maple | 30 | 2to30 | 10.2 | 8 | 10.64 | 12.5 | 7.28 |
| | | pine | 5 | 4to13 | 9 | 10 | 1.77 | 9.6 | 0.72 |
| | | spruce | 5 | 8to24 | 14.6 | 15 | 1.77 | 15.6 | 1.89 |
| | | skunkwood | 1 | 4 | 4 | 4 | 0.35 | 4 | 0.02 |
| | | vine maple | 8 | 2to50 | 3.9 | 4 | 2.84 | 4 | 0.20 |
| | | willow | 21 | 3to12 | 5.7 | 5 | 7.45 | 6.2 | 1.25 |

| Twp-Range | 39-4e | species | # WTs | Diam Range | av Diam | Median Diam | rel Freq% | QMD | rel DOM% |
|---|---|---|---|---|---|---|---|---|---|
| # allWTs | 268 | red alder | 33 | 3to15 | 7.6 | 7 | 12.31 | 8.4 | 1.79 |
| all avDiam | 17.0 | birch | 5 | 3to6 | 4.4 | 4 | 1.87 | 4.5 | 0.08 |
| all QMD | 22.1 | barberry | 3 | 5to8 | 6 | 5 | 1.12 | 6.2 | 0.09 |
| SrvyYr | 1884 | redcedar | 82 | 4to70 | 25.8 | 20 | 30.60 | 30.4 | 58.15 |
| | | cottonwood | 8 | 4to30 | 11 | 6 | 2.99 | 14.2 | 1.24 |
| | | dogwood | 1 | 6 | 6 | 6 | 0.37 | 6 | 0.03 |
| | | Douglas-fir | 61 | 3to60 | 19.3 | 18 | 22.76 | 23.7 | 26.29 |
| | | grand fir | 1 | 36 | 36 | 36 | 0.37 | 36 | 0.99 |
| | | hemlock | 30 | 3to48 | 16.1 | 14 | 11.19 | 18.8 | 8.14 |
| | | bl maple | 16 | 5to25 | 11.8 | 8 | 5.97 | 13.6 | 2.27 |
| | | pine | 1 | 20 | 20 | 20 | 0.37 | 20 | 0.31 |
| | | spruce | 2 | 8to12 | 10 | 10 | 0.75 | 10.2 | 0.16 |
| | | vine maple | 20 | 3to12 | 5.1 | 4.5 | 7.46 | 5.5 | 0.46 |
| | | willow | 5 | 3to5 | 4 | 4 | 1.87 | 4 | 0.06 |

| Twp-Range | 39-5e | species | # WTs | Diam Range | av Diam | Median Diam | rel Freq% | QMD | rel DOM% |
|---|---|---|---|---|---|---|---|---|---|
| # allWTs | 252 | red alder | 38 | 4to24 | 10.2 | 9 | 15.08 | 11.6 | 5.96 |
| all avDiam | 14.8 | birch | 1 | 10 | 10 | 10 | 0.40 | 10 | 0.12 |
| all QMD | 18.5 | redcedar | 48 | 3to72 | 22.1 | 19 | 19.05 | 26.4 | 38.99 |
| SrvyYr | 1890 | cherry | 1 | 6 | 6 | 6 | 0.40 | 6 | 0.04 |
| | | Douglas-fir | 49 | 4to36 | 11.7 | 10 | 19.44 | 14.3 | 11.68 |
| | | hemlock | 74 | 4to48 | 18.3 | 16 | 29.37 | 21 | 38.04 |
| | | bl maple | 12 | 4to36 | 14 | 10 | 4.76 | 16.6 | 3.85 |
| | | spruce | 2 | 8to10 | 9 | 9 | 0.79 | 9.1 | 0.19 |
| | | cascara | 4 | 6to10 | 9 | 10 | 1.59 | 9.2 | 0.39 |
| | | vine maple | 23 | 4to8 | 4.8 | 4 | 9.13 | 4.9 | 0.64 |

| Twp-Range | 38-1e | species | # WTs | Diam Range | av Diam | Median Diam | rel Freq% | QMD | rel DOM% |
|---|---|---|---|---|---|---|---|---|---|
| # allWTs | 82 | red alder | 15 | 3to14 | 7.7 | 6 | 18.29 | 8.7 | 4.36 |
| all avDiam | 12.9 | redcedar | 4 | 24to48 | 34.5 | 33 | 4.88 | 35.6 | 19.47 |
| all QMD | 17.8 | crabapple | 8 | 3to24 | 8.3 | 5 | 9.76 | 10.7 | 3.52 |
| SrvyYr | 1859&73 | cottonwood | 3 | 3to5 | 4.3 | 5 | 3.66 | 4.4 | 0.22 |
| | | Douglas-fir | 7 | 12to48 | 34.9 | 36 | 8.54 | 38.1 | 39.03 |
| | | hemlock | 9 | 3to30 | 18.8 | 24 | 10.98 | 20.5 | 14.53 |
| | | bl maple | 19 | 3to30 | 12.3 | 12 | 23.17 | 15.3 | 17.08 |
| | | spruce | 3 | 8to10 | 8.7 | 8 | 3.66 | 8.7 | 0.87 |
| | | vine maple | 10 | 3to6 | 4 | 4 | 12.20 | 4.1 | 0.65 |
| | | willow | 4 | 3to5 | 3.8 | 3.5 | 4.88 | 3.8 | 0.22 |

| Twp-Range | 38-2e | species | # WTs | Diam Range | av Diam | Median Diam | rel Freq% | QMD | rel DOM% |
|---|---|---|---|---|---|---|---|---|---|
| # allWTs | 144 | red alder | 15 | 2to30 | 10.3 | 10 | 10.42 | 12.5 | 2.87 |
| all avDiam | 19.2 | redcedar | 31 | 8to60 | 23.5 | 20 | 21.53 | 26.1 | 25.86 |
| all QMD | 23.8 | crabapple | 6 | 4to16 | 8.2 | 7 | 4.17 | 9.1 | 0.61 |
| SrvyYr | 1859 | cottonwood | 1 | 3 | 3 | 3 | 0.69 | 3 | 0.01 |
| | &73 | Douglas-fir | 31 | 4to60 | 32.8 | 35 | 21.53 | 36.7 | 51.13 |
| | | hemlock | 31 | 4to40 | 18.7 | 18 | 21.53 | 20.8 | 16.42 |
| | | bl maple | 15 | 3to24 | 10.1 | 8 | 10.42 | 11.7 | 2.51 |
| | | vine maple | 3 | 5to6 | 5.7 | 6 | 2.08 | 5.7 | 0.12 |
| | | willow | 11 | 3to10 | 5.9 | 6 | 7.64 | 6.2 | 0.52 |

| Twp-Range | 38-3e | species | # WTs | Diam Range | av Diam | Median Diam | rel Freq% | QMD | rel DOM% |
|---|---|---|---|---|---|---|---|---|---|
| # allWTs | 260 | red alder | 47 | 2to20 | 6.6 | 5 | 18.08 | 7.7 | 3.74 |
| all avDiam | 11.9 | birch | 10 | 4to10 | 5.4 | 5 | 3.85 | 5.7 | 0.44 |
| all QMD | 16.9 | redcedar | 42 | 3to48 | 20.5 | 18 | 16.15 | 23.1 | 30.11 |
| SrvyYr | 1874 | cherry | 10 | 2to10 | 4.6 | 3.5 | 3.85 | 5.4 | 0.39 |
| | | cottonwood | 8 | 6to40 | 13.8 | 10 | 3.08 | 17.3 | 3.22 |
| | | Douglas-fir | 60 | 2to60 | 19.5 | 9.5 | 23.08 | 25.9 | 54.07 |
| | | grand fir | 3 | 5to8 | 6.3 | 6 | 1.15 | 6.5 | 0.17 |
| | | hemlock | 19 | 4to40 | 11.1 | 8 | 7.31 | 14.2 | 5.15 |
| | | bl maple | 15 | 3to12 | 6.1 | 5 | 5.77 | 6.9 | 0.96 |
| | | pine | 4 | 4to14 | 7.3 | 5.5 | 1.54 | 8.3 | 0.37 |
| | | vine maple | 4 | 4to8 | 5 | 4 | 1.54 | 5.3 | 0.15 |
| | | willow | 38 | 4to9 | 4.8 | 5 | 14.62 | 5 | 1.28 |

| Twp-Range | 38-4e | species | # WTs | Diam Range | av Diam | Median Diam | rel Freq% | QMD | rel DOM% |
|---|---|---|---|---|---|---|---|---|---|
| # allWTs | 261 | red alder | 16 | 2to14 | 8.1 | 8 | 6.13 | 8.5 | 0.95 |
| all avDiam | 17.2 | birch | 2 | 6to11 | 8.5 | 8.5 | 0.77 | 8.9 | 0.13 |
| all QMD | 21.5 | redcedar | 67 | 6to62 | 23.5 | 20 | 25.67 | 27.4 | 41.52 |
| SrvyYr | 1884 | cherry | 3 | 2to8 | 4.3 | 3 | 1.15 | 5.1 | 0.06 |
| | | Douglas-fir | 54 | 2to72 | 17.6 | 12 | 20.69 | 23.3 | 24.20 |
| | | hemlock | 98 | 3to48 | 16.4 | 12.5 | 37.55 | 19.4 | 30.44 |
| | | bl maple | 11 | 3to24 | 11.5 | 10 | 4.21 | 13.7 | 1.70 |
| | | pine | 2 | 14to24 | 17 | 17 | 0.77 | 19.6 | 0.63 |
| | | spruce | 1 | 12 | 12 | 12 | 0.38 | 12 | 0.12 |
| | | vine maple | 4 | 3to5 | 4 | 4 | 1.53 | 4.3 | 0.06 |
| | | willow | 2 | 3 | 3 | 3 | 0.77 | 3 | 0.01 |
| | | yew | 1 | 5 | 5 | 5 | 0.38 | 5 | 0.02 |

| Twp-Range | 38-5e | species | # WTs | Diam Range | av Diam | Median Diam | rel Freq% | QMD | rel DOM% |
|---|---|---|---|---|---|---|---|---|---|
| # allWTs | 111 | red alder | 5 | 6to24 | 12 | 8 | 4.50 | 13.8 | 1.64 |
| all avDiam | 18.1 | redcedar | 16 | 10to60 | 27.4 | 22 | 14.41 | 31.3 | 26.93 |
| all QMD | 22.9 | cherry | 1 | 6 | 6 | 6 | 0.90 | 6 | 0.06 |
| SrvyYr | 1895 | Douglas-fir | 40 | 4to70 | 20.8 | 12 | 36.04 | 27.6 | 52.34 |
| | | hemlock | 40 | 4to36 | 14.3 | 13 | 36.04 | 15.6 | 16.72 |
| | | larch | 1 | 10 | 10 | 10 | 0.90 | 10 | 0.17 |
| | | bl maple | 6 | 8to16 | 10.8 | 9 | 5.41 | 11.3 | 1.32 |
| | | spruce | 1 | 19 | 19 | 19 | 0.90 | 19 | 0.62 |
| | | vine maple | 1 | 4 | 4 | 4 | 0.90 | 4 | 0.03 |

| Twp-Range | 37-3e | species | # WTs | Diam Range | av Diam | Median Diam | rel Freq% | QMD | rel DOM% |
|---|---|---|---|---|---|---|---|---|---|
| # allWTs | 280 | red alder | 25 | 3to16 | 7.8 | 8 | 8.93 | 8.4 | 1.64 |
| all avDiam | 15.8 | redcedar | 69 | 4to60 | 23.6 | 24 | 24.64 | 26.3 | 44.37 |
| all QMD | 19.6 | cherry | 4 | 3to12 | 6.5 | 5.5 | 1.43 | 7.3 | 0.20 |
| SrvyYr | 1873 | Douglas-fir | 73 | 3to60 | 15 | 6 | 26.07 | 20.3 | 27.96 |
| | | hemlock | 87 | 2to36 | 14.5 | 12 | 31.07 | 16.7 | 22.55 |
| | | bl maple | 11 | 3to18 | 10.4 | 10 | 3.93 | 11.4 | 1.33 |
| | | spruce | 2 | 30 | 30 | 30 | 0.71 | 30 | 1.67 |
| | | vine maple | 1 | 6 | 6 | 6 | 0.36 | 6 | 0.03 |
| | | willow | 8 | 3to6 | 4.5 | 4 | 2.86 | 4.7 | 0.16 |

| Twp-Range | 37-4e | species | # WTs | Diam Range | av Diam | Median Diam | rel Freq% | QMD | rel DOM% |
|---|---|---|---|---|---|---|---|---|---|
| # allWTs | 259 | red alder | 8 | 5to10 | 7.1 | 6.5 | 3.09 | 7.4 | 0.22 |
| all avDiam | 20.4 | birch | 1 | 6 | 6 | 6 | 0.39 | 6 | 0.02 |
| all QMD | 27.6 | redcedar | 72 | 3to180 | 30.8 | 24 | 27.80 | 41 | 61.27 |
| SrvyYr | 1883 | cherry | 1 | 4 | 4 | 4 | 0.39 | 4 | 0.01 |
| | | Douglas-fir | 42 | 2to55 | 20.9 | 8.5 | 16.22 | 27.9 | 16.55 |
| | | hemlock | 122 | 4to48 | 16.1 | 14 | 47.10 | 18.1 | 20.23 |
| | | bl maple | 8 | 5to24 | 10.4 | 7 | 3.09 | 12.1 | 0.59 |
| | | pine | 1 | 28 | 28 | 28 | 0.39 | 28 | 0.40 |
| | | spruce | 1 | 30 | 30 | 30 | 0.39 | 30 | 0.46 |
| | | willow | 2 | 5to8 | 6.5 | 6.5 | 0.77 | 6.7 | 0.05 |
| | | yew | 1 | 7 | 7 | 7 | 0.39 | 7 | 0.02 |

| Twp-Range | 37-5e | species | # WTs | Diam Range | av Diam | Median Diam | rel Freq% | QMD | rel DOM% |
|---|---|---|---|---|---|---|---|---|---|
| # allWTs | 288 | red alder | 31 | 3to24 | 11.4 | 10 | 10.76 | 12.4 | 4.43 |
| all avDiam | 14.6 | barberry | 2 | 5to10 | 7.5 | 7.5 | 0.69 | 7.9 | 0.12 |
| all QMD | 19.3 | redcedar | 59 | 7to108 | 23 | 18 | 20.49 | 30.2 | 50.05 |
| SrvyYr | 1885 | cherry | 3 | 5to6 | 5.7 | 6 | 1.04 | 5.7 | 0.09 |
| | | cottonwood | 1 | 30 | 30 | 30 | 0.35 | 30 | 0.84 |
| | | Douglas-fir | 42 | 3to60 | 11.8 | 7 | 14.58 | 17.5 | 11.96 |
| | | hemlock | 129 | 3to48 | 13.3 | 11 | 44.79 | 15.1 | 27.36 |
| | | bl maple | 2 | 5to12 | 8.5 | 8.5 | 0.69 | 9.2 | 0.16 |
| | | pine | 2 | 4to6 | 5 | 5 | 0.69 | 5.1 | 0.05 |
| | | spruce | 5 | 8to60 | 25.4 | 20 | 1.74 | 31.2 | 4.53 |
| | | cascara | 1 | 8 | 8 | 8 | 0.35 | 8 | 0.06 |
| | | vine maple | 7 | 3to6 | 4.3 | 4 | 2.43 | 4.4 | 0.13 |
| | | willow | 2 | 6to8 | 7 | 7 | 0.69 | 7.1 | 0.09 |
| | | yew | 2 | 8to9 | 8.5 | 8.5 | 0.69 | 8.5 | 0.13 |

| Twp-Range | 36-3e | species | # WTs | Diam Range | av Diam | Median Diam | rel Freq% | QMD | rel DOM% |
|---|---|---|---|---|---|---|---|---|---|
| # allWTs | 191 | red alder | 11 | 6to24 | 11.9 | 8 | 5.76 | 13.2 | 2.09 |
| all avDiam | 18.0 | birch | 4 | 6to10 | 7 | 6 | 2.09 | 7.2 | 0.23 |
| all QMD | 21.9 | redcedar | 34 | 6to60 | 22.2 | 19 | 17.80 | 25 | 23.13 |
| SrvyYr | 1872 | crabapple | 2 | 6to8 | 7 | 7 | 1.05 | 7.1 | 0.11 |
| | | Douglas-fir | 29 | 4to60 | 20.1 | 15 | 15.18 | 25.8 | 21.01 |
| | | grand fir | 2 | 12to48 | 30 | 30 | 1.05 | 35 | 2.67 |
| | | hemlock | 95 | 5to48 | 17.6 | 16 | 49.74 | 20.5 | 43.45 |
| | | bl maple | 3 | 6to24 | 15.3 | 16 | 1.57 | 17 | 0.94 |
| | | spruce | 2 | 20to72 | 46 | 46 | 1.05 | 52.6 | 6.02 |
| | | vine maple | 6 | 6to7 | 6.2 | 6 | 3.14 | 6.2 | 0.25 |
| | | willow | 1 | 10 | 10 | 10 | 0.52 | 10 | 0.11 |
| | | yew | 2 | 6to7 | 6.5 | 6.5 | 1.05 | 6.5 | 0.09 |

| Twp-Range | 36-4e | species | # WTs | Diam Range | av Diam | Median Diam | rel Freq% | QMD | rel DOM% |
|---|---|---|---|---|---|---|---|---|---|
| # allWTs | 287 | red alder | 22 | 4to30 | 15.1 | 14 | 7.67 | 16.6 | 5.08 |
| all avDiam | 17.3 | redcedar | 57 | 5to60 | 23 | 20 | 19.86 | 26.5 | 33.56 |
| all QMD | 20.4 | cherry | 3 | 8to16 | 13.3 | 16 | 1.05 | 13.9 | 0.49 |
| SrvyYr | 1874 | Douglas-fir | 12 | 5to60 | 26.4 | 28 | 4.18 | 30.1 | 9.11 |
| | | hemlock | 166 | 4to48 | 16.1 | 14 | 57.84 | 18.5 | 47.63 |
| | | bl maple | 10 | 6to30 | 13.2 | 10 | 3.48 | 15.1 | 1.91 |
| | | spruce | 5 | 10to30 | 18.6 | 15 | 1.74 | 20 | 1.68 |
| | | vine maple | 11 | 5to8 | 5.8 | 6 | 3.83 | 5.9 | 0.32 |
| | | willow | 1 | 6 | 6 | 6 | 0.35 | 6 | 0.03 |

| Twp-Range | 36-5e | species | # WTs | Diam Range | av Diam | Median Diam | rel Freq% | QMD | rel DOM% |
|---|---|---|---|---|---|---|---|---|---|
| # allWTs | 241 | red alder | 8 | 8to70 | 21 | 16 | 3.32 | 28.2 | 3.65 |
| all avDiam | 22.6 | redcedar | 41 | 8to70 | 30.1 | 30 | 17.01 | 35.5 | 29.68 |
| all QMD | 26.9 | Douglas-fir | 29 | 8to72 | 24.3 | 14 | 12.03 | 30.1 | 15.09 |
| SrvyYr | 1890 | grand fir | 13 | 4to30 | 17.1 | 12 | 5.39 | 19.4 | 2.81 |
| | | hemlock | 132 | 4to60 | 21.7 | 20 | 54.77 | 24.3 | 44.77 |
| | | bl maple | 6 | 4to20 | 11.3 | 11 | 2.49 | 12.3 | 0.52 |
| | | spruce | 4 | 8to60 | 35 | 36 | 1.66 | 39.5 | 3.58 |
| | | vine maple | 5 | 4to7 | 5.8 | 6 | 2.07 | 5.9 | 0.10 |
| | | yew | 3 | 8to12 | 9.3 | 8 | 1.24 | 9.5 | 0.16 |

| Twp-Range | 35-3e | species | # WTs | Diam Range | av Diam | Median Diam | rel Freq% | QMD | rel DOM% |
|---|---|---|---|---|---|---|---|---|---|
| # allWTs | 226 | red alder | 47 | 3to30 | 11.6 | 10 | 20.80 | 12.7 | 7.22 |
| all avDiam | 18.0 | barberry | 6 | 6to12 | 8 | 8 | 2.65 | 8.2 | 0.38 |
| all QMD | 21.6 | redcedar | 54 | 8to48 | 25.5 | 24 | 23.89 | 27.7 | 39.45 |
| SrvyYr | 1870 | crabapple | 7 | 6to9 | 7.4 | 8 | 3.10 | 7.5 | 0.37 |
| | | cottonwood | 4 | 8to16 | 12.3 | 12.5 | 1.77 | 12.7 | 0.61 |
| | | Douglas-fir | 16 | 3to60 | 26 | 21 | 7.08 | 31.3 | 14.92 |
| | | grand fir | 5 | 10to40 | 23 | 15 | 2.21 | 26.9 | 3.44 |
| | | hemlock | 35 | 6to40 | 17.7 | 18 | 15.49 | 19.5 | 12.67 |
| | | bl maple | 8 | 8to20 | 14.8 | 15 | 3.54 | 15.3 | 1.78 |
| | | pine | 2 | 12to14 | 13 | 13 | 0.88 | 13 | 0.32 |
| | | spruce | 23 | 8to60 | 23.3 | 18 | 10.18 | 27.8 | 16.92 |
| | | vine maple | 2 | 6to8 | 7 | 7 | 0.88 | 7 | 0.09 |
| | | willow | 16 | 5to18 | 9.3 | 8 | 7.08 | 10 | 1.52 |
| | | yew | 1 | 12 | 12 | 12 | 0.44 | 12 | 0.14 |

| Twp-Range | 35-4e | species | # WTs | Diam Range | av Diam | Median Diam | rel Freq% | QMD | rel DOM% |
|---|---|---|---|---|---|---|---|---|---|
| # allWTs | 244 | red alder | 30 | 4to24 | 11.8 | 10 | 12.30 | 13 | 4.72 |
| all avDiam | 16.4 | birch | 5 | 3to18 | 8 | 6 | 2.05 | 9.6 | 0.43 |
| all QMD | 21.0 | barberry | 1 | 6 | 6 | 6 | 0.41 | 6 | 0.03 |
| SrvyYr | 1873 | redcedar | 52 | 3to108 | 27.5 | 20 | 21.31 | 33.4 | 53.98 |
| | | cherry | 3 | 5to6 | 5.7 | 6 | 1.23 | 5.7 | 0.09 |
| | | crabapple | 2 | 5to6 | 5.5 | 5.5 | 0.82 | 5.5 | 0.06 |
| | | cottonwood | 3 | 16to36 | 25.5 | 24 | 1.23 | 36.6 | 3.74 |
| | | Douglas-fir | 20 | 6to50 | 20.8 | 16.5 | 8.20 | 24.7 | 11.35 |
| | | hemlock | 78 | 4to36 | 14.4 | 12 | 31.97 | 16.1 | 18.81 |
| | | hazel | 1 | 4 | 4 | 4 | 0.41 | 4 | 0.01 |
| | | bl maple | 27 | 4to40 | 11.2 | 8 | 11.07 | 13.6 | 4.65 |
| | | spruce | 10 | 14to40 | 15.5 | 12 | 4.10 | 18.9 | 3.32 |
| | | vine maple | 8 | 5to8 | 6.1 | 6 | 3.28 | 6.2 | 0.29 |
| | | willow | 4 | 6to8 | 6.8 | 6.5 | 1.64 | 6.8 | 0.17 |

| Twp-Range | 35-5e | species | # WTs | Diam Range | av Diam | Median Diam | rel Freq% | QMD | rel DOM% |
|---|---|---|---|---|---|---|---|---|---|
| # allWTs | 280 | red alder | 36 | 4to24 | 11.2 | 13 | 12.86 | 14.8 | 4.18 |
| all avDiam | 19.2 | redcedar | 49 | 4to150 | 41.2 | 40 | 17.50 | 48.4 | 60.80 |
| QMD | 26.0 | crabapple | 2 | 10to12 | 11 | 11 | 0.71 | 11 | 0.13 |
| SrvyYr | 1877 | cottonwood | 6 | 4to40 | 21 | 20 | 2.14 | 23.8 | 1.80 |
| | | elder | 2 | 4 | 4 | 4 | 0.71 | 4 | 0.02 |
| | | Douglas-fir | 21 | 4to60 | 20.2 | 18 | 7.50 | 24 | 6.41 |
| | | grand fir | 1 | 14 | 14 | 14 | 0.36 | 14 | 0.10 |
| | | hemlock | 95 | 4to40 | 14.2 | 12 | 33.93 | 16.2 | 13.21 |
| | | hazel | 1 | 4 | 4 | 4 | 0.36 | 4 | 0.01 |
| | | bl maple | 13 | 8to60 | 20.3 | 12 | 4.64 | 24.1 | 4.00 |
| | | pine | 1 | 18 | 18 | 18 | 0.36 | 18 | 0.17 |
| | | spruce | 15 | 10to60 | 29.7 | 24 | 5.36 | 33.2 | 8.76 |
| | | cascara | 3 | 4to6 | 5 | 5 | 1.07 | 5.1 | 0.04 |
| | | vine maple | 31 | 3to6 | 4.4 | 4 | 11.07 | 4.5 | 0.33 |
| | | willow | 3 | 4 | 4 | 4 | 1.07 | 4 | 0.03 |
| | | yew | 1 | 10 | 10 | 10 | 0.36 | 10 | 0.05 |

| Twp-Range | 34-2e | species | # WTs | Diam Range | av Diam | Median Diam | rel Freq% | QMD | rel DOM% |
|---|---|---|---|---|---|---|---|---|---|
| # allWTs | 137 | red alder | 23 | 6to18 | 9 | 8 | 16.79 | 9.4 | 2.45 |
| all avDiam | 19.7 | redcedar | 18 | 6to48 | 26.7 | 28 | 13.14 | 28.2 | 17.26 |
| all QMD | 24.6 | cherry | 2 | 6to8 | 7 | 7 | 1.46 | 7.1 | 0.12 |
| SrvyYr | 1871 | cottonwood | 1 | 6 | 6 | 6 | 0.73 | 6 | 0.04 |
| | | Douglas-fir | 71 | 4to60 | 23.9 | 20 | 51.82 | 29.1 | 72.49 |
| | | grand fir | 2 | 18to40 | 29 | 29 | 1.46 | 21 | 1.06 |
| | | hemlock | 7 | 6to24 | 15.1 | 18 | 5.11 | 16.3 | 2.24 |
| | | spruce | 1 | 30 | 30 | 30 | 0.73 | 30 | 1.09 |
| | | willow | 12 | 6to24 | 8.1 | 6 | 8.76 | 9.4 | 1.28 |

| Twp-Range | 34-3e | species | # WTs | Diam Range | av Diam | Median Diam | rel Freq% | QMD | rel DOM% |
|---|---|---|---|---|---|---|---|---|---|
| # allWTs | 207 | red alder | 61 | 3to48 | 10.4 | 9 | 29.47 | 12.6 | 14.98 |
| all avDiam | 13.2 | birch | 7 | 5to12 | 10.1 | 12 | 3.38 | 10.5 | 1.19 |
| all QMD | 17.7 | barberry | 3 | 6to8 | 7 | 7 | 1.45 | 7 | 0.23 |
| SrvyYr | 1872 | redcedar | 29 | 3to60 | 21.4 | 12 | 14.01 | 26.9 | 32.46 |
| | | crabapple | 6 | 3to7 | 4.5 | 4 | 2.90 | 4.7 | 0.20 |
| | | cottonwood | 2 | 8to16 | 12 | 12 | 0.97 | 12.6 | 0.49 |
| | | elder | 1 | 6 | 6 | 6 | 0.48 | 6 | 0.06 |
| | | Douglas-fir | 5 | 3to24 | 15.6 | 15 | 2.42 | 17.5 | 2.37 |
| | | grand fir | 5 | 6to24 | 11.8 | 10 | 2.42 | 13.5 | 1.41 |
| | | hazel | 1 | 6 | 6 | 6 | 0.48 | 6 | 0.06 |
| | | hemlock | 2 | 6to7 | 6.5 | 6.5 | 0.97 | 6.5 | 0.13 |
| | | juniper | 6 | 6to24 | 12.3 | 10 | 2.90 | 14 | 1.82 |
| | | bl maple | 7 | 4to20 | 11.7 | 12 | 3.38 | 12.9 | 1.80 |
| | | spruce | 34 | 5to60 | 22.4 | 15 | 16.43 | 27.3 | 39.19 |
| | | vine maple | 5 | 4to6 | 4.6 | 4 | 2.42 | 4.7 | 0.17 |
| | | willow | 33 | 3to24 | 7.2 | 6 | 15.94 | 8.2 | 3.43 |

| Twp-Range | 34-4e | species | # WTs | Diam Range | av Diam | Median Diam | rel Freq% | QMD | rel DOM% |
|---|---|---|---|---|---|---|---|---|---|
| # allWTs | 273 | red alder | 36 | 5to36 | 12.9 | 12 | 13.19 | 14 | 3.54 |
| all avDiam | 21.89 | barberry | 1 | 12 | 12 | 12 | 0.37 | 12 | 0.07 |
| all QMD | 27.01 | birch | 1 | 7 | 7 | 7 | 0.37 | 7 | 0.02 |
| SrvyYr | 1872 | redcedar | 78 | 6to96 | 24 | 35.9 | 28.57 | 35.9 | 50.48 |
| | | cherry | 4 | 10to15 | 11.3 | 10 | 1.47 | 11.5 | 0.27 |
| | | cottonwood | 5 | 10to36 | 30.8 | 36 | 1.83 | 32.5 | 2.65 |
| | | crabapple | 4 | 6to15 | 10.8 | 11 | 1.47 | 11.2 | 0.25 |
| | | Douglas-fir | 22 | 6to84 | 26.8 | 24 | 8.06 | 32.6 | 11.74 |
| | | grand fir | 1 | 18 | 18 | 18 | 0.37 | 18 | 0.16 |
| | | hemlock | 66 | 5to80 | 19 | 16 | 24.18 | 22.2 | 16.33 |
| | | bl maple | 19 | 4to24 | 12.8 | 12 | 6.96 | 13.7 | 1.79 |
| | | spruce | 23 | 6to60 | 28.7 | 30 | 8.42 | 32.5 | 12.20 |
| | | vine maple | 6 | 4to8 | 6.2 | 6 | 2.20 | 6.3 | 0.12 |
| | | willow | 7 | 4to18 | 11.3 | 10 | 2.56 | 12 | 0.51 |

| Twp-Range | 34-5e | species | # WTs | Diam Range | av Diam | Median Diam | rel Freq% | QMD | rel DOM% |
|---|---|---|---|---|---|---|---|---|---|
| # allWTs | 288 | red alder | 9 | 5to14 | 10.3 | 12 | 3.13 | 10.7 | 0.69 |
| all avDiam | 18.20 | barberry | 1 | 3 | 3 | 3 | 0.35 | 3 | 0.01 |
| all QMD | 22.77 | redcedar | 44 | 6to108 | 24 | 19 | 15.28 | 30.2 | 26.87 |
| SrvyYr | 1883& | Douglas-fir | 13 | 6to50 | 15.9 | 10 | 4.51 | 20.4 | 3.62 |
| | 1900 | hemlock | 145 | 2to72 | 17.6 | 14 | 50.35 | 21.8 | 46.13 |
| | | larch | 49 | 4to60 | 20.5 | 20 | 17.01 | 23 | 17.35 |
| | | bl maple | 12 | 5to30 | 12.8 | 11 | 4.17 | 14.8 | 1.76 |
| | | spruce | 5 | 4to48 | 26 | 30 | 1.74 | 31.2 | 3.26 |
| | | vine maple | 10 | 3to6 | 3.9 | 4 | 3.47 | 4 | 0.11 |

| Twp-Range | 34-6e | species | # WTs | Diam Range | av Diam | Median Diam | rel Freq% | QMD | rel DOM% |
|---|---|---|---|---|---|---|---|---|---|
| # allWTs | 289 | red alder | 1 | 14 | 14 | 14 | 0.35 | 14 | 0.09 |
| all avDiam | 23.18 | redcedar | 28 | 12to72 | 35.4 | 30 | 9.69 | 40.9 | 22.31 |
| all QMD | 26.95 | Douglas-fir | 4 | 12to48 | 30 | 30 | 1.38 | 32.6 | 2.02 |
| SrvyYr | 1900 | grand fir | 1 | 5 | 5 | 5 | 0.35 | 5 | 0.01 |
| | | hemlock | 160 | 3to60 | 22 | 20 | 55.36 | 25.1 | 48.01 |
| | | larch | 82 | 6to60 | 21.2 | 18 | 28.37 | 24.5 | 23.44 |
| | | pine | 2 | 24to36 | 30 | 30 | 0.69 | 30.6 | 0.89 |
| | | spruce | 10 | 6to48 | 21.8 | 19 | 3.46 | 25.7 | 3.15 |
| | | yew | 1 | 6 | 6 | 6 | 0.35 | 6 | 0.02 |

| Twp-Range | 33-3e | species | # WTs | Diam Range | av Diam | Median Diam | rel Freq% | QMD | rel DOM% |
|---|---|---|---|---|---|---|---|---|---|
| # allWTs | 125 | red alder | 46 | 3to48 | 11.3 | 10 | 36.80 | 13.4 | 20.57 |
| all avDiam | 13.55 | redcedar | 10 | 4to50 | 19.7 | 15.5 | 8.00 | 24.4 | 14.83 |
| all QMD | 17.92 | crabapple | 8 | 4to11 | 6.5 | 6 | 6.40 | 7 | 0.98 |
| SrvyYr | 1872 | Douglas-fir | 7 | 4to72 | 28.1 | 30 | 5.60 | 35.7 | 22.22 |
| | | hemlock | 1 | 6 | 6 | 6 | 0.80 | 6 | 0.09 |
| | | juniper | 5 | 4to18 | 12.8 | 15 | 4.00 | 13.7 | 2.34 |
| | | spruce | 28 | 5to48 | 18.5 | 13 | 22.40 | 22.8 | 36.25 |
| | | willow | 18 | 3to14 | 6.8 | 6 | 14.40 | 7.5 | 2.52 |
| | | yew | 2 | 7to11 | 9 | 9 | 1.60 | 9.2 | 0.42 |

| Twp-Range | 33-4e | species | # WTs | Diam Range | av Diam | Median Diam | rel Freq% | QMD | rel DOM% |
|---|---|---|---|---|---|---|---|---|---|
| # allWTs | 279 | red alder | 45 | 3to32 | 12 | 10 | 16.13 | 14 | 4.56 |
| all avDiam | 20.8 | birch | 2 | 8 | 8 | 8 | 0.72 | 8 | 0.07 |
| all QMD | 26.3 | redcedar | 76 | 5to130 | 33.1 | 30 | 27.24 | 39.9 | 62.52 |
| SrvyYr | 1872 | crabapple | 2 | 8to11 | 9.5 | 9.5 | 0.72 | 9.6 | 0.10 |
| | | Douglas-fir | 13 | 5to60 | 24.2 | 20 | 4.66 | 28.9 | 5.61 |
| | | grand fir | 1 | 24 | 24 | 24 | 0.36 | 24 | 0.30 |
| | | hemlock | 105 | 5to36 | 17.2 | 18 | 37.63 | 18.8 | 19.18 |
| | | bl maple | 8 | 10to36 | 23.6 | 22 | 2.87 | 25.8 | 2.75 |
| | | spruce | 11 | 4to40 | 22.3 | 18 | 3.94 | 27.4 | 4.27 |
| | | vine maple | 1 | 8 | 8 | 8 | 0.36 | 8 | 0.03 |
| | | willow | 14 | 5to15 | 8.1 | 7 | 5.02 | 8.6 | 0.54 |
| | | yew | 1 | 10 | 10 | 10 | 0.36 | 10 | 0.05 |

| Twp-Range | 33-5e | species | # WTs | Diam Range | av Diam | Median Diam | rel Freq% | QMD | rel DOM% |
|---|---|---|---|---|---|---|---|---|---|
| # allWTs | 296 | red alder | 6 | 8to14 | 10.7 | 10 | 2.03 | 10.8 | 0.54 |
| all avDiam | 17.1 | redcedar | 74 | 5to60 | 28.3 | 30 | 25.00 | 31.5 | 56.30 |
| all QMD | 21.0 | Douglas-fir | 13 | 6to20 | 12.7 | 10 | 4.39 | 13.6 | 1.84 |
| SrvyYr | 1883 | grand fir | 1 | 30 | 30 | 30 | 0.34 | 30 | 0.69 |
| | &91 | hemlock | 169 | 5to60 | 13.7 | 12 | 57.09 | 16.1 | 33.59 |
| | | bigleaf map | 10 | 4to36 | 14.7 | 12 | 3.38 | 18 | 2.48 |
| | | spruce | 10 | 4to48 | 18.1 | 11 | 3.38 | 23 | 4.06 |
| | | vine maple | 10 | 3to6 | 4.2 | 4 | 3.38 | 4.3 | 0.14 |
| | | willow | 2 | 3 | 3 | 3 | 0.68 | 3 | 0.01 |
| | | yew | 1 | 10 | 10 | 10 | 0.34 | 10 | 0.08 |

| Twp-Range | 33-6e | species | # WTs | Diam Range | av Diam | Median Diam | rel Freq% | QMD | rel DOM% |
|---|---|---|---|---|---|---|---|---|---|
| # allWTs | 277 | red alder | 4 | 4to12 | 7.25 | 6 | 1.44 | 7.8 | 0.19 |
| all avDiam | 18.1 | redcedar | 21 | 5to60 | 26.2 | 6 | 7.58 | 33.9 | 18.60 |
| all QMD | 21.6 | Douglas-fir | 17 | 5to36 | 11.1 | 10 | 6.14 | 13.1 | 2.25 |
| SrvyYr | 1891 | grand fir | 1 | 20 | 20 | 20 | 0.36 | 20 | 0.31 |
| | | hemlock | 189 | 5to60 | 13.8 | 15 | 68.23 | 21.8 | 69.24 |
| | | larch | 4 | 6to12 | 10.3 | 10.5 | 1.44 | 10.7 | 0.35 |
| | | bigleaf map | 1 | 6 | 6 | 6 | 0.36 | 6 | 0.03 |
| | | pine | 1 | 6 | 6 | 6 | 0.36 | 6 | 0.03 |
| | | spruce | 39 | 6to36 | 15.3 | 14 | 14.08 | 17.4 | 9.10 |

| Twp-Range | 33-7e | species | # WTs | Diam Range | av Diam | Median Diam | rel Freq% | QMD | rel DOM% |
|---|---|---|---|---|---|---|---|---|---|
| # allWTs | 282 | red alder | 6 | 6to14 | 10.3 | 10.5 | 2.13 | 10.6 | 0.60 |
| all avDiam | 17.2 | redcedar | 4 | 16to28 | 21 | 20 | 1.42 | 21.5 | 1.66 |
| all QMD | 19.9 | Douglas-fir | 5 | 12to36 | 19 | 16 | 1.77 | 20.9 | 1.96 |
| SrvyYr | 1895 | grand fir | 94 | 4to48 | 18.6 | 14.5 | 33.33 | 21.6 | 39.27 |
| | | hemlock | 173 | 5to48 | 16.6 | 14 | 61.35 | 19.1 | 56.51 |

| Twp-Range | 32-4e | species | # WTs | Diam Range | av Diam | Median Diam | rel Freq% | QMD | rel DOM% |
|---|---|---|---|---|---|---|---|---|---|
| # allWTs | 253 | red alder | 25 | 3to24 | 9.2 | 8 | 9.88 | 10.3 | 1.37 |
| all avDiam | 22.9 | barberry | 1 | 3 | 3 | 3 | 0.40 | 3 | 0.00 |
| all QMD | 27.7 | redcedar | 78 | 8to80 | 26.1 | 22 | 30.83 | 29.9 | 35.95 |
| SrvyYr | 1872 | crabapple | 4 | 6to12 | 9 | 9 | 1.58 | 9.4 | 0.18 |
| | | cottonwood | 1 | 48 | 48 | 48 | 0.40 | 48 | 1.19 |
| | | Douglas-fir | 48 | 5to72 | 34.1 | 36 | 18.97 | 37.6 | 34.98 |
| | | hemlock | 66 | 4to40 | 17.5 | 18 | 26.09 | 19.3 | 12.67 |
| | | bl maple | 6 | 10to30 | 20 | 20 | 2.37 | 20.1 | 1.25 |
| | | pine | 1 | 3 | 3 | 3 | 0.40 | 3 | 0.00 |
| | | spruce | 15 | 7to80 | 32.9 | 24 | 5.93 | 39.5 | 12.06 |
| | | vine maple | 5 | 3to6 | 4.8 | 5 | 1.98 | 4.9 | 0.06 |
| | | willow | 3 | 5to6 | 5.3 | 5 | 1.19 | 5.4 | 0.05 |

| Twp-Range | 32-5e | species | # WTs | Diam Range | av Diam | Median Diam | rel Freq% | QMD | rel DOM% |
|---|---|---|---|---|---|---|---|---|---|
| # allWTs | 292 | red alder | 13 | 8to16 | 13 | 14 | 4.45 | 13.3 | 1.29 |
| all avDiam | 20.3 | barberry | 1 | 6 | 6 | 6 | 0.34 | 6 | 0.02 |
| all QMD | 24.7 | redcedar | 66 | 3to80 | 27.2 | 28 | 22.60 | 30.9 | 35.29 |
| SrvyYr | 1875 | cottonwood | 3 | 30to60 | 43.3 | 40 | 1.03 | 45.1 | 3.42 |
| | | dogwood | 2 | 9to12 | 10.5 | 10.5 | 0.68 | 10.6 | 0.13 |
| | | Douglas-fir | 45 | 4to96 | 29.4 | 25 | 15.41 | 34.5 | 29.99 |
| | | hemlock | 127 | 4to50 | 15.7 | 14 | 43.49 | 17.5 | 21.78 |
| | | bigleaf map | 5 | 10to30 | 17.6 | 14 | 1.71 | 18.9 | 1.00 |
| | | pine | 2 | 10to15 | 12.5 | 12.5 | 0.68 | 12.7 | 0.18 |
| | | spruce | 9 | 10to60 | 31.8 | 30 | 3.08 | 36.5 | 6.71 |
| | | vine maple | 18 | 3to5 | 4.4 | 4 | 6.16 | 4.1 | 0.17 |
| | | willow | 1 | 4 | 4 | 4 | 0.34 | 4 | 0.01 |

| Twp-Range | 32-6e | species | # WTs | Diam Range | av Diam | Median Diam | rel Freq% | QMD | rel DOM% |
|---|---|---|---|---|---|---|---|---|---|
| # allWTs | 273 | red alder | 9 | 6to11 | 8.4 | 9 | 3.30 | 8.7 | 0.62 |
| all avDiam | 15.4 | barberry | 1 | 6 | 6 | 6 | 0.37 | 6 | 0.03 |
| all QMD | 20.1 | redcedar | 33 | 4to96 | 30.7 | 28 | 12.09 | 37.1 | 41.07 |
| SrvyYr | 1890 | cherry | 1 | 6 | 6 | 6 | 0.37 | 6 | 0.03 |
| | | chittemwoo | 1 | 4 | 4 | 4 | 0.37 | 4 | 0.01 |
| | | cottonwood | 1 | 20 | 20 | 20 | 0.37 | 20 | 0.36 |
| | | dogwood | 1 | 5 | 5 | 5 | 0.37 | 5 | 0.02 |
| | | Douglas-fir | 18 | 6to60 | 27.4 | 27 | 6.59 | 32.4 | 17.08 |
| | | hemlock | 167 | 3to40 | 12.6 | 12 | 61.17 | 14.3 | 30.87 |
| | | larch | 5 | 12to30 | 16.4 | 14 | 1.83 | 17.8 | 1.43 |
| | | bl maple | 3 | 13to20 | 17 | 18 | 1.10 | 17.3 | 0.81 |
| | | pine | 2 | 8to12 | 10 | 10 | 0.73 | 15.2 | 0.42 |
| | | spruce | 6 | 7to60 | 27 | 18 | 2.20 | 33.7 | 6.16 |
| | | vine maple | 24 | 3to24 | 6.3 | 5 | 8.79 | 7.5 | 1.22 |
| | | willow | 1 | 5 | 5 | 5 | 0.37 | 5 | 0.02 |

| Twp-Range | 32-7e | species | # WTs | Diam Range | av Diam | Median Diam | rel Freq% | QMD | rel DOM% |
|---|---|---|---|---|---|---|---|---|---|
| # allWTs | 277 | red alder | 8 | 7to20 | 12.1 | 11.5 | 2.89 | 12.9 | 0.80 |
| all avDiam | 19.8 | barberry | 1 | 10 | 10 | 10 | 0.36 | 10 | 0.06 |
| all QMD | 24.5 | redcedar | 18 | 6to60 | 32.3 | 33 | 6.50 | 37 | 14.79 |
| SrvyYr | 1890& | Douglas-fir | 10 | 6to72 | 41.4 | 45 | 3.61 | 46.8 | 13.14 |
| | 1906 | hemlock | 193 | 4to70 | 18.8 | 14 | 69.68 | 22.8 | 60.20 |
| | | larch | 15 | 8to50 | 23.5 | 24 | 5.42 | 27.2 | 6.66 |
| | | bl maple | 9 | 6to24 | 13.9 | 12 | 3.25 | 15 | 1.22 |
| | | spruce | 9 | 8to36 | 20.8 | 20 | 3.25 | 22.7 | 2.78 |
| | | vine maple | 10 | 4to7 | 5.6 | 6 | 3.61 | 5.7 | 0.19 |
| | | yew | 4 | 6to9 | 7.3 | 7 | 1.44 | 7.4 | 0.13 |

| Twp-Range | 31-4e | species | # WTs | Diam Range | av Diam | Median Diam | rel Freq% | QMD | rel DOM% |
|---|---|---|---|---|---|---|---|---|---|
| # allWTs | 244 | red alder | 14 | 4to16 | 10.2 | 10 | 5.74 | 10.8 | 1.27 |
| all avDiam | 19.3 | redcedar | 69 | 2to72 | 22 | 20 | 28.28 | 25 | 33.50 |
| all QMD | 23.0 | cottonwood | 2 | 8to48 | 28 | 28 | 0.82 | 34.4 | 1.84 |
| SrvyYr | 1872 | dogwood | 1 | 8 | 8 | 8 | 0.41 | 8 | 0.05 |
| | | elder | 1 | 4 | 4 | 4 | 0.41 | 4 | 0.01 |
| | | Douglas-fir | 63 | 4to60 | 26.8 | 22 | 25.82 | 30.3 | 44.93 |
| | | hemlock | 73 | 4to30 | 14.5 | 14 | 29.92 | 15.7 | 13.98 |
| | | bl maple | 6 | 9to18 | 12.7 | 10.5 | 2.46 | 13.2 | 0.81 |
| | | spruce | 2 | 30to60 | 45 | 45 | 0.82 | 47.4 | 3.49 |
| | | vine maple | 7 | 3to7 | 4.7 | 4 | 2.87 | 5 | 0.14 |
| | | willow | 6 | 5to13 | 7.8 | 8 | 2.46 | 8.3 | 0.32 |

| Twp-Range | 31-5e | species | # WTs | Diam Range | av Diam | Median Diam | rel Freq% | QMD | rel DOM% |
|---|---|---|---|---|---|---|---|---|---|
| # allWTs | 264 | red alder | 32 | 3to20 | 10.2 | 10 | 12.12 | 11.1 | 4.88 |
| all avDiam | 13.1 | redcedar | 46 | 3to72 | 19.8 | 14 | 17.42 | 25.4 | 36.71 |
| all QMD | 17.5 | cherry | 3 | 2to14 | 8.7 | 10 | 1.14 | 10 | 0.37 |
| SrvyYr | | crabapple | 5 | 4to14 | 7.2 | 6 | 1.89 | 8.1 | 0.41 |
| | | cottonwood | 11 | 3to80 | 19.5 | 12 | 4.17 | 28.9 | 11.36 |
| | | Douglas-fir | 26 | 3to50 | 12.9 | 9 | 9.85 | 16.8 | 9.08 |
| | | hemlock | 81 | 3to60 | 12.3 | 10 | 30.68 | 14.4 | 20.77 |
| | | hazel | 2 | 5to8 | 6.5 | 6.5 | 0.76 | 6.7 | 0.11 |
| | | bl maple | 7 | 4to20 | 12.6 | 12 | 2.65 | 13.3 | 1.53 |
| | | pine | 8 | 4to16 | 8.6 | 7 | 3.03 | 9.6 | 0.91 |
| | | spruce | 13 | 10to60 | 23.3 | 20 | 4.92 | 28.8 | 13.34 |
| | | vine maple | 17 | 4to7 | 4.2 | 4 | 6.44 | 4.4 | 0.41 |
| | | willow | 13 | 3to8 | 4.6 | 4 | 4.92 | 5 | 0.40 |

| Twp-Range | 31-6e | species | # WTs | Diam Range | av Diam | Median Diam | rel Freq% | QMD | rel DOM% |
|---|---|---|---|---|---|---|---|---|---|
| # allWTs | 281 | red alder | 12 | 3to16 | 10 | 10 | 4.27 | 10.8 | 1.87 |
| all avDiam | 11.7 | barberry | 3 | 4to5 | 4.3 | 4 | 1.07 | 4.4 | 0.08 |
| all QMD | 16.3 | redcedar | 28 | 4to70 | 23.6 | 14 | 9.96 | 31.5 | 37.18 |
| SrvyYr | 1877 | cherry | 4 | 4to5 | 4.8 | 5 | 1.42 | 4.8 | 0.12 |
| | | cottonwood | 1 | 10 | 10 | 10 | 0.36 | 10 | 0.13 |
| | | dogwood | 1 | 5 | 5 | 5 | 0.36 | 5 | 0.03 |
| | | Douglas-fir | 132 | 3to60 | 11.1 | 6 | 46.98 | 15.1 | 40.27 |
| | | hemlock | 62 | 3to30 | 12.4 | 11 | 22.06 | 14 | 16.26 |
| | | bl maple | 10 | 3to16 | 7.7 | 5 | 3.56 | 9.1 | 1.11 |
| | | pine | 1 | 7 | 7 | 7 | 0.36 | 7 | 0.07 |
| | | spruce | 1 | 40 | 40 | 40 | 0.36 | 40 | 2.14 |
| | | vine maple | 19 | 3to4 | 3.4 | 3 | 6.76 | 3.5 | 0.31 |
| | | willow | 6 | 4to7 | 5 | 4.5 | 2.14 | 5.1 | 0.21 |
| | | yew | 1 | 5 | 5 | 5 | 0.36 | 5 | 0.03 |

| Twp-Range | 31-7e | species | # WTs | Diam Range | av Diam | Median Diam | rel Freq% | QMD | rel DOM% |
|---|---|---|---|---|---|---|---|---|---|
| # allWTs | 284 | red alder | 6 | 5to20 | 11.3 | 11 | 2.11 | 12.3 | 0.62 |
| all avDiam | 19.0 | redcedar | 22 | 10to120 | 33 | 19.5 | 7.75 | 44.5 | 29.68 |
| all QMD | 22.7 | Douglas-fir | 8 | 6to20 | 9.9 | 8.5 | 2.82 | 10.8 | 0.64 |
| SrvyYr | 1894 | grand fir | 38 | 7to36 | 15.9 | 13.5 | 13.38 | 17.3 | 7.75 |
| | | hemlock | 182 | 4to48 | 18.3 | 17 | 64.08 | 20.2 | 50.60 |
| | | larch | 7 | 16to36 | 27.4 | 30 | 2.46 | 28.5 | 3.87 |
| | | bl maple | 2 | 7to8 | 6.5 | 7.5 | 0.70 | 7.5 | 0.08 |
| | | spruce | 19 | 6to48 | 21.3 | 20 | 6.69 | 23.3 | 7.03 |

| Twp-Range | 30-1W& | species | # WTs | Diam Range | av Diam | Median Diam | rel Freq% | QMD | rel DOM% |
|---|---|---|---|---|---|---|---|---|---|
| # allWTs | 243 | red alder | 24 | 3to10 | 5.1 | 5 | 9.88 | 5.5 | 0.68 |
| ttl all avDiam | 17.4 | redcedar | 20 | 3to36 | 14.5 | 13.5 | 8.23 | 16.4 | 5.01 |
| ttl QMD | 21.0 | cherry | 1 | 6 | 6 | 6 | 0.41 | 6 | 0.03 |
| SrvyYr | 1858 | crabapple | 7 | 3to10 | 5 | 4 | 2.88 | 5.4 | 0.19 |
| | | cottonwood | 2 | 10to20 | 15 | 15 | 0.82 | 15.8 | 0.46 |
| | | Douglas-fir | 167 | 3to60 | 21.3 | 20 | 68.72 | 24.2 | 91.04 |
| | | hemlock | 11 | 6to18 | 9.9 | 10 | 4.53 | 10.4 | 1.11 |
| | | bl maple | 1 | 7 | 7 | 7 | 0.41 | 7 | 0.05 |
| | | spruce | 1 | 36 | 36 | 36 | 0.41 | 36 | 1.21 |
| | | willow | 9 | 3to60 | 4.2 | 4 | 3.70 | 4.3 | 0.15 |

| Twp-Range | 30-4e | species | # WTs | Diam Range | av Diam | Median Diam | rel Freq% | QMD | rel DOM% |
|---|---|---|---|---|---|---|---|---|---|
| # allWTs | 191 | red alder | 9 | 4to10 | 8.6 | 8 | 4.71 | 9.4 | 1.07 |
| all avDiam | 16.9 | redcedar | 69 | 4to40 | 18.2 | 16 | 36.13 | 20.2 | 37.84 |
| all QMD | 19.7 | dogwood | 1 | 8 | 8 | 8 | 0.52 | 8 | 0.09 |
| SrvyYr | 1859 | Douglas-fir | 57 | 5to60 | 21.5 | 18 | 29.84 | 25.2 | 48.64 |
| | | hemlock | 41 | 6to30 | 11.7 | 11 | 21.47 | 12.4 | 8.47 |
| | | bl maple | 11 | 4to18 | 12.2 | 12 | 5.76 | 12.8 | 2.42 |
| | | spruce | 1 | 12 | 12 | 12 | 0.52 | 12 | 0.19 |
| | | skunkwood | 1 | 12 | 12 | 12 | 0.52 | 12 | 0.19 |
| | | vine maple | 1 | 5 | 5 | 5 | 0.52 | 5 | 0.03 |

| Twp-Range | 30-5e | species | # WTs | Diam Range | av Diam | Median Diam | rel Freq% | QMD | rel DOM% |
|---|---|---|---|---|---|---|---|---|---|
| # allWTs | 252 | red alder | 2 | 8to10 | 9 | 9 | 0.79 | 9.1 | 0.16 |
| all avDiam | 15.3 | birch | 9 | 4to12 | 6.9 | 6 | 3.57 | 7.3 | 0.46 |
| all QMD | 20.2 | redcedar | 69 | 4to75 | 18.6 | 14 | 27.38 | 22.9 | 35.02 |
| SrvyYr | 1872 | cherry | 4 | 4 | 4 | 4 | 1.59 | 4 | 0.06 |
| | | cascara | 1 | 5 | 5 | 5 | 0.40 | 5 | 0.02 |
| | | crabapple | 1 | 6 | 6 | 6 | 0.40 | 6 | 0.03 |
| | | cottonwood | 1 | 14 | 14 | 14 | 0.40 | 14 | 0.19 |
| | | Douglas-fir | 68 | 4to70 | 20.8 | 18 | 26.98 | 26.5 | 46.22 |
| | | hemlock | 64 | 4to36 | 10 | 6 | 25.40 | 12.6 | 9.83 |
| | | bl maple | 12 | 3to32 | 13.2 | 9 | 4.76 | 16.5 | 3.16 |
| | | pine | 1 | 6 | 6 | 6 | 0.40 | 6 | 0.03 |
| | | spruce | 11 | 5to40 | 17.5 | 16 | 4.37 | 19.8 | 4.17 |
| | | vine maple | 1 | 6 | 6 | 6 | 0.40 | 6 | 0.03 |
| | | willow | 7 | 3to5 | 3.9 | 4 | 2.78 | 3.9 | 0.10 |
| | | yew | 1 | 4 | 4 | 4 | 0.40 | 4 | 0.02 |

| Twp-Range | 30-6e | species | # WTs | Diam Range | av Diam | Median Diam | rel Freq% | QMD | rel DOM% |
|---|---|---|---|---|---|---|---|---|---|
| # allWTs | 285 | red alder | 9 | 3to14 | 7.9 | 8 | 3.16 | 8.6 | 0.48 |
| all avDiam | 14.6 | bearb | 10 | 3to18 | 7.4 | 5 | 3.51 | 9 | 0.58 |
| QMD | 22.1 | redcedar | 63 | 3to84 | 31.7 | 24 | 22.11 | 39.3 | 70.07 |
| SrvyYr | 1881 | cherry | 3 | 4to6 | 4.7 | 4 | 1.05 | 4.8 | 0.05 |
| | | dogwood | 2 | 4to6 | 5 | 5 | 0.70 | 5 | 0.04 |
| | | Douglas-fir | 56 | 3to72 | 13.6 | 8 | 19.65 | 20.1 | 16.29 |
| | | hemlock | 87 | 3to35 | 10.5 | 8 | 30.53 | 12.6 | 9.95 |
| | | hazel | 5 | 3to4 | 3.4 | 3 | 1.75 | 3.4 | 0.04 |
| | | bigleaf map | 1 | 14 | 14 | 14 | 0.35 | 14 | 0.14 |
| | | madrone | 1 | 6 | 6 | 6 | 0.35 | 6 | 0.03 |
| | | pine | 2 | 6to12 | 9 | 9 | 0.70 | 9.5 | 0.13 |
| | | spruce | 5 | 5to36 | 19.4 | 20 | 1.75 | 23.1 | 1.92 |
| | | vine maple | 33 | 2to6 | 3.8 | 3 | 11.58 | 4 | 0.38 |
| | | willow | 7 | 3to6 | 3.9 | 3 | 2.46 | 4 | 0.08 |
| | | yew | 1 | 5 | 5 | 5 | 0.35 | 5 | 0.02 |

| Twp-Range | 30-7e | species | # WTs | Diam Range | av Diam | Median Diam | rel Freq% | QMD | rel DOM% |
|---|---|---|---|---|---|---|---|---|---|
| # allWTs | 278 | red alder | 8 | 3to20 | 10.4 | 9.5 | 2.88 | 11.6 | 1.24 |
| all avDiam | 13.1 | redcedar | 27 | 5to120 | 27.1 | 16 | 9.71 | 38.2 | 45.55 |
| all QMD | 17.6 | cherry | 1 | 4 | 4 | 4 | 0.36 | 4 | 0.02 |
| SrvyYr | 1890 | cottonwood | 1 | 22 | 22 | 22 | 0.36 | 22 | 0.56 |
| | | dogwood | 1 | 4 | 4 | 4 | 0.36 | 4 | 0.02 |
| | | Douglas-fir | 47 | 5to120 | 12.5 | 8 | 16.91 | 15.9 | 13.74 |
| | | grand fir | 1 | 10 | 10 | 10 | 0.36 | 10 | 0.12 |
| | | hemlock | 162 | 4to48 | 12.2 | 11 | 58.27 | 13.9 | 36.18 |
| | | larch | 8 | 6to30 | 12.5 | 11.5 | 2.88 | 14.3 | 1.89 |
| | | bigleaf map | 5 | 6to7 | 6.4 | 6 | 1.80 | 6.4 | 0.24 |
| | | spruce | 1 | 10 | 10 | 10 | 0.36 | 10 | 0.12 |
| | | vine maple | 10 | 3to8 | 4.6 | 4 | 3.60 | 4.8 | 0.27 |
| | | willow | 1 | 7 | 7 | 7 | 0.36 | 7 | 0.06 |
| | | yew | 5 | 6to11 | 7.6 | 7 | 1.80 | 7.8 | 0.35 |

| Twp-Range | 30-8e | species | # WTs | Diam Range | av Diam | Median Diam | rel Freq% | QMD | rel DOM% |
|---|---|---|---|---|---|---|---|---|---|
| # allWTs | 253 | red alder | 11 | 5to18 | 11.7 | 10 | 4.35 | 12.4 | 1.57 |
| all avDiam | 17.5 | redcedar | 4 | 10to120 | 41 | 10 | 1.58 | 61.8 | 14.19 |
| all QMD | 20.6 | cottonwood | 2 | 9to22 | 15.5 | 15.5 | 0.79 | 16.6 | 0.51 |
| SrvyYr | 1895 | Douglas-fir | 5 | 9to24 | 15.4 | 12 | 1.98 | 16.6 | 1.28 |
| | | grand fir | 53 | 6to48 | 17.8 | 14 | 20.95 | 20.7 | 21.10 |
| | | hemlock | 173 | 4to48 | 17.5 | 16 | 68.38 | 19.3 | 59.88 |
| | | larch | 1 | 9 | 9 | 9 | 0.40 | 9 | 0.08 |
| | | bl maple | 1 | 24 | 24 | 24 | 0.40 | 24 | 0.54 |
| | | pine | 1 | 22 | 22 | 22 | 0.40 | 22 | 0.45 |
| | | spruce | 2 | 5to10 | 7.5 | 7.5 | 0.79 | 7.9 | 0.12 |

| Twp-Range | 30-2w | species | # WTs | Diam Range | av Diam | Median Diam | rel Freq% | QMD | rel DOM% |
|---|---|---|---|---|---|---|---|---|---|
| # allWTs | 123 | red alder | 10 | 4to20 | 8.7 | 8 | 8.13 | 9.8 | 2.40 |
| all avDiam | 15.3 | redcedar | 15 | 6to36 | 15.8 | 14 | 12.20 | 18.1 | 12.27 |
| all QMD | 18.0 | Douglas-fir | 78 | 4to60 | 17.7 | 16 | 63.41 | 20.3 | 80.26 |
| SrvyYr | 1858 | hemlock | 15 | 5to24 | 9.3 | 8 | 12.20 | 10.2 | 3.90 |
| | | madrone | 1 | 8 | 8 | 8 | 0.81 | 8 | 0.16 |
| | | pine | 1 | 12 | 12 | 12 | 0.81 | 12 | 0.36 |
| | | spruce | 1 | 8 | 8 | 8 | 0.81 | 8 | 0.16 |
| | | willow | 2 | 4 | 4 | 4 | 1.63 | 4 | 0.08 |

| Twp-Range | 30-3w | species | # WTs | Diam Range | av Diam | Median Diam | rel Freq% | QMD | rel DOM% |
|---|---|---|---|---|---|---|---|---|---|
| # allWTs | 181 | red alder | 15 | 4to30 | 96 | 8 | 8.29 | 11.5 | 2.67 |
| all avDiam | 17.1 | redcedar | 19 | 6to50 | 16.1 | 12 | 10.50 | 18.9 | 9.15 |
| all QMD | 20.2 | cottonwood | 2 | 8to16 | 12 | 12 | 1.10 | 12.6 | 0.43 |
| SrvyYr | 1858 | Douglas-fir | 123 | 3to60 | 18.8 | 18 | 67.96 | 21.6 | 77.36 |
| | | hemlock | 4 | 4to18 | 10.5 | 10 | 2.21 | 11.7 | 0.74 |
| | | bl maple | 5 | 4to30 | 19.2 | 24 | 2.76 | 21.8 | 3.20 |
| | | oak | 5 | 8to24 | 14.8 | 12 | 2.76 | 15.8 | 1.68 |
| | | spruce | 1 | 60 | 60 | 60 | 0.55 | 60 | 4.85 |
| | | willow | 7 | 3to8 | 4.6 | 4 | 3.87 | 5 | 0.24 |

| Twp-Range | 30-4w | species | # WTs | Diam Range | av Diam | Median Diam | rel Freq% | QMD | rel DOM% |
|---|---|---|---|---|---|---|---|---|---|
| # allWTs | 277 | red alder | 22 | 5to24 | 9.9 | 8 | 7.94 | 11.1 | 2.38 |
| all avDiam | 15.6 | redcedar | 46 | 5to80 | 26.9 | 22 | 16.61 | 32.4 | 42.48 |
| all QMD | 20.3 | crabapple | 4 | 6to10 | 7.8 | 7.5 | 1.44 | 7.9 | 0.22 |
| SrvyYr | 1858 | cottonwood | 2 | 8to15 | 11.5 | 11.5 | 0.72 | 12 | 0.25 |
| | | Douglas-fir | 124 | 3to55 | 16.2 | 12 | 44.77 | 20.2 | 44.51 |
| | | hemlock | 58 | 4to30 | 11.4 | 10 | 20.94 | 13.1 | 8.76 |
| | | bl maple | 4 | 8to24 | 14.8 | 13.5 | 1.44 | 15.9 | 0.89 |
| | | willow | 17 | 3to10 | 5.1 | 4 | 6.14 | 5.5 | 0.45 |

| Twp-Range | 30-5w | species | # WTs | Diam Range | av Diam | Median Diam | rel Freq% | QMD | rel DOM% |
|---|---|---|---|---|---|---|---|---|---|
| # allWTs | 236 | red alder | 52 | 2to18 | 7.1 | 6 | 22.03 | 8 | 6.25 |
| all avDiam | 11.8 | redcedar | 35 | 6to48 | 15 | 12 | 14.83 | 17.3 | 19.66 |
| all QMD | 15.0 | cherry | 1 | 10 | 10 | 10 | 0.42 | 10 | 0.19 |
| SrvyYr | 1862 | Douglas-fir | 107 | 3to55 | 15.4 | 12 | 45.34 | 18.7 | 70.23 |
| | | hemlock | 4 | 8to18 | 11.5 | 10 | 1.69 | 12.2 | 1.12 |
| | | bl maple | 8 | 4to14 | 9 | 9 | 3.39 | 9.6 | 1.38 |
| | | willow | 29 | 1to10 | 3.9 | 4 | 12.29 | 4.3 | 1.01 |

| Twp-Range | 30-6w | species | # WTs | Diam Range | av Diam | Median Diam | rel Freq% | QMD | rel DOM% |
|---|---|---|---|---|---|---|---|---|---|
| # allWTs | 229 | red alder | 27 | 2to20 | 8.9 | 8 | 11.79 | 10 | 5.46 |
| all avDiam | 11.3 | aspen | 2 | 5 | 5 | 5 | 0.87 | 5 | 0.10 |
| all QMD | 14.7 | redcedar | 29 | 3to60 | 20.7 | 18 | 12.66 | 24.4 | 34.92 |
| SrvyYr | 1879 | cherry | 2 | 8to24 | 16 | 16 | 0.87 | 17.9 | 1.30 |
| | | crabapple | 2 | 3 | 3 | 3 | 0.87 | 3 | 0.04 |
| | | Douglas-fir | 82 | 2to40 | 11.1 | 7.5 | 35.81 | 14.6 | 35.36 |
| | | hemlock | 65 | 3to36 | 10.4 | 9 | 28.38 | 12.4 | 20.22 |
| | | bl maple | 6 | 3to30 | 9.7 | 6.5 | 2.62 | 13.4 | 2.18 |
| | | willow | 14 | 2to7 | 3.3 | 3 | 6.11 | 3.6 | 0.37 |

| Twp-Range | 29-1e | species | # WTs | Diam Range | av Diam | Median Diam | rel Freq% | QMD | rel DOM% |
|---|---|---|---|---|---|---|---|---|---|
| # allWTs | 72 | red alder | 3 | 6to8 | 7.3 | 8 | 4.17 | 7.4 | 0.42 |
| all avDiam | 17.9 | redcedar | 17 | 6to72 | 25.1 | 24 | 23.61 | 30.7 | 40.83 |
| all QMD | 23.3 | Douglas-fir | 20 | 4to66 | 20.8 | 13 | 27.78 | 26.9 | 36.88 |
| SrvyYr | 1858 | grand fir | 1 | 40 | 40 | 40 | 1.39 | 40 | 4.08 |
| | | hemlock | 24 | 5to36 | 12.7 | 11 | 33.33 | 14.6 | 13.04 |
| | | bl maple | 2 | 12to18 | 15 | 15 | 2.78 | 15.3 | 1.19 |
| | | spruce | 1 | 36 | 36 | 36 | 1.39 | 36 | 3.30 |
| | | willow | 4 | 3to4 | 3.5 | 3.5 | 5.56 | 3.5 | 0.12 |

| Twp-Range | 29-5e | species | # WTs | Diam Range | av Diam | Median Diam | rel Freq% | QMD | rel DOM% |
|---|---|---|---|---|---|---|---|---|---|
| # allWTs | 197 | red alder | 10 | 3to28 | 10.6 | 8 | 5.08 | 12.8 | 1.43 |
| all avDiam | 16.6 | redcedar | 35 | 5to120 | 31.9 | 18 | 17.77 | 44.7 | 61.12 |
| all QMD | 24.1 | cherry | 2 | 7 | 7 | 7 | 1.02 | 7 | 0.09 |
| SrvyYr | 1869 | cascara | 4 | 2to6 | 5 | 5 | 2.03 | 5 | 0.09 |
| | | crabapple | 7 | 3to11 | 5.6 | 5 | 3.55 | 6.1 | 0.23 |
| | | Douglas-fir | 59 | 2to130 | 13.4 | 11 | 29.95 | 16.3 | 13.70 |
| | | hemlock | 31 | 3to45 | 14.9 | 12 | 15.74 | 17.3 | 8.11 |
| | | juniper | 5 | 3to7 | 4.8 | 5 | 2.54 | 5.1 | 0.11 |
| | | bl maple | 5 | 6to20 | 15 | 16 | 2.54 | 15.9 | 1.10 |
| | | pine | 18 | 3to10 | 6.1 | 6 | 9.14 | 6.5 | 0.66 |
| | | spruce | 18 | 10to48 | 26.2 | 27 | 9.14 | 29.1 | 13.32 |
| | | willow | 1 | 1 | 7 | 7 | 0.51 | 7 | 0.04 |
| | | yew | 1 | 14 | 14 | 14 | 0.51 | 14 | 0.17 |

| Twp-Range | 29-6e | species | # WTs | Diam Range | av Diam | Median Diam | rel Freq% | QMD | rel DOM% |
|---|---|---|---|---|---|---|---|---|---|
| # allWTs | 263 | red alder | 14 | 5to50 | 13.2 | 9.5 | 5.32 | 17.3 | 6.61 |
| all avDiam | 11.1 | barberry | 3 | 3to9 | 6.3 | 7 | 1.14 | 6.8 | 0.22 |
| all QMD | 15.5 | redcedar | 46 | 3to80 | 16.5 | 12 | 17.49 | 23.9 | 41.48 |
| SrvyYr | 1878 | cherry | 5 | 5to8 | 6.4 | 6 | 1.90 | 6.5 | 0.33 |
| | | crabapple | 4 | 2to6 | 4 | 4 | 1.52 | 4.3 | 0.12 |
| | | cottonwood | 3 | 3to50 | 18.7 | 3 | 1.14 | 30 | 4.26 |
| | | Douglas-fir | 82 | 3to48 | 10.7 | 8 | 31.18 | 14.1 | 25.73 |
| | | hemlock | 66 | 4to30 | 10.1 | 9 | 25.10 | 11.6 | 14.02 |
| | | hazel | 4 | 3to48 | 3.3 | 3 | 1.52 | 3.3 | 0.07 |
| | | bl maple | 18 | 3to24 | 9 | 6 | 6.84 | 11.1 | 3.50 |
| | | spruce | 4 | 6to40 | 18 | 13 | 1.52 | 22.3 | 3.14 |
| | | skookum | 1 | 9 | 9 | 9 | 0.38 | 9 | 0.13 |
| | | vine maple | 12 | 2to6 | 4 | 4 | 4.56 | 4.2 | 0.33 |
| | | willow | 1 | 8 | 8 | 8 | 0.38 | 8 | 0.10 |

| Twp-Range | 29-7e | species | # WTs | Diam Range | av Diam | Median Diam | rel Freq% | QMD | rel DOM% |
|---|---|---|---|---|---|---|---|---|---|
| # allWTs | 280 | red alder | 10 | 2to9 | 5.2 | 4.5 | 3.57 | 5.6 | 0.67 |
| all avDiam | 10.4 | barberry | 4 | 5to6 | 5.5 | 5.5 | 1.43 | 5.5 | 0.26 |
| all QMD | 12.9 | redcedar | 26 | 4to48 | 23.3 | 19 | 9.29 | 26.2 | 38.07 |
| SrvyYr | 1892 | cherry | 1 | 6 | 6 | 6 | 0.36 | 6 | 0.08 |
| | | crabapple | 2 | 4to7 | 5.5 | 5.5 | 0.71 | 5.5 | 0.13 |
| | | dogwood | 2 | 5to6 | 5.5 | 5.5 | 0.71 | 5.5 | 0.13 |
| | | Douglas-fir | 28 | 4to48 | 11.9 | 10 | 10.00 | 14.5 | 12.56 |
| | | grand fir | 7 | 4to8 | 6.1 | 6 | 2.50 | 6.3 | 0.59 |
| | | hemlock | 168 | 3to30 | 9.7 | 8 | 60.00 | 11.1 | 44.15 |
| | | larch | 1 | 11 | 11 | 11 | 0.36 | 11 | 0.26 |
| | | bl maple | 9 | 3to24 | 8.4 | 5 | 3.21 | 10.9 | 2.28 |
| | | spruce | 2 | 8to15 | 11.5 | 11.5 | 0.71 | 12 | 0.61 |
| | | vine maple | 17 | 3to5 | 3.9 | 4 | 6.07 | 4 | 0.58 |
| | | willow | 2 | 3to4 | 3.5 | 3.5 | 0.71 | 3.5 | 0.05 |
| | | yew | 1 | 7 | 7 | 7 | 0.36 | 7 | 0.10 |

| Twp-Range | 29-8e | species | # WTs | Diam Range | av Diam | Median Diam | rel Freq% | QMD | rel DOM% |
|---|---|---|---|---|---|---|---|---|---|
| # allWTs | 208 | red alder | 1 | 16 | 16 | 16 | 0.48 | 16 | 0.35 |
| all avDiam | 16.8 | redcedar | 14 | 7to36 | 21 | 19 | 6.73 | 22.7 | 9.76 |
| all QMD | 18.9 | Douglas-fir | 5 | 13to30 | 20.4 | 18 | 2.40 | 21.4 | 3.10 |
| SrvyYr | 1894 | grand fir | 44 | 5to36 | 14.8 | 13 | 21.15 | 16.2 | 15.62 |
| | | hemlock | 141 | 4to48 | 17.1 | 15 | 67.79 | 19.3 | 71.02 |
| | | larch | 3 | 7to11 | 8.7 | 8 | 1.44 | 8.8 | 0.31 |

| Twp-Range | 29-1w | species | # WTs | Diam Range | av Diam | Median Diam | rel Freq% | QMD | rel DOM% |
|---|---|---|---|---|---|---|---|---|---|
| # allWTs | 252 | red alder | 41 | 4to14 | 6.8 | 6 | 16.27 | 7.2 | 2.52 |
| all avDiam | 14.8 | redcedar | 51 | 6to60 | 22.1 | 18 | 20.24 | 26.1 | 41.22 |
| all QMD | 18.3 | cherry | 2 | 6to8 | 7 | 7 | 0.79 | 7.1 | 0.12 |
| SrvyYr | 1862 | crabapple | 6 | 4to8 | 6 | 6 | 2.38 | 6.1 | 0.26 |
| | | Douglas-fir | 89 | 3to50 | 17.2 | 12 | 35.32 | 20.2 | 43.09 |
| | | hemlock | 46 | 5to24 | 11.3 | 10 | 18.25 | 12.8 | 8.94 |
| | | bl maple | 5 | 11 | 10 | | 1.98 | 11.7 | 0.81 |
| | | pine | 2 | 6to18 | 12 | 12 | 0.79 | 13.4 | 0.43 |
| | | spruce | 6 | 12to28 | 17.3 | 16 | 2.38 | 18.3 | 2.38 |
| | | willow | 3 | 4to8 | 6 | 6 | 1.19 | 6.2 | 0.14 |
| | | yew | 1 | 10 | 10 | 10 | 0.40 | 10 | 0.12 |

| Twp-Range | 29-2w | species | # WTs | Diam Range | av Diam | Median Diam | rel Freq% | QMD | rel DOM% |
|---|---|---|---|---|---|---|---|---|---|
| # allWTs | 287 | red alder | 11 | 4to24 | 14.2 | 14 | 3.83 | 15.8 | 3.01 |
| all avDiam | 14.9 | redcedar | 66 | 4to96 | 17.7 | 12 | 23.00 | 23 | 38.22 |
| all QMD | 17.8 | cherry | 3 | 5to7 | 6 | 6 | 1.05 | 6.1 | 0.12 |
| SrvyYr | 1873 | Douglas-fir | 45 | 3to48 | 14.8 | 14 | 15.68 | 17.3 | 14.74 |
| | | grand fir | 1 | 40 | 40 | 40 | 0.35 | 40 | 1.75 |
| | | hemlock | 155 | 4to45 | 14.2 | 14 | 54.01 | 15.8 | 42.36 |
| | | bl maple | 4 | 8to24 | 13 | 12 | 1.39 | 14.6 | 0.93 |
| | | vine maple | 1 | 6 | 6 | 6 | 0.35 | 6 | 0.04 |
| | | yew | 1 | 10 | 10 | 10 | 0.35 | 10 | 0.11 |

| Twp-Range | 29-3w | species | # WTs | Diam Range | av Diam | Median Diam | rel Freq% | QMD | rel DOM% |
|---|---|---|---|---|---|---|---|---|---|
| # allWTs | 212 | red alder | 4 | 4to30 | 12.5 | 8 | 1.89 | 16.2 | 1.56 |
| all avDiam | 15.1 | redcedar | 75 | 5to50 | 16.1 | 13 | 35.38 | 18.2 | 36.88 |
| all QMD | 17.8 | Douglas-fir | 99 | 4to55 | 15.3 | 11.5 | 46.70 | 18.5 | 50.29 |
| SrvyYr | 1891 | hemlock | 32 | 4to24 | 11.7 | 11 | 15.09 | 13.4 | 8.53 |
| | | bl maple | 2 | 29 | 29 | 29 | 0.94 | 29 | 2.50 |

| Twp-Range | 29-4w | species | # WTs | Diam Range | av Diam | Median Diam | rel Freq% | QMD | rel DOM% |
|---|---|---|---|---|---|---|---|---|---|
| # allWTs | 170 | red alder | 9 | 4to24 | 9.3 | 8 | 5.29 | 11 | 1.63 |
| all avDiam | 16.8 | redcedar | 50 | 5to48 | 18.9 | 15 | 29.41 | 21 | 32.97 |
| all QMD | 19.8 | cherry | 3 | 3to10 | 5.3 | 3 | 1.76 | 6.3 | 0.18 |
| SrvyYr | 1893 | Douglas-fir | 73 | 3to48 | 18.7 | 18 | 42.94 | 22.2 | 53.80 |
| | | hemlock | 32 | 4to30 | 13.4 | 11 | 18.82 | 15.4 | 11.35 |
| | | bl maple | 1 | 6 | 6 | 6 | 0.59 | 6 | 0.05 |
| | | madrone | 1 | 5 | 5 | 5 | 0.59 | 5 | 0.04 |
| | | willow | 1 | 4 | 4 | 4 | 0.59 | 4 | 0.02 |

| Twp-Range | 29-5w | species | # WTs | Diam Range | av Diam | Median Diam | rel Freq% | QMD | rel DOM% |
|---|---|---|---|---|---|---|---|---|---|
| # allWTs | 182 | red alder | 7 | 6to10 | 7.6 | 7 | 3.85 | 7.8 | 0.66 |
| all avDiam | 16.1 | redcedar | 17 | 8to36 | 17.8 | 14 | 9.34 | 19.6 | 10.14 |
| all QMD | 18.8 | Douglas-fir | 68 | 4to72 | 18.2 | 14 | 37.36 | 21.9 | 50.62 |
| SrvyYr | 1897 | grand fir | 6 | 8to16 | 10 | 9 | 3.30 | 10.4 | 1.01 |
| | | hemlock | 84 | 5to36 | 15.1 | 12 | 46.15 | 17 | 37.68 |

| Twp-Range | 28-1e | species | # WTs | Diam Range | av Diam | Median Diam | rel Freq% | QMD | rel DOM% |
|---|---|---|---|---|---|---|---|---|---|
| # allWTs | 179 | red alder | 15 | 4to15 | 7.9 | 8 | 8.52 | 8.4 | 1.24 |
| all avDiam | 17.8 | redcedar | 59 | 4to70 | 23.9 | 24 | 33.52 | 28.5 | 55.98 |
| all QMD | 22.1 | Douglas-fir | 28 | 6to70 | 20.5 | 14.5 | 15.91 | 25.6 | 21.43 |
| SrvyYr | 1859 | hemlock | 68 | 3to36 | 14.3 | 12 | 38.64 | 15.9 | 20.08 |
| | | pine | 1 | 24 | 24 | 24 | 0.57 | 24 | 0.67 |
| | | spruce | 2 | 12to40 | 26 | 26 | 1.14 | 29.5 | 2.03 |
| | | willow | 3 | 6to9 | 7.7 | 8 | 1.70 | 7.8 | 0.21 |

| Twp-Range | 28-2e | species | # WTs | Diam Range | av Diam | Median Diam | rel Freq% | QMD | rel DOM% |
|---|---|---|---|---|---|---|---|---|---|
| # allWTs | 89 | red alder | 10 | 4to8 | 5.8 | 5.5 | 11.24 | 5.9 | 0.77 |
| all avDiam | 18.3 | redcedar | 27 | 6to48 | 14.4 | 18 | 30.34 | 24.6 | 35.97 |
| all QMD | 22.6 | Douglas-fir | 23 | 4to70 | 25.2 | 20 | 25.84 | 31 | 48.66 |
| SrvyYr | 1859 | hemlock | 27 | 6to30 | 14.1 | 12 | 30.34 | 15.5 | 14.28 |
| | | willow | 2 | 6to10 | 8 | 8 | 2.25 | 8.2 | 0.30 |

| Twp-Range | 28-4e | species | # WTs | Diam Range | av Diam | Median Diam | rel Freq% | QMD | rel DOM% |
|---|---|---|---|---|---|---|---|---|---|
| # allWTs | 200 | red alder | 7 | 4to12 | 7.9 | 6 | 3.50 | 8.5 | 0.64 |
| all avDiam | 16.6 | redcedar | 67 | 4to 60 | 21.7 | 16 | 33.50 | 25.8 | 56.12 |
| all QMD | 19.9 | Douglas-fir | 18 | 4to48 | 24.1 | 20 | 9.00 | 26.5 | 15.91 |
| SrvyYr | 1860 | hemlock | 102 | 4to36 | 12.7 | 12 | 51.00 | 14.2 | 25.88 |
| | | bl maple | 3 | 12to18 | 15.3 | 16 | 1.50 | 15.5 | 0.91 |
| | | pine | 1 | 8 | 8 | 8 | 0.50 | 8 | 0.08 |
| | | spruce | 1 | 14 | 14 | 14 | 0.50 | 14 | 0.25 |
| | | willow | 1 | 6 | 6 | 6 | 0.50 | 6 | 0.05 |

| Twp-Range | 28-5e | species | # WTs | Diam Range | av Diam | Median Diam | rel Freq% | QMD | rel DOM% |
|---|---|---|---|---|---|---|---|---|---|
| # allWTs | 253 | red alder | 24 | 4to25 | 14.9 | 12 | 9.49 | 17 | 5.63 |
| all avDiam | 19.0 | redcedar | 31 | 6to60 | 25.3 | 24 | 12.25 | 28.3 | 20.15 |
| all QMD | 22.1 | crabapple | 2 | 4to6 | 5 | 5 | 0.79 | 5 | 0.04 |
| SrvyYr | 1871 | Douglas-fir | 50 | 4to72 | 27.2 | 28 | 19.76 | 30.2 | 37.00 |
| | | hemlock | 97 | 2to40 | 17.4 | 15 | 38.34 | 19.4 | 29.62 |
| | | bl maple | 6 | 12to20 | 16.8 | 17.5 | 2.37 | 17.1 | 1.42 |
| | | pine | 24 | 5to20 | 9.6 | 8 | 9.49 | 10.4 | 2.11 |
| | | spruce | 12 | 8to28 | 17.3 | 16 | 4.74 | 18.6 | 3.37 |
| | | sknkwd | 1 | 8 | 8 | 8 | 0.40 | 8 | 0.05 |
| | | vine maple | 1 | 6 | 6 | 6 | 0.40 | 6 | 0.03 |
| | | willow | 4 | 6to12 | 8.25 | 7 | 1.58 | 8.6 | 0.24 |
| | | yew | 1 | 10 | 10 | 10 | 0.40 | 10 | 0.08 |

| Twp-Range | 28-6e | species | # WTs | Diam Range | av Diam | Median Diam | rel Freq% | QMD | rel DOM% |
|---|---|---|---|---|---|---|---|---|---|
| # allWTs | 286 | red alder | 26 | 3to24 | 7.9 | 8.5 | 9.09 | 9.2 | 3.42 |
| all avDiam | 9.7 | barberry | 7 | 3to8 | 5.1 | 5 | 2.45 | 5.5 | 0.33 |
| all QMD | 15.0 | redcedar | 51 | 3to85 | 23.1 | 20 | 17.83 | 28.3 | 63.56 |
| SrvyYr | 1866 | cherry | 1 | 3 | 3 | 3 | 0.35 | 3 | 0.01 |
| | | crabapple | 13 | 3to10 | 3.6 | 3 | 4.55 | 4.1 | 0.34 |
| | | cottonwood | 2 | 16to46 | 31 | 31 | 0.70 | 34.4 | 3.68 |
| | | dogwood | 2 | 3to6 | 4.5 | 4.5 | 0.70 | 4.7 | 0.07 |
| | | Douglas-fir | 57 | 2to48 | 8.6 | 4 | 19.93 | 13.6 | 16.41 |
| | | hemlock | 28 | 3to20 | 9 | 8 | 9.79 | 10 | 4.36 |
| | | hazel | 3 | 3 | 3 | 3 | 1.05 | 3 | 0.04 |
| | | bl maple | 8 | 3to15 | 6.6 | 4 | 2.80 | 8 | 0.80 |
| | | pine | 4 | 3to12 | 6 | 4.5 | 1.40 | 7 | 0.30 |
| | | spruce | 11 | 3to30 | 13.6 | 3 | 3.85 | 16.9 | 4.89 |
| | | vine maple | 49 | 3to6 | 3.5 | 3 | 17.13 | 3.6 | 0.99 |
| | | willow | 22 | 3to10 | 3.9 | 3 | 7.69 | 4.4 | 0.66 |
| | | yew | 2 | 3to4 | 3.5 | 3.5 | 0.70 | 3.5 | 0.04 |

| Twp-Range | 28-7e | species | # WTs | Diam Range | av Diam | Median Diam | rel Freq% | QMD | rel DOM% |
|---|---|---|---|---|---|---|---|---|---|
| # allWTs | 285 | red alder | 9 | 3to18 | 5.8 | 5 | 3.16 | 7.3 | 0.64 |
| all avDiam | 11.1 | redcedar | 38 | 3to48 | 25.4 | 18 | 13.33 | 32.5 | 53.17 |
| all QMD | 16.3 | cherry | 1 | 3 | 3 | 3 | 0.35 | 3 | 0.01 |
| SrvyYr | 1873 | dogwood | 2 | 3to6 | 4.5 | 4.5 | 0.70 | 4.7 | 0.06 |
| | | Douglas-fir | 28 | 3to60 | 10.7 | 5 | 9.82 | 17.9 | 11.88 |
| | | grand fir | 8 | 4to20 | 7.6 | 6 | 2.81 | 9 | 0.86 |
| | | hemlock | 146 | 3to50 | 9.7 | 8 | 51.23 | 11.9 | 27.39 |
| | | spruce | 1 | 16 | 16 | 16 | 0.35 | 16 | 0.34 |
| | | bl maple | 39 | 3to30 | 7.3 | 4 | 13.68 | 9.8 | 4.96 |
| | | vine maple | 12 | 3to48 | 3.8 | 4 | 4.21 | 3.9 | 0.24 |
| | | willow | 1 | 2 | 2 | 2 | 0.35 | 2 | 0.01 |

| Twp-Range | 28-8e | species | # WTs | Diam Range | av Diam | Median Diam | rel Freq% | QMD | rel DOM% |
|---|---|---|---|---|---|---|---|---|---|
| # allWTs | 269 | red alder | 23 | 3to20 | 8.2 | 7 | 8.55 | 8.9 | 1.66 |
| all avDiam | 15.6 | barberry | 5 | 4to8 | 6 | 6 | 1.86 | 6.2 | 0.17 |
| all QMD | 20.2 | redcedar | 20 | 4to96 | 23.2 | 15 | 7.43 | 38.7 | 27.21 |
| SrvyYr | 1883&91 | Douglas-fir | 35 | 6to54 | 24.5 | 20 | 13.01 | 28.9 | 26.56 |
| | | hemlock | 148 | 2to54 | 14.4 | 12 | 55.02 | 16.9 | 38.40 |
| | | larch | 4 | 8to22 | 14.5 | 14 | 1.49 | 15.4 | 0.86 |
| | | bl maple | 9 | 4to30 | 18 | 14 | 3.35 | 22.8 | 4.25 |
| | | spruce | 2 | 20to30 | 25 | 25 | 0.74 | 25.5 | 1.18 |
| | | vine maple | 17 | 3to6 | 4.2 | 4 | 6.32 | 4.2 | 0.27 |
| | | willow | 4 | 3to6 | 4.5 | 3 | 1.49 | 4.2 | 0.06 |
| | | yew | 2 | 3to4 | 3.5 | 3.5 | 0.74 | 3.5 | 0.02 |

| Twp-Range | 28-1w | species | # WTs | Diam Range | av Diam | Median Diam | rel Freq% | QMD | rel DOM% |
|---|---|---|---|---|---|---|---|---|---|
| # allWTs | 301 | red alder | 10 | 5to10 | 7.5 | 7 | 3.32 | 7.7 | 0.65 |
| all avDiam | 15.2 | redcedar | 80 | 4to50 | 17.3 | 14.5 | 26.58 | 20 | 35.12 |
| all QMD | 17.4 | cherry | 1 | 8 | 8 | 8 | 0.33 | 8 | 0.07 |
| SrvyYr | 1873 | Douglas-fir | 89 | 3to48 | 15.6 | 14 | 29.57 | 18 | 31.65 |
| | | grand fir | 1 | 16 | 16 | 16 | 0.33 | 16 | 0.28 |
| | | hemlock | 112 | 4to30 | 14.1 | 13 | 37.21 | 15.3 | 28.77 |
| | | bl maple | 1 | 11 | 11 | 11 | 0.33 | 11 | 0.13 |
| | | pine | 2 | 12to16 | 14 | 14 | 0.66 | 14.1 | 0.44 |
| | | spruce | 5 | 10to40 | 20.8 | 20 | 1.66 | 23.4 | 3.00 |

| Twp-Range | 28-2w | species | # WTs | Diam Range | av Diam | Median Diam | rel Freq% | QMD | rel DOM% |
|---|---|---|---|---|---|---|---|---|---|
| # allWTs | 214 | red alder | 15 | 5to24 | 10.7 | 10 | 7.01 | 11.8 | 3.11 |
| all avDiam | 15.2 | redcedar | 26 | 8to72 | 20.6 | 17 | 12.15 | 24.6 | 23.46 |
| all QMD | 17.7 | cherry | 1 | 8 | 8 | 8 | 0.47 | 8 | 0.10 |
| SrvyYr | 1873 | crabapple | 1 | 5 | 5 | 5 | 0.47 | 5 | 0.04 |
| | | Douglas-fir | 59 | 6to46 | 18.6 | 18 | 27.57 | 20.9 | 38.43 |
| | | grand fir | 1 | 16 | 16 | 16 | 0.47 | 16 | 0.38 |
| | | hemlock | 99 | 4to32 | 13.4 | 12 | 46.26 | 14.6 | 31.47 |
| | | bl maple | 9 | 3to24 | 6.7 | 4 | 4.21 | 9.2 | 1.14 |
| | | spruce | 1 | 36 | 36 | 36 | 0.47 | 36 | 1.93 |
| | | willow | 2 | 3 | 3 | 3 | 0.93 | 3 | 0.03 |

| Twp-Range | 27-1e | species | # WTs | Diam Range | av Diam | Median Diam | rel Freq% | QMD | rel DOM% |
|---|---|---|---|---|---|---|---|---|---|
| # allWTs | 153 | red alder | 18 | 4to12 | 8 | 7.5 | 11.76 | 8.6 | 2.32 |
| all avDiam | 16.4 | redcedar | 33 | 6to50 | 20.8 | 18 | 21.57 | 23.8 | 32.59 |
| all QMD | 19.4 | Douglas-fir | 46 | 3to98 | 20.5 | 18 | 30.07 | 23 | 42.42 |
| SrvyYr | 1859 | hemlock | 48 | 6to26 | 12.8 | 11 | 31.37 | 14 | 16.40 |
| | | spruce | 6 | 5to40 | 17.8 | 8 | 3.92 | 23.8 | 5.92 |
| | | willow | 2 | 6to8 | 7 | 7 | 1.31 | 7.1 | 0.18 |

| Twp-Range | 27-2e | species | # WTs | Diam Range | av Diam | Median Diam | rel Freq% | QMD | rel DOM% |
|---|---|---|---|---|---|---|---|---|---|
| # allWTs | 222 | red alder | 18 | 4to15 | 8.5 | 8 | 8.11 | 9.2 | 1.71 |
| all avDiam | 16.2 | redcedar | 62 | 6to70 | 19.5 | 13.5 | 27.93 | 24.1 | 40.33 |
| all QMD | 20.1 | Douglas-fir | 50 | 3to60 | 22 | 20 | 22.52 | 26.4 | 39.03 |
| SrvyYr | 1859 | hemlock | 85 | 6to30 | 12 | 10 | 38.29 | 13.1 | 16.34 |
| | | bl maple | 5 | 12to24 | 20 | 20 | 2.25 | 20.5 | 2.35 |
| | | spruce | 2 | 6 | 6 | 6 | 0.90 | 6 | 0.08 |

| Twp-Range | 27-4e | species | # WTs | Diam Range | av Diam | Median Diam | rel Freq% | QMD | rel DOM% |
|---|---|---|---|---|---|---|---|---|---|
| # allWTs | 278 | red alder | 24 | 3to18 | 12.7 | 12 | 8.63 | 13.6 | 4.65 |
| all avDiam | 15.5 | redcedar | 78 | 6to60 | 21.5 | 16 | 28.06 | 25.7 | 53.96 |
| all QMD | 18.5 | Douglas-fir | 52 | 5to48 | 15.1 | 12 | 18.71 | 17.8 | 17.26 |
| SrvyYr | 1859 | hemlock | 114 | 6to36 | 12.4 | 12 | 41.01 | 13.6 | 22.08 |
| | | bl maple | 5 | 6to24 | 14.2 | 15 | 1.80 | 15.4 | 1.24 |
| | | pine | 2 | 10to24 | 9 | 9 | 0.72 | 9.1 | 0.17 |
| | | spruce | 1 | 24 | 24 | 24 | 0.36 | 24 | 0.60 |
| | | vine maple | 2 | 4to10 | 7 | 7 | 0.72 | 7.6 | 0.12 |

| Twp-Range | 27-5e | species | # WTs | Diam Range | av Diam | Median Diam | rel Freq% | QMD | rel DOM% |
|---|---|---|---|---|---|---|---|---|---|
| # allWTs | 287 | red alder | 9 | 8to18 | 10.1 | 9 | 3.14 | 10.9 | 1.11 |
| all avDiam | 14.1 | redcedar | 75 | 4to60 | 17.3 | 12 | 26.13 | 21.7 | 36.54 |
| ttl QMD | 18.3 | cherry | 1 | 5 | 5 | 5 | 0.35 | 5 | 0.03 |
| SrvyYr | 1874 | crabapple | 1 | 5 | 5 | 5 | 0.35 | 5 | 0.03 |
| | | dogwood | 2 | 5to8 | 6.5 | 6.5 | 0.70 | 6.7 | 0.09 |
| | | Douglas-fir | 68 | 3to60 | 16.1 | 10 | 23.69 | 21.3 | 31.92 |
| | | hemlock | 113 | 3to50 | 11.9 | 9 | 39.37 | 14.9 | 25.96 |
| | | bl maple | 5 | 3to36 | 16.8 | 10 | 1.74 | 20.6 | 2.20 |
| | | pine | 1 | 5 | 5 | 5 | 0.35 | 5 | 0.03 |
| | | spruce | 3 | 5to36 | 22.7 | 24 | 1.05 | 25.4 | 2.00 |
| | | vine maple | 8 | 3to5 | 4.1 | 4 | 2.79 | 4.2 | 0.15 |
| | | willow | 1 | 6 | 6 | 6 | 0.35 | 6 | 0.04 |

| Twp-Range | 27-6e | species | # WTs | Diam Range | av Diam | Median Diam | rel Freq% | QMD | rel DOM% |
|---|---|---|---|---|---|---|---|---|---|
| # allWTs | 268 | red alder | 18 | 4to18 | 11.8 | 12 | 6.72 | 12.7 | 2.89 |
| all avDiam | 14.4 | redcedar | 45 | 6to100 | 21.1 | 14 | 16.79 | 29.5 | 39.04 |
| all QMD | 19.3 | cascara | 10 | 4to14 | 8.7 | 10 | 3.73 | 9.1 | 0.83 |
| SrvyYr | 1871 | cottonwood | 3 | 8to36 | 24.7 | 30 | 1.12 | 27.4 | 2.25 |
| | | crabappale | 6 | 3to6 | 4.5 | 4.5 | 2.24 | 4.7 | 0.13 |
| | | Douglas-fir | 59 | 3to60 | 19.7 | 16 | 22.01 | 23.5 | 32.48 |
| | | hemlock | 65 | 4to30 | 10.5 | 8 | 24.25 | 12.5 | 10.12 |
| | | bl maple | 36 | 3to48 | 13.3 | 10 | 13.43 | 17.2 | 10.62 |
| | | pine | 1 | 5 | 5 | 5 | 0.37 | 5 | 0.02 |
| | | spruce | 8 | 5to10 | 8.1 | 10 | 2.99 | 8.5 | 0.58 |
| | | vine maple | 8 | 4to10 | 6.3 | 4 | 2.99 | 6.9 | 0.38 |
| | | willow | 8 | 3to11 | 6.3 | 6 | 2.99 | 7 | 0.39 |
| | | yew | 1 | 18 | 18 | 18 | 0.37 | 18 | 0.32 |

| Twp-Range | 27-7e | species | # WTs | Diam Range | av Diam | Median Diam | rel Freq% | QMD | rel DOM% |
|---|---|---|---|---|---|---|---|---|---|
| # allWTs | 286 | red alder | 11 | 3to24 | 13.14 | 12 | 3.85 | 14.1 | 2.07 |
| all avDiam | 14.0 | redcedar | 48 | 4to80 | 25.7 | 24 | 16.78 | 31.3 | 44.53 |
| ttl QMD | 19.2 | cottonwood | 7 | 5to24 | 12.3 | 13 | 2.45 | 13.6 | 1.23 |
| SrvyYr | 1873 | dogwood | 2 | 4 | 4 | 4 | 0.70 | 4 | 0.03 |
| | | Douglas-fir | 28 | 3to70 | 21.6 | 14 | 9.79 | 28.7 | 21.84 |
| | | grand fir | 1 | 20 | 20 | 20 | 0.35 | 20 | 0.38 |
| | | hemlock | 127 | 3to40 | 10.9 | 8 | 44.41 | 13.2 | 20.95 |
| | | hazel | 1 | 4 | 4 | 4 | 0.35 | 4 | 0.02 |
| | | bl maple | 21 | 3to30 | 9.1 | 4 | 7.34 | 12 | 2.86 |
| | | spruce | 5 | 7to50 | 23.4 | 20 | 1.75 | 28.1 | 3.74 |
| | | cascara | 4 | 6to24 | 10.5 | 6 | 1.40 | 13.1 | 0.65 |
| | | vine maple | 28 | 3to30 | 5.5 | 4 | 9.79 | 7.4 | 1.45 |
| | | willow | 2 | 4to10 | 7 | 7 | 0.70 | 7.6 | 0.11 |
| | | yew | 1 | 12 | 12 | 12 | 0.35 | 12 | 0.14 |

| Twp-Range | 27-1w | species | # WTs | Diam Range | av Diam | Median Diam | rel Freq% | QMD | rel DOM% |
|---|---|---|---|---|---|---|---|---|---|
| # allWTs | 223 | red alder | 20 | 3to15 | 6.9 | 6 | 8.97 | 7.7 | 1.00 |
| all avDiam | 18.3 | redcedar | 60 | 8to90 | 24.5 | 24 | 26.91 | 28.6 | 41.39 |
| all QMD | 23.1 | cherry | 4 | 4to6 | 4.8 | 4.5 | 1.79 | 4.8 | 0.08 |
| SrvyYr | 1859 | Douglas-fir | 77 | 3to60 | 23.2 | 24 | 34.53 | 27.6 | 49.47 |
| | | hemlock | 35 | 4to24 | 12.1 | 12 | 15.70 | 13.2 | 5.14 |
| | | bl maple | 10 | 3to20 | 9.9 | 6.5 | 4.48 | 11.9 | 1.19 |
| | | pine | 1 | 12 | 12 | 12 | 0.45 | 12 | 0.12 |
| | | spruce | 4 | 15to24 | 20.8 | 22 | 1.79 | 21.1 | 1.50 |
| | | vine maple | 3 | 4 | 4 | 4 | 1.35 | 4 | 0.04 |
| | | willow | 9 | 3to5 | 3.9 | 4 | 4.04 | 3.9 | 0.12 |

| Twp-Range | 27-2w | species | # WTs | Diam Range | av Diam | Median Diam | rel Freq% | QMD | rel DOM% |
|---|---|---|---|---|---|---|---|---|---|
| # allWTs | 161 | red alder | 17 | 4to20 | 10.2 | 10 | 10.56 | 11.2 | 4.78 |
| all avDiam | 12.9 | redcedar | 13 | 9to48 | 23.8 | 18 | 8.07 | 26.7 | 20.76 |
| all QMD | 16.7 | cherry | 3 | 5to6 | 5.3 | 5 | 1.86 | 5.6 | 0.21 |
| SrvyYr | 1877 | crabapple | 1 | 6 | 6 | | 0.62 | 6 | 0.08 |
| | | Douglas-fir | 91 | 3to60 | 14 | 10 | 56.52 | 18.2 | 67.52 |
| | | hemlock | 27 | 4to14 | 7.5 | 7 | 16.77 | 8 | 3.87 |
| | | bl maple | 8 | 3to20 | 8.3 | 6 | 4.97 | 9.7 | 1.69 |
| | | spruce | 1 | 20 | 20 | 20 | 0.62 | 20 | 0.90 |

| Twp-Range | 26-1e | species | # WTs | Diam Range | av Diam | Median Diam | rel Freq% | QMD | rel DOM% |
|---|---|---|---|---|---|---|---|---|---|
| # allWTs | 241 | red alder | 20 | 4to12 | 7.2 | 6 | 8.30 | 7.9 | 2.11 |
| all avDiam | 13.3 | redcedar | 58 | 4to36 | 14.3 | 11 | 24.07 | 15.7 | 24.22 |
| all QMD | 15.7 | cherry | 1 | 4 | 4 | 4 | 0.41 | 4 | 0.03 |
| SrvyYr | 1859 | Douglas-fir | 74 | 3to50 | 16.7 | 14 | 30.71 | 19.9 | 49.65 |
| | | grand fir | 1 | 38 | 38 | 38 | 0.41 | 38 | 2.45 |
| | | hemlock | 70 | 3to26 | 10.2 | 10 | 29.05 | 11.3 | 15.14 |
| | | bl maple | 2 | 8to20 | 14 | 14 | 0.83 | 15.2 | 0.78 |
| | | madrone | 1 | 16 | 16 | 16 | 0.41 | 16 | 0.43 |
| | | pine | 14 | 6to26 | 14.1 | 12 | 5.81 | 15.7 | 5.85 |

| Twp-Range | 26-2e | species | # WTs | Diam Range | av Diam | Median Diam | rel Freq% | QMD | rel DOM% |
|---|---|---|---|---|---|---|---|---|---|
| # allWTs | 157 | red alder | 13 | 4to10 | 5.3 | 4 | 8.28 | 5.7 | 0.75 |
| all avDiam | 15.0 | redcedar | 44 | 3to50 | 17.6 | 15 | 28.03 | 20.6 | 33.08 |
| all QMD | 19.0 | Douglas-fir | 45 | 4to50 | 19.8 | 12 | 28.66 | 24.5 | 47.86 |
| SrvyYr | 1859 | hemlock | 50 | 4to36 | 11.4 | 8 | 31.85 | 13.7 | 16.63 |
| | | madrone | 3 | 4to18 | 9 | 5 | 1.91 | 11 | 0.64 |
| | | pine | 1 | 12 | 12 | 12 | 0.64 | 12 | 0.26 |
| | | spruce | 1 | 16 | 16 | 16 | 0.64 | 16 | 0.45 |

| Twp-Range | 26-4e | species | # WTs | Diam Range | av Diam | Median Diam | rel Freq% | QMD | rel DOM% |
|---|---|---|---|---|---|---|---|---|---|
| # allWTs | 239 | red alder | 16 | 3to14 | 6.6 | 5.5 | 6.69 | 7.4 | 0.71 |
| all avDiam | 17.7 | ash | 2 | 5to8 | 6.5 | 6.5 | 0.84 | 6.7 | 0.07 |
| all QMD | 22.7 | redcedar | 64 | 4to72 | 21.5 | 17 | 26.78 | 25.7 | 34.46 |
| SrvyYr | 1859 | cherry | 2 | 3to4 | 3.5 | 3.5 | 0.84 | 3.5 | 0.02 |
| | | crabapple | 2 | 4to9 | 6.5 | 6.5 | 0.84 | 7 | 0.08 |
| | | cottonwood | 2 | 12to16 | 14 | 14 | 0.84 | 14 | 0.32 |
| | | Douglas-fir | 102 | 3to72 | 22.1 | 15.5 | 42.68 | 26.7 | 59.28 |
| | | hemlock | 34 | 3to18 | 9.5 | 8.5 | 14.23 | 10.4 | 3.00 |
| | | bl maple | 6 | 3to40 | 13.8 | 10 | 2.51 | 18.8 | 1.73 |
| | | pine | 2 | 4 | 4 | 4 | 0.84 | 4 | 0.03 |
| | | vine maple | 4 | 3to4 | 3.5 | 3.5 | 1.67 | 3.5 | 0.04 |
| | | willow | 2 | 3to6 | 4.5 | 4.5 | 0.84 | 4.7 | 0.04 |
| | | yew | 1 | 4 | 4 | 4 | 0.42 | 4 | 0.01 |

| Twp-Range | 26-5e | species | # WTs | Diam Range | av Diam | Median Diam | rel Freq% | QMD | rel DOM% |
|---|---|---|---|---|---|---|---|---|---|
| # allWTs | 284 | alder | 13 | 4to12 | 7.3 | 7 | 4.58 | 7.7 | 0.51 |
| all avDiam | 18.2 | ash | 11 | 6to24 | 11.9 | 9 | 3.87 | 13.6 | 1.34 |
| ttl QMD | 23.1 | redcedar | 86 | 4to86 | 23.2 | 20 | 30.28 | 27.2 | 41.98 |
| SrvyYr | 1870 | cherry | 3 | 5 | 5 | 5 | 1.06 | 5 | 0.05 |
| | | crabapple | 10 | 3to10 | 5.7 | 5.5 | 3.52 | 5.9 | 0.23 |
| | | Douglas-fir | 103 | 3to90 | 21.1 | 18 | 36.27 | 26.4 | 47.37 |
| | | grand fir | 2 | 6to8 | 7 | 7 | 0.70 | 7.1 | 0.07 |
| | | hemlock | 34 | 3to46 | 12.6 | 10 | 11.97 | 15.1 | 5.12 |
| | | BL maple | 16 | 4to30 | 11.7 | 10 | 5.63 | 13.7 | 1.98 |
| | | spruce | 2 | 24to36 | 30 | 30 | 0.70 | 30.6 | 1.24 |
| | | willow | 3 | 4to8 | 5.3 | 4 | 1.06 | 5.7 | 0.06 |
| | | yew | 1 | 14 | 14 | 14 | 0.35 | 14 | 0.13 |

| Twp-Range | 26-6e | species | # WTs | Diam Range | av Diam | Median Diam | rel Freq% | QMD | rel DOM% |
|---|---|---|---|---|---|---|---|---|---|
| # allWTs | 297 | red alder | 20 | 2to36 | 11.3 | 8.5 | 6.73 | 14.4 | 3.29 |
| all avDiam | 15.8 | redcedar | 63 | 3to50 | 20.5 | 16 | 21.21 | 23.9 | 28.54 |
| all QMD | 20.6 | cherry | 1 | 4 | 4 | 4 | 0.34 | 4 | 0.01 |
| SrvyYr | 1873 | cottonwood | 1 | 44 | 44 | 44 | 0.34 | 44 | 1.54 |
| | | crabapple | 6 | 3to10 | 6 | 5.5 | 2.02 | 6.5 | 0.20 |
| | | dogwood | 1 | 2 | 2 | 2 | 0.34 | 2 | 0.00 |
| | | Douglas-fir | 91 | 3to80 | 21.4 | 22 | 30.64 | 25.9 | 48.42 |
| | | hemlock | 54 | 2to36 | 10.8 | 8 | 18.18 | 13.8 | 8.16 |
| | | hazel | 2 | 2to36 | 2.5 | 2.5 | 0.67 | 2.5 | 0.01 |
| | | bl maple | 20 | 2to48 | 16.5 | 10 | 6.73 | 22.1 | 7.75 |
| | | spruce | 11 | 4to34 | 11.5 | 10 | 3.70 | 13.9 | 1.69 |
| | | cascara | 1 | 12 | 12 | 12 | 0.34 | 12 | 0.11 |
| | | vine maple | 21 | 2to7 | 4.4 | 4 | 7.07 | 4.7 | 0.37 |
| | | willow | 5 | 2to36 | 2.4 | 2 | 1.68 | 6 | 0.14 |

| Twp-Range | 26-7e | species | # WTs | Diam Range | av Diam | Median Diam | rel Freq% | QMD | rel DOM% |
|---|---|---|---|---|---|---|---|---|---|
| # allWTs | 290 | red alder | 9 | 2to32 | 12.7 | 10 | 3.10 | 15.8 | 1.43 |
| all avDiam | 18.4 | barberry | 2 | 5to6 | 5.5 | 5.5 | 0.69 | 5.5 | 0.04 |
| all QMD | 23.3 | redcedar | 67 | 4to72 | 30.9 | 30 | 23.10 | 34.8 | 51.55 |
| SrvyYr | 1885 | cherry | 1 | 7 | 7 | 7 | 0.34 | 7 | 0.03 |
| | | cascara | 1 | 10 | 10 | 10 | 0.34 | 10 | 0.06 |
| | | cottonwood | 1 | 45 | 45 | 45 | 0.34 | 45 | 1.29 |
| | | crabapple | 5 | 4to12 | 6.6 | 5 | 1.72 | 7.3 | 0.17 |
| | | dogwood | 2 | 5to6 | 6.5 | 6.5 | 0.69 | 6.7 | 0.06 |
| | | Douglas-fir | 40 | 4to60 | 20.4 | 14.5 | 13.79 | 25.7 | 16.78 |
| | | hemlock | 139 | 4to50 | 14.1 | 10 | 47.93 | 17.1 | 25.82 |
| | | bl maple | 12 | 6to30 | 16.1 | 14 | 4.14 | 18.4 | 2.58 |
| | | spruce | 1 | 10 | 10 | 10 | 0.34 | 10 | 0.06 |
| | | vine maple | 7 | 4to6 | 4.7 | 4 | 2.41 | 4.8 | 0.10 |
| | | willow | 2 | 4to50 | 4.5 | 4.5 | 0.69 | 4.5 | 0.03 |
| | | yew | 1 | 9 | 9 | 9 | 0.34 | 9 | 0.05 |

| Twp-Range | 26-8e | species | # WTs | Diam Range | av Diam | Median Diam | rel Freq% | QMD | rel DOM% |
|---|---|---|---|---|---|---|---|---|---|
| # allWTs | 286 | red alder | 2 | 12to14 | 13 | 13 | 0.70 | 13 | 0.13 |
| all avDiam | 25.75 | barberry | 1 | 5 | 5 | 5 | 0.35 | 5 | 0.01 |
| all QMD | 30.31 | redcedar | 39 | 10to84 | 39.9 | 36 | 13.64 | 43.9 | 28.60 |
| SrvyYr | 1895 | Douglas-fir | 54 | 4to108 | 27.7 | 20 | 18.88 | 34.5 | 24.46 |
| | | hemlock | 172 | 4to60 | 23.8 | 20.5 | 60.14 | 25.3 | 41.89 |
| | | bl maple | 5 | 8to26 | 13.4 | 10 | 1.75 | 15 | 0.43 |
| | | spruce | 12 | 6to72 | 23.8 | 13 | 4.20 | 31.5 | 4.53 |
| | | yew | 1 | 6 | 6 | 6 | 0.35 | 6 | 0.01 |

| Twp-Range | 26-1w | species | # WTs | Diam Range | av Diam | Median Diam | rel Freq% | QMD | rel DOM% |
|---|---|---|---|---|---|---|---|---|---|
| # allWTs | 119 | red alder | 9 | 4to14 | 7.3 | 6 | 7.56 | 8 | 1.00 |
| all avDiam | 18.5 | redcedar | 39 | 8to70 | 23.4 | 20 | 32.77 | 26.8 | 48.73 |
| all QMD | 22.0 | Douglas-fir | 34 | 3to40 | 23.1 | 24 | 28.57 | 25.6 | 38.76 |
| SrvyYr | 1859 | grand fir | 1 | 12 | 12 | 12 | 0.84 | 12 | 0.25 |
| | | hemlock | 29 | 3to30 | 11.8 | 12 | 24.37 | 13 | 8.53 |
| | | hazel | 1 | 4 | 4 | | 0.84 | 4 | 0.03 |
| | | bl maple | 3 | 4to20 | 14.7 | 20 | 2.52 | 16.5 | 1.42 |
| | | madrone | 1 | 4 | 4 | 4 | 0.84 | 4 | 0.03 |
| | | pine | 1 | 24 | 24 | 24 | 0.84 | 24 | 1.00 |
| | | willow | 1 | 10 | 10 | 10 | 0.84 | 10 | 0.17 |

| Twp-Range | 26-2w | species | # WTs | Diam Range | av Diam | Median Diam | rel Freq% | QMD | rel DOM% |
|---|---|---|---|---|---|---|---|---|---|
| # allWTs | 69 | red alder | 3 | 10to12 | 10.7 | 10 | 4.35 | 10.7 | 0.95 |
| all avDiam | 19.3 | redcedar | 8 | 10to48 | 23.5 | 20 | 11.59 | 26.1 | 15.06 |
| all QMD | 22.9 | Douglas-fir | 34 | 5to70 | 22.8 | 19 | 49.28 | 27.1 | 69.00 |
| SrvyYr | 1871 | hemlock | 21 | 6to30 | 14.2 | 14 | 30.43 | 15.2 | 13.41 |
| | | bl maple | 2 | 10to14 | 12 | 12 | 2.90 | 12 | 0.80 |
| | | spruce | 1 | 14 | 14 | 14 | 1.45 | 14 | 0.54 |

| Twp-Range | 25-1e | species | # WTs | Diam Range | av Diam | Median Diam | rel Freq% | QMD | rel DOM% |
|---|---|---|---|---|---|---|---|---|---|
| # allWTs | 239 | red alder | 42 | 3to16 | 5 | 4 | 17.57 | 5.5 | 2.12 |
| all avDiam | 12.4 | redcedar | 16 | 4to20 | 10.3 | 9 | 6.69 | 11.2 | 3.35 |
| all QMD | 15.8 | dogwood | 2 | 5to6 | 5.5 | 5.5 | 0.84 | 5.5 | 0.10 |
| SrvyYr | 1858 | Douglas-fir | 141 | 2to60 | 16.4 | 14 | 59.00 | 19.5 | 89.54 |
| | | hemlock | 21 | 4to24 | 8.9 | 8 | 8.79 | 10 | 3.51 |
| | | bl maple | 2 | 4 | 4 | 4 | 0.84 | 4 | 0.05 |
| | | madrone | 3 | 3to10 | 4 | 4 | 1.26 | 4 | 0.08 |
| | | pine | 5 | 6to10 | 8.8 | 10 | 2.09 | 8.9 | 0.66 |
| | | skunkwood | 2 | 5to6 | 5.5 | 5.5 | 0.84 | 5.5 | 0.10 |
| | | willow | 5 | 3to4 | 3.2 | 3 | 2.09 | 3.2 | 0.09 |

| Twp-Range | 25-3e | species | # WTs | Diam Range | av Diam | Median Diam | rel Freq% | QMD | rel DOM% |
|---|---|---|---|---|---|---|---|---|---|
| # allWTs | 94 | red alder | 13 | 6to70 | 16.1 | 12 | 13.83 | 22.7 | 14.31 |
| all avDiam | 19.2 | ash | 1 | 10 | 10 | 10 | 1.06 | 10 | 0.21 |
| all QMD | 22.3 | barberry | 1 | 7 | 7 | 7 | 1.06 | 7 | 0.10 |
| SrvyYr | 1855 | redcedar | 22 | 9to40 | 24.9 | 24 | 23.40 | 26.6 | 33.25 |
| | | Douglas-fir | 44 | 7to48 | 20 | 18 | 46.81 | 22.4 | 47.16 |
| | | hemlock | 5 | 4to9 | 7.2 | 8 | 5.32 | 7.5 | 0.60 |
| | | bl maple | 7 | 10to24 | 15.7 | 18 | 7.45 | 16.6 | 4.12 |
| | | willow | 1 | 4 | 4 | 4 | 1.06 | 4 | 0.03 |

| Twp-Range | 25-4e | species | # WTs | Diam Range | av Diam | Median Diam | rel Freq% | QMD | rel DOM% |
|---|---|---|---|---|---|---|---|---|---|
| # allWTs | 164 | red alder | 11 | 6to24 | 12.6 | 12 | 6.71 | 13.7 | 3.35 |
| all avDiam | 16.1 | ash | 7 | 2to20 | 9 | 10 | 4.27 | 11.2 | 1.42 |
| all QMD | 19.4 | barberry | 2 | 3to15 | 11.5 | 11.5 | 1.22 | 14.3 | 0.66 |
| SrvyYr | 1855 | redcedar | 44 | 6to40 | 19.7 | 18 | 26.83 | 22 | 34.55 |
| | | crabapple | 1 | 5 | 5 | 5 | 0.61 | 5 | 0.04 |
| | | cottonwood | 3 | 10to20 | 16.7 | 20 | 1.83 | 17.3 | 1.46 |
| | | Douglas-fir | 74 | 2to50 | 17.2 | 14 | 45.12 | 21.2 | 53.95 |
| | | grand fir | 1 | 15 | 15 | 15 | 0.61 | 15 | 0.36 |
| | | hemlock | 14 | 2to18 | 8.2 | 8 | 8.54 | 9.2 | 1.92 |
| | | bl maple | 6 | 5to20 | 13.8 | 15 | 3.66 | 14.8 | 2.13 |
| | | willow | 1 | 10 | 10 | 10 | 0.61 | 10 | 0.16 |

| Twp-Range | 25-5e | species | # WTs | Diam Range | av Diam | Median Diam | rel Freq% | QMD | rel DOM% |
|---|---|---|---|---|---|---|---|---|---|
| # allWTs | 256 | red alder | 17 | 2to20 | 7.8 | 7 | 6.64 | 9.2 | 1.39 |
| all avDiam | 17.0 | ash | 1 | 16 | 16 | 16 | 0.39 | 16 | 0.25 |
| all QMD | 20.1 | redecedar | 82 | 5to60 | 20.9 | 20 | 32.03 | 24.1 | 46.00 |
| SrvyYr | 1870 | cottonwood | 3 | 16to20 | 17.3 | 16 | 1.17 | 17.4 | 0.88 |
| | | crabapple | 4 | 5to10 | 6.5 | 5.5 | 1.56 | 6.8 | 0.18 |
| | | Douglas-fir | 111 | 4to54 | 17.7 | 18 | 43.36 | 20.5 | 45.05 |
| | | hemlock | 18 | 4to30 | 12.1 | 10.5 | 7.03 | 13.5 | 3.17 |
| | | bl maple | 16 | 6to20 | 12.1 | 11 | 6.25 | 12.8 | 2.53 |
| | | pine | 1 | 12 | 12 | 12 | 0.39 | 12 | 0.14 |
| | | vine maple | 1 | 6 | 6 | 6 | 0.39 | 6 | 0.03 |
| | | willow | 2 | 6 | 6 | 6 | 0.78 | 6 | 0.07 |

| Twp-Range | 25-6e | species | # WTs | Diam Range | av Diam | Median Diam | rel Freq% | QMD | rel DOM% |
|---|---|---|---|---|---|---|---|---|---|
| # allWTs | 243 | red alder | 24 | 2to36 | 8.3 | 6 | 9.88 | 10.8 | 3.14 |
| all avDiam | 15.5 | ash | 1 | 8 | 8 | 8 | 0.41 | 8 | 0.07 |
| all QMD | 19.1 | redcedar | 46 | 3to48 | 19.7 | 16 | 18.93 | 22.8 | 26.86 |
| SrvyYr | 1874 | chittemwoo | 1 | 3 | 3 | 3 | 0.41 | 3 | 0.01 |
| | | cottonwood | 1 | 44 | 44 | 44 | 0.41 | 44 | 2.17 |
| | | dogwood | 1 | 10 | 10 | 10 | 0.41 | 10 | 0.11 |
| | | Douglas-fir | 100 | 3to48 | 17.7 | 14 | 41.15 | 21.2 | 50.48 |
| | | hemlock | 48 | 4to 36 | 11.7 | 11 | 19.75 | 14.3 | 11.02 |
| | | bl maple | 20 | 3to 42 | 13.1 | 11 | 8.23 | 17.2 | 6.65 |
| | | willow | 1 | 5 | 5 | 5 | 0.41 | 5 | 0.03 |

| Twp-Range | 25-7e | species | # WTs | Diam Range | av Diam | Median Diam | rel Freq% | QMD | rel DOM% |
|---|---|---|---|---|---|---|---|---|---|
| # allWTs | 279 | red alder | 24 | 2to30 | 9.4 | 6.5 | 8.60 | 11.9 | 4.57 |
| all avDiam | 12.2 | barberry | 6 | 3to10 | 6.5 | 6 | 2.15 | 7 | 0.40 |
| all QMD | 16.3 | redcedar | 56 | 3to60 | 14.8 | 12 | 20.07 | 17.5 | 23.08 |
| SrvyYr | 1873 | cherry | 1 | 4 | 4 | 4 | 0.36 | 4 | 0.02 |
| | | cascara | 2 | 5to10 | 7.6 | 7.5 | 0.72 | 7.9 | 0.17 |
| | | crabapple | 5 | 3to6 | 3.8 | 3 | 1.79 | 4 | 0.11 |
| | | cottonwood | 7 | 4to45 | 18.1 | 15 | 2.51 | 21.9 | 4.52 |
| | | dogwood | 1 | 3 | 3 | 3 | 0.36 | 3 | 0.01 |
| | | Douglas-fir | 43 | 3to48 | 13.3 | 10 | 15.41 | 16.9 | 16.52 |
| | | hemlock | 90 | 3to36 | 11.4 | 8 | 32.26 | 14 | 23.73 |
| | | hazel | 1 | 3 | 3 | | 3.00 | | 0.00 |
| | | bl maple | 14 | 3to72 | 23 | 13 | 5.02 | 32.5 | 19.90 |
| | | spruce | 2 | 10to60 | 35 | 35 | 0.72 | 43 | 4.98 |
| | | vine maple | 24 | 3to30 | 5.6 | 4 | 8.60 | 8.2 | 2.17 |
| | | willow | 2 | 3to5 | 4 | 4 | 0.72 | 4.1 | 0.05 |
| | | yew | 1 | 7 | 7 | 7 | 0.36 | 7 | 0.07 |

| Twp-Range | 25-8e | species | # WTs | Diam Range | av Diam | Median Diam | rel Freq% | QMD | rel DOM% |
|---|---|---|---|---|---|---|---|---|---|
| # allWTs | 279 | red alder | 5 | 5to36 | 17 | 18 | 1.79 | 20.2 | 1.26 |
| all avDiam | 19.9 | barberry | 1 | 4 | 4 | 4 | 0.36 | 4 | 0.01 |
| all QMD | 24.1 | redcedar | 45 | 5to96 | 31.3 | 28 | 16.13 | 36.5 | 37.09 |
| SrvyYr | 1891 | Douglas-fir | 16 | 3to100 | 32.4 | 29 | 5.73 | 40.2 | 16.00 |
| | | hemlock | 194 | 3to40 | 17 | 18 | 69.53 | 18.9 | 42.88 |
| | | bl maple | 6 | 8to30 | 14.2 | 12 | 2.15 | 16 | 0.95 |
| | | spruce | 6 | 6to28 | 19 | 22 | 2.15 | 21 | 1.64 |
| | | vine maple | 6 | 3to5 | 3.5 | 3 | 2.15 | 3.6 | 0.05 |

| Twp-Range | 25-1w | species | # WTs | Diam Range | av Diam | Median Diam | rel Freq% | QMD | rel DOM% |
|---|---|---|---|---|---|---|---|---|---|
| # allWTs | 183 | red alder | 7 | 3to18 | 7.3 | 6 | 3.83 | 8.6 | 0.63 |
| all avDiam | 17.9 | redcedar | 49 | 7to50 | 16 | 15 | 26.78 | 17.9 | 19.00 |
| all QMD | 21.3 | cherry | 1 | 4 | 4 | 4 | 0.55 | 4 | 0.02 |
| SrvyYr | 1859 | Douglas-fir | 95 | 3to55 | 21.6 | 18 | 51.91 | 25.3 | 73.58 |
| | | hemlock | 20 | 4to18 | 10 | 9.5 | 10.93 | 10.6 | 2.72 |
| | | bl maple | 7 | 15to20 | 17.6 | 18 | 3.83 | 17.7 | 2.65 |
| | | madrone | 3 | 16to20 | 18 | 18 | 1.64 | 18.1 | 1.19 |
| | | willow | 1 | 10 | 10 | 10 | 0.55 | 10 | 0.12 |

| Twp-Range | 25-2w | species | # WTs | Diam Range | av Diam | Median Diam | rel Freq% | QMD | rel DOM% |
|---|---|---|---|---|---|---|---|---|---|
| # allWTs | 137 | red alder | 2 | 12to14 | 13 | 13 | 1.46 | 13 | 0.52 |
| all avDiam | 19.4 | redcedar | 25 | 7to45 | 17.6 | 15 | 18.25 | 19.5 | 14.54 |
| all QMD | 21.8 | Douglas-fir | 66 | 10to55 | 23.3 | 19 | 48.18 | 25.9 | 67.74 |
| SrvyYr | 1871 | hemlock | 35 | 7to30 | 14.6 | 14 | 25.55 | 15.9 | 13.54 |
| | | bl maple | 3 | 14 | 14 | 14 | 2.19 | 14 | 0.90 |
| | | madrone | 5 | 8to24 | 16.4 | 20 | 3.65 | 17.5 | 2.34 |
| | | spruce | 1 | 14 | 14 | 14 | 0.73 | 14 | 0.30 |

| Twp-Range | 24-1e | species | # WTs | Diam Range | av Diam | Median Diam | rel Freq% | QMD | rel DOM% |
|---|---|---|---|---|---|---|---|---|---|
| # allWTs | 222 | red alder | 27 | 3to15 | 7 | 6 | 12.16 | 7.8 | 1.70 |
| all avDiam | 16.7 | ash | 1 | 6 | 6 | 6 | 0.45 | 6 | 0.04 |
| all QMD | 20.9 | redcedar | 33 | 7to50 | 20.2 | 18 | 14.86 | 23.1 | 18.23 |
| SrvyYr | 1858 | cherry | 1 | 5 | 5 | 5 | 0.45 | 5 | 0.03 |
| | | crabapple | 2 | 3to6 | 4.5 | 4.5 | 0.90 | 4.7 | 0.05 |
| | | dogwood | 3 | 5to12 | 7.7 | 6 | 1.35 | 8.3 | 0.21 |
| | | elder | 1 | 4 | 4 | 4 | 0.45 | 4 | 0.02 |
| | | Douglas-fir | 107 | 3to72 | 22.2 | 18 | 48.20 | 25.6 | 72.59 |
| | | hemlock | 26 | 5to40 | 10.3 | 8 | 11.71 | 12.6 | 4.27 |
| | | bl maple | 10 | 3to30 | 10.3 | 4.5 | 4.50 | 14.4 | 2.15 |
| | | madrone | 3 | 9to10 | 9.7 | 10 | 1.35 | 9.7 | 0.29 |
| | | pine | 2 | 6to10 | 7 | 7 | 0.90 | 8.2 | 0.14 |
| | | skunkwood | 1 | 4 | 4 | 4 | 0.45 | 4 | 0.02 |
| | | willow | 5 | 2to4 | 2.4 | 3 | 2.25 | 3.1 | 0.05 |

| Twp-Range | 24-2e | species | # WTs | Diam Range | av Diam | Median Diam | rel Freq% | QMD | rel DOM% |
|---|---|---|---|---|---|---|---|---|---|
| # allWTs | 108 | red alder | 18 | 3to16 | 5.6 | 4 | 16.67 | 6.5 | 1.63 |
| all avDiam | 16.3 | redcedar | 14 | 8to40 | 23 | 10 | 12.96 | 25.6 | 19.61 |
| all QMD | 20.8 | cherry | 1 | 3 | 3 | 3 | 0.93 | 3 | 0.02 |
| SrvyYr | 1858 | Douglas-fir | 64 | 3to50 | 18.9 | 15.5 | 59.26 | 23.2 | 73.64 |
| | | hemlock | 7 | 4to36 | 13.3 | 9 | 6.48 | 17.4 | 4.53 |
| | | bl maple | 2 | 10to12 | 11 | 11 | 1.85 | 11 | 0.52 |
| | | spruce | 1 | 7 | 7 | 7 | 0.93 | 7 | 0.10 |
| | | willow | 1 | 3 | 3 | 3 | 0.93 | 3 | 0.02 |

| Twp-Range | 24-3e | species | # WTs | Diam Range | av Diam | Median Diam | rel Freq% | QMD | rel DOM% |
|---|---|---|---|---|---|---|---|---|---|
| # allWTs | 82 | red alder | 8 | 3to9 | 6 | 6 | 9.76 | 6.5 | 1.37 |
| all avDiam | 13.1 | redcedar | 31 | 3to24 | 14.3 | 12 | 37.80 | 15.5 | 30.27 |
| all QMD | 17.3 | dogwood | 1 | 3 | 3 | 3 | 1.22 | 3 | 0.04 |
| SrvyYr | 1862 | Douglas-fir | 29 | 3to50 | 17.5 | 12 | 35.37 | 23.7 | 66.21 |
| | | hazel | 2 | 3 | 3 | 3 | 2.44 | 3 | 0.07 |
| | | hemlock | 4 | 9to11 | 10.3 | 10.5 | 4.88 | 10.3 | 1.72 |
| | | bl maple | 2 | 3 | 3 | 3 | 2.44 | 3 | 0.07 |
| | | willow | 4 | 3 | 3 | 3 | 4.88 | 3 | 0.15 |
| | | yew | 1 | 8 | 8 | 8 | 1.22 | 8 | 0.26 |

| Twp-Range | 24-4e | species | # WTs | Diam Range | av Diam | Median Diam | rel Freq% | QMD | rel DOM% |
|---|---|---|---|---|---|---|---|---|---|
| # allWTs | 196 | red alder | 36 | 3to18 | 7.1 | 7 | 18.37 | 7.9 | 4.95 |
| all avDiam | 12.2 | ash | 7 | 3to12 | 8 | 9 | 3.57 | 8.5 | 1.11 |
| all QMD | 15.2 | barberry | 2 | 8 | 8 | 8 | 1.02 | 8 | 0.28 |
| SrvyYr | 1861 | redcedar | 32 | 4to40 | 16.4 | 12 | 16.33 | 19.4 | 26.53 |
| | | cherry | 1 | 8 | 8 | 8 | 0.51 | 8 | 0.14 |
| | | crabapple | 5 | 3to8 | 4.4 | 4 | 2.55 | 4.8 | 0.25 |
| | | cottonwood | 14 | 4to36 | 13.7 | 9.5 | 7.14 | 16.7 | 8.60 |
| | | dogwood | 3 | 3to11 | 5.7 | 3 | 1.53 | 6.8 | 0.31 |
| | | Douglas-fir | 70 | 3to40 | 15.6 | 11.5 | 35.71 | 18.5 | 52.78 |
| | | hemlock | 3 | 8to10 | 8.7 | 8 | 1.53 | 8.7 | 0.50 |
| | | bl maple | 9 | 5to20 | 10.7 | 10 | 4.59 | 11.5 | 2.62 |
| | | madrone | 2 | 3to40 | 3.5 | 3.5 | 1.02 | 3.5 | 0.05 |
| | | vine maple | 2 | 4to5 | 4.5 | 4.5 | 1.02 | 4.5 | 0.09 |
| | | willow | 9 | 3to15 | 7.2 | 5 | 4.59 | 8.2 | 1.33 |
| | | yew | 1 | 9 | 9 | 9 | 0.51 | 9 | 0.18 |

| Twp-Range | 24-5e | species | # WTs | Diam Range | av Diam | Median Diam | rel Freq% | QMD | rel DOM% |
|---|---|---|---|---|---|---|---|---|---|
| # allWTs | 252 | red alder | 30 | 3to15 | 5.7 | 5 | 11.90 | 6.3 | 2.64 |
| all avDiam | 10.9 | ash | 3 | 3to6 | 4 | 3 | 1.19 | 4.2 | 0.12 |
| all QMD | 13.4 | redcedar | 73 | 3to 36 | 12.4 | 10 | 28.97 | 14.4 | 33.59 |
| SrvyYr | 1864 | cherry | 1 | 3 | 3 | 3 | 0.40 | 3 | 0.02 |
| | | crabapple | 1 | 3 | 3 | 3 | 0.40 | 3 | 0.02 |
| | | Douglas-fir | 79 | 3to40 | 14.8 | 14 | 31.35 | 17.3 | 52.46 |
| | | hemlock | 25 | 3to30 | 9.2 | 8 | 9.92 | 10.6 | 6.23 |
| | | hazel | 1 | 3 | 3 | 3 | 0.40 | 3 | 0.02 |
| | | bl maple | 16 | 3to18 | 8.3 | 7.5 | 6.35 | 9.6 | 3.27 |
| | | pine | 4 | 5to12 | 8.8 | 9 | 1.59 | 9.3 | 0.77 |
| | | vine maple | 7 | 3to7 | 4.1 | 3 | 2.78 | 4.4 | 0.30 |
| | | willow | 12 | 3to7 | 3.6 | 3 | 4.76 | 3.8 | 0.38 |

| Twp-Range | 24-6e | species | # WTs | Diam Range | av Diam | Median Diam | rel Freq% | QMD | rel DOM% |
|---|---|---|---|---|---|---|---|---|---|
| # allWTs | 249 | red alder | 45 | 3to24 | 6.5 | 5 | 18.07 | 7.9 | 5.78 |
| all avDiam | 9.8 | ash | 1 | 4 | 4 | 4 | 0.40 | 4 | 0.03 |
| all QMD | 14.0 | barberry | 2 | 7to9 | 8 | 8 | 0.80 | 8.1 | 0.27 |
| SrvyYr | 1864 | redcedar | 37 | 6to40 | 17.5 | 12 | 14.86 | 20.5 | 31.99 |
| | | cherry | 8 | 3to4 | 3.3 | 3 | 3.21 | 3.3 | 0.18 |
| | | crabapple | 2 | 3 | 3 | 3 | 0.80 | 3 | 0.04 |
| | | dogwood | 2 | 3to4 | 3.5 | 3.5 | 0.80 | 3.5 | 0.05 |
| | | Douglas-fir | 94 | 3to72 | 11.3 | 8 | 37.75 | 16.2 | 50.75 |
| | | hemlock | 4 | 8to40 | 17 | 10 | 1.61 | 21.6 | 3.84 |
| | | bl maple | 26 | 3to36 | 7.4 | 4 | 10.44 | 10.6 | 6.01 |
| | | vine maple | 12 | 3to7 | 4.3 | 4 | 4.82 | 4.4 | 0.48 |
| | | willow | 16 | 3to11 | 3.6 | 3 | 6.43 | 4.1 | 0.55 |

| Twp-Range | 24-7e | species | # WTs | Diam Range | av Diam | Median Diam | rel Freq% | QMD | rel DOM% |
|---|---|---|---|---|---|---|---|---|---|
| # allWTs | 286 | red alder | 23 | 3to14 | 6 | 5 | 8.04 | 6.7 | 0.98 |
| all avDiam | 12.6 | barberry | 3 | 4to7 | 5 | 4 | 1.05 | 5.2 | 0.08 |
| all QMD | 19.2 | redcedar | 36 | 3to120 | 19.5 | 10 | 12.59 | 30.5 | 31.66 |
| SrvyYr | 1873 | cherry | 9 | 3to10 | 4.7 | 4 | 3.15 | 5.3 | 0.24 |
| | | crabapple | 1 | 2 | 2 | 2 | 0.35 | 2 | 0.00 |
| | | Douglas-fir | 72 | 3to100 | 15.6 | 8 | 25.17 | 23.8 | 38.56 |
| | | hemlock | 69 | 3to36 | 13.7 | 12 | 24.13 | 16.2 | 17.12 |
| | | bl maple | 25 | 3to45 | 13.1 | 12 | 8.74 | 16.9 | 6.75 |
| | | spruce | 3 | 12to42 | 28 | 30 | 1.05 | 30.6 | 2.66 |
| | | vine maple | 39 | 3to36 | 4.9 | 3 | 13.64 | 6.9 | 1.76 |
| | | willow | 4 | 3to45 | 3.3 | 3 | 1.40 | 3.3 | 0.04 |
| | | yew | 2 | 7to10 | 8.5 | 8.5 | 0.70 | 8.6 | 0.14 |

| Twp-Range | 24-8e | species | # WTs | Diam Range | av Diam | Median Diam | rel Freq% | QMD | rel DOM% |
|---|---|---|---|---|---|---|---|---|---|
| # allWTs | 213 | red alder | 23 | 3to24 | 6.3 | 5 | 10.80 | 7.4 | 3.86 |
| all avDiam | 9.1 | barberry | 4 | 6to10 | 8 | 8 | 1.88 | 8.1 | 0.80 |
| all QMD | 12.4 | redcedar | 21 | 3to48 | 18.8 | 12 | 9.86 | 23.4 | 35.26 |
| SrvyYr | 1865 | cherry | 2 | 3to5 | 4 | 4 | 0.94 | 4.4 | 0.12 |
| | | crabapple | 2 | 3to5 | 4 | 4 | 0.94 | 4.1 | 0.10 |
| | | cottonwood | 1 | 10 | 10 | 10 | 0.47 | 10 | 0.31 |
| | | Douglas-fir | 29 | 3to50 | 10.2 | 8 | 13.62 | 12.2 | 13.23 |
| | | hemlock | 82 | 3to36 | 10.5 | 9 | 38.50 | 12.5 | 39.29 |
| | | hazel | 5 | 3to5 | 3.4 | 3 | 2.35 | 3.5 | 0.19 |
| | | bl maple | 7 | 3to16 | 5.3 | 3 | 3.29 | 6.9 | 1.02 |
| | | spruce | 1 | 8 | 8 | 8 | 0.47 | 8 | 0.20 |
| | | vine maple | 35 | 3to5 | 34 | 3 | 16.43 | 3.5 | 1.31 |
| | | willow | 1 | 5 | 5 | 5 | 0.47 | 5 | 0.08 |

| Twp-Range | 24-1w | species | # WTs | Diam Range | av Diam | Median Diam | rel Freq% | QMD | rel DOM% |
|---|---|---|---|---|---|---|---|---|---|
| # allWTs | 287 | red alder | 11 | 4to10 | 6.7 | 6 | 3.83 | 6.9 | 0.40 |
| all avDiam | 17.9 | redcedar | 35 | 6to44 | 16 | 14 | 12.20 | 17.7 | 8.45 |
| all QMD | 21.3 | dogwood | 1 | 12 | 12 | 12 | 0.35 | 12 | 0.11 |
| SrvyYr | 1880 | Douglas-fir | 193 | 3to50 | 20.9 | 20 | 67.25 | 24 | 85.69 |
| | | hemlock | 41 | 3to30 | 9.6 | 9 | 14.29 | 11 | 3.82 |
| | | bl maple | 2 | 10 | 10 | 10 | 0.70 | 10 | 0.15 |
| | | pine | 3 | 14 | 4 | | 1.05 | 19.9 | 0.92 |
| | | cascara | 1 | 8 | 8 | 8 | 0.35 | 8 | 0.05 |

| Twp-Range | 24-2w | species | # WTs | Diam Range | av Diam | Median Diam | rel Freq% | QMD | rel DOM% |
|---|---|---|---|---|---|---|---|---|---|
| # allWTs | 251 | red alder | 20 | 3to12 | 6.5 | 5 | 7.97 | 7.2 | 1.49 |
| all avDiam | 13.8 | redcedar | 40 | 4to40 | 18 | 14 | 15.94 | 20.7 | 24.56 |
| all QMD | 16.7 | Douglas-fir | 133 | 3to50 | 16 | 12 | 52.99 | 18.6 | 65.94 |
| SrvyYr | 1877 | hemlock | 43 | 3to30 | 8.5 | 8 | 17.13 | 9.8 | 5.92 |
| | | bl maple | 2 | 8to20 | 14 | 14 | 0.80 | 14 | 0.56 |
| | | pine | 8 | 6to24 | 10.5 | 9.5 | 3.19 | 11.8 | 1.60 |
| | | cascara | 1 | 4 | 4 | 4 | 0.40 | 4 | 0.02 |
| | | vine maple | 1 | 4 | 4 | 4 | 0.40 | 4 | 0.02 |
| | | willow | 3 | 4to5 | 4.7 | 5 | 1.20 | 4.7 | 0.09 |

| Twp-Range | 24-3w | species | # WTs | Diam Range | av Diam | Median Diam | rel Freq% | QMD | rel DOM% |
|---|---|---|---|---|---|---|---|---|---|
| # allWTs | 136 | red alder | 1 | 14 | 14 | 14 | 0.74 | 14 | 0.41 |
| all avDiam | 16.1 | redcedar | 12 | 5to30 | 16.8 | 14 | 8.82 | 18.3 | 8.37 |
| all QMD | 18.8 | dogwood | 1 | 4 | 4 | 4 | 0.74 | 4 | 0.03 |
| SrvyYr | 1875 | Douglas-fir | 93 | 4to50 | 17.4 | 15 | 68.38 | 19.8 | 75.98 |
| | | grand fir | 2 | 30to50 | 40 | 40 | 1.47 | 41.2 | 7.07 |
| | | hemlock | 18 | 4to24 | 10.1 | 9 | 13.24 | 11 | 4.54 |
| | | bl maple | 4 | 7to36 | 16 | 10.5 | 2.94 | 19.8 | 3.27 |
| | | pine | 1 | 7 | 7 | 7 | 0.74 | 7 | 0.10 |
| | | vine maple | 3 | 3to4 | 3.7 | 4 | 2.21 | 3.7 | 0.09 |
| | | yew | 1 | 8 | 8 | 8 | 0.74 | 8 | 0.13 |

| Twp-Range | 23-1e | species | # WTs | Diam Range | av Diam | Median Diam | rel Freq% | QMD | rel DOM% |
|---|---|---|---|---|---|---|---|---|---|
| # allWTs | 278 | red alder | 29 | 3to14 | 5.3 | 5 | 10.43 | 5.7 | 0.91 |
| all avDiam | 15.6 | aspen | 2 | 4to5 | 4.5 | 4.5 | 0.72 | 4.5 | 0.04 |
| all QMD | 19.3 | redcedar | 19 | 6to45 | 16.1 | 16 | 6.83 | 18.1 | 6.02 |
| SrvyYr | 1858 | cherry | 3 | 3to4 | 3.7 | 4 | 1.08 | 3.7 | 0.04 |
| | | dogwood | 3 | 4to5 | 4.3 | 4 | 1.08 | 4.4 | 0.06 |
| | | Douglas-fir | 177 | 3to90 | 20 | 18 | 63.67 | 23 | 90.55 |
| | | grand fir | 2 | 12 | 12 | 12 | 0.72 | 12 | 0.28 |
| | | hemlock | 12 | 4to22 | 7.8 | 7 | 4.32 | 9.1 | 0.96 |
| | | hazel | 1 | 3 | 3 | 3 | 0.36 | 3 | 0.01 |
| | | bl maple | 1 | 4 | 4 | 4 | 0.36 | 4 | 0.02 |
| | | madrone | 1 | 4 | 4 | 4 | 0.36 | 4 | 0.02 |
| | | pine | 15 | 4to18 | 8.7 | 8 | 5.40 | 9.5 | 1.31 |
| | | spruce | 1 | 5 | 5 | 5 | 0.36 | 5 | 0.02 |
| | | willow | 12 | 3to4 | 3.8 | 4 | 4.32 | 3.9 | 0.18 |

| Twp-Range | 23-2e | species | # WTs | Diam Range | av Diam | Median Diam | rel Freq% | QMD | rel DOM% |
|---|---|---|---|---|---|---|---|---|---|
| # allWTs | 163 | red alder | 26 | 3to16 | 6.4 | 5.5 | 15.95 | 7 | 1.42 |
| all avDiam | 18.4 | redcedar | 18 | 5to30 | 15.2 | 12 | 11.04 | 16.9 | 5.73 |
| all QMD | 23.5 | cherry | 2 | 4 | 4 | 4 | 1.23 | 4 | 0.04 |
| SrvyYr | 1858 | crabapple | 1 | 6 | 6 | 6 | 0.61 | 6 | 0.04 |
| | | elder | 1 | 4 | 4 | 4 | 0.61 | 4 | 0.02 |
| | | Douglas-fir | 100 | 3to90 | 24.3 | 20 | 61.35 | 28.6 | 91.13 |
| | | hemlock | 8 | 5to10 | 7 | 7 | 4.91 | 7.2 | 0.46 |
| | | bl maple | 5 | 3to28 | 11.4 | 6 | 3.07 | 14.7 | 1.20 |
| | | willow | 2 | 3to4 | 3.5 | 3.5 | 1.23 | 3.5 | 0.03 |

| Twp-Range | 23-4e | species | # WTs | Diam Range | av Diam | Median Diam | rel Freq% | QMD | rel DOM% |
|---|---|---|---|---|---|---|---|---|---|
| # allWTs | 271 | red alder | 40 | 3to24 | 6.1 | 4 | 14.76 | 7.3 | 2.63 |
| all avDiam | 12.2 | ash | 13 | 3to18 | 8.3 | 8.5 | 4.80 | 9.1 | 1.33 |
| all QMD | 17.3 | barberry | 2 | 3to6 | 4.5 | 4.5 | 0.74 | 4.7 | 0.05 |
| SrvyYr | 1862 | redcedar | 59 | 3to100 | 17.5 | 12 | 21.77 | 23.1 | 38.86 |
| | | cherry | 3 | 3to4 | 3.3 | 3 | 1.11 | 3.4 | 0.04 |
| | | crabapple | 3 | 5to10 | 7 | 6 | 1.11 | 7.3 | 0.20 |
| | | cottonwood | 4 | 3to40 | 23.5 | 25.5 | 1.48 | 28.9 | 4.12 |
| | | dogwood | 1 | 3 | 3 | 3 | 0.37 | 3 | 0.01 |
| | | Douglas-fir | 98 | 3to70 | 14.9 | 11 | 36.16 | 20 | 48.38 |
| | | hazel | 1 | 3 | 3 | 3 | 0.37 | 3 | 0.01 |
| | | hemlock | 2 | 10to12 | 11 | 11 | 0.74 | 11 | 0.30 |
| | | bl maple | 8 | 3to25 | 12.9 | 14 | 2.95 | 15.3 | 2.31 |
| | | vine maple | 4 | 3to8 | 5.5 | 5.5 | 1.48 | 5.8 | 0.17 |
| | | willow | 33 | 2to18 | 5.3 | 4 | 12.18 | 6.3 | 1.62 |

| Twp-Range | 23-5e | species | # WTs | Diam Range | av Diam | Median Diam | rel Freq% | QMD | rel DOM% |
|---|---|---|---|---|---|---|---|---|---|
| # allWTs | 255 | red alder | 40 | 3to22 | 6.6 | 5 | 15.69 | 7.9 | 4.56 |
| all avDiam | 11.0 | ash | 3 | 8to10 | 9 | 9 | 1.18 | 9 | 0.44 |
| all QMD | 14.7 | barberry | 2 | 8to11 | 9.5 | 9.5 | 0.78 | 9.5 | 0.33 |
| SrvyYr | 1865 | redcedar | 44 | 4to72 | 14.1 | 11.5 | 17.25 | 17.9 | 25.73 |
| | | cherry | 4 | 3to9 | 5 | 4 | 1.57 | 5.6 | 0.23 |
| | | crabapple | 7 | 3to11 | 5.9 | 5 | 2.75 | 6.4 | 0.52 |
| | | cottonwood | 3 | 10to36 | 19.3 | 12 | 1.18 | 22.7 | 2.82 |
| | | dogwood | 1 | 10 | 10 | 10 | 0.39 | 10 | 0.18 |
| | | Douglas-fir | 71 | 3to72 | 15 | 12 | 27.84 | 19.7 | 50.28 |
| | | hemlock | 40 | 6to30 | 11.4 | 10 | 15.69 | 12.6 | 11.59 |
| | | hazel | 1 | 3 | 3 | 3 | 0.39 | 3 | 0.02 |
| | | bl maple | 10 | 5to14 | 10 | 11 | 3.92 | 10.4 | 1.97 |
| | | spruce | 1 | 14 | 14 | 14 | 0.39 | 14 | 0.36 |
| | | vine maple | 18 | 2to6 | 3.5 | 3 | 7.06 | 3.6 | 0.43 |
| | | willow | 10 | 3to6 | 3.7 | 3 | 3.92 | 3.9 | 0.28 |

| Twp-Range | 23-6e | species | # WTs | Diam Range | av Diam | Median Diam | rel Freq% | QMD | rel DOM% |
|---|---|---|---|---|---|---|---|---|---|
| # allWTs | 270 | red alder | 24 | 3to24 | 9.5 | 8 | 8.89 | 11.3 | 3.27 |
| all avDiam | 13.5 | ash | 1 | 20 | 20 | 20 | 0.37 | 20 | 0.43 |
| all QMD | 18.6 | barberry | 4 | 3to6 | 5 | 6 | 1.48 | 5.4 | 0.12 |
| SrvyYr | 1873 | redcedar | 39 | 3to60 | 21.8 | 18 | 14.44 | 26.5 | 29.26 |
| | | cherry | 1 | 3 | 3 | 3 | 0.37 | 3 | 0.01 |
| | | cottonwood | 2 | 8 | 8 | 8 | 0.74 | 8 | 0.14 |
| | | Douglas-fir | 92 | 3to60 | 14.8 | 10 | 34.07 | 21.2 | 44.17 |
| | | grand fir | 1 | 5 | 5 | 5 | 0.37 | 5 | 0.03 |
| | | hemlock | 65 | 3to60 | 11.9 | 10 | 24.07 | 15.7 | 17.12 |
| | | bl maple | 21 | 3to24 | 12.3 | 12 | 7.78 | 14 | 4.40 |
| | | spruce | 2 | 6to20 | 13 | 13 | 0.74 | 14.8 | 0.47 |
| | | vine maple | 13 | 3to6 | 4 | 4 | 4.81 | 4.1 | 0.23 |
| | | willow | 5 | 3to4 | 3.6 | 4 | 1.85 | 3.7 | 0.07 |

| Twp-Range | 23-7e | species | # WTs | Diam Range | av Diam | Median Diam | rel Freq% | QMD | rel DOM% |
|---|---|---|---|---|---|---|---|---|---|
| # allWTs | 291 | red alder | 6 | 10to18 | 14 | 13.5 | 2.06 | 14.4 | 0.54 |
| all avDiam | 23.1 | redcedar | 31 | 3to120 | 43.7 | 40 | 10.65 | 50.1 | 33.88 |
| all QMD | 28.1 | Douglas-fir | 24 | 6to72 | 36.8 | 36 | 8.25 | 43.5 | 19.77 |
| SrvyYr | 1892 | grand fir | 15 | 8to24 | 15.3 | 12 | 5.15 | 16.8 | 1.84 |
| | | hemlock | 193 | 3to60 | 19.4 | 20 | 66.32 | 21.4 | 38.48 |
| | | bl maple | 3 | 8to18 | 12 | 10 | 1.03 | 12.8 | 0.21 |
| | | spruce | 14 | 8to60 | 25.9 | 23 | 4.81 | 29.5 | 5.30 |
| | | vine maple | 4 | 3to6 | 3.8 | 3 | 1.37 | 4 | 0.03 |
| | | yew | 1 | 10 | 10 | 10 | 0.34 | 10 | 0.04 |

| Twp-Range | 23-8e | species | # WTs | Diam Range | av Diam | Median Diam | rel Freq% | QMD | rel DOM% |
|---|---|---|---|---|---|---|---|---|---|
| # allWTs | 274 | red alder | 3 | 3to8 | 5.7 | 6 | 1.09 | 6 | 0.07 |
| all avDiam | 19.4 | barberry | 1 | 8 | 8 | 8 | 0.36 | 8 | 0.04 |
| all QMD | 24.0 | redcedar | 26 | 3to72 | 28.1 | 29 | 9.49 | 33 | 17.89 |
| SrvyYr | 1866&91 | Douglas-fir | 62 | 3to93 | 27.9 | 26 | 22.63 | 31.8 | 39.61 |
| | | grand fir | 17 | 9to110 | 18.3 | 18 | 6.20 | 19.7 | 4.17 |
| | | hemlock | 122 | 3to40 | 17.4 | 18 | 44.53 | 19.6 | 29.61 |
| | | bl maple | 4 | 3to28 | 18.3 | 21 | 1.46 | 20.6 | 1.07 |
| | | spruce | 8 | 8to100 | 26.3 | 18 | 2.92 | 38.8 | 7.61 |
| | | vine maple | 26 | 3to93 | 3.6 | 3 | 9.49 | 3.8 | 0.24 |
| | | willow | 5 | 3to5 | 3.4 | 3 | 1.82 | 3.5 | 0.04 |

| Twp-Range | 23-1w | species | # WTs | Diam Range | av Diam | Median Diam | rel Freq% | QMD | rel DOM% |
|---|---|---|---|---|---|---|---|---|---|
| # allWTs | 289 | red alder | 12 | 3to30 | 11.3 | 8 | 4.15 | 14.3 | 1.49 |
| all avDiam | 19.1 | redcedar | 15 | 6to120 | 36.9 | 30 | 5.19 | 47.7 | 20.73 |
| all QMD | 23.9 | cherry | 3 | 3to9 | 5.7 | 5 | 1.04 | 9.6 | 0.17 |
| SrvyYr | 1872 | crabapple | 4 | 4to6 | 5.3 | 5 | 1.38 | 5.3 | 0.07 |
| | | dogwood | 1 | 3 | 3 | 3 | 0.35 | 3 | 0.01 |
| | | Douglas-fir | 208 | 3to66 | 20.9 | 18 | 71.97 | 24 | 72.77 |
| | | grand fir | 2 | 18to30 | 24 | 24 | 0.69 | 24.7 | 0.74 |
| | | hemlock | 16 | 3to36 | 9.5 | 8 | 5.54 | 12.1 | 1.42 |
| | | bl maple | 7 | 3to36 | 16.7 | 12 | 2.42 | 21.4 | 1.95 |
| | | madrone | 1 | 4 | 4 | 4 | 0.35 | 4 | 0.01 |
| | | pine | 12 | 5to18 | 9.3 | 9 | 4.15 | 9.9 | 0.71 |
| | | skookumwo | 1 | 6 | 6 | 6 | 0.35 | 6 | 0.02 |
| | | willow | 7 | 3to4 | 3.1 | 3 | 2.42 | 3.2 | 0.04 |

| Twp-Range | 23-2w | species | # WTs | Diam Range | av Diam | Median Diam | rel Freq% | QMD | rel DOM% |
|---|---|---|---|---|---|---|---|---|---|
| # allWTs | 285 | red alder | 19 | 5to18 | 11.6 | 11 | 6.67 | 12.3 | 3.14 |
| all avDiam | 15.7 | redcedar | 8 | 8to40 | 18 | 14.5 | 2.81 | 20.5 | 3.67 |
| all QMD | 17.9 | crabapple | 1 | 8 | 8 | 8 | 0.35 | 8 | 0.07 |
| SrvyYr | 1874 | Douglas-fir | 194 | 3to50 | 17.7 | 15 | 68.07 | 19.8 | 83.08 |
| | | hemlock | 22 | 3to14 | 10.3 | 10 | 7.72 | 10.8 | 2.80 |
| | | madrone | 1 | 3 | 3 | 3 | 0.35 | 3 | 0.01 |
| | | pine | 40 | 4to48 | 10.6 | 10 | 14.04 | 12.7 | 7.05 |

| Twp-Range | 23-3w | species | # WTs | Diam Range | av Diam | Median Diam | rel Freq% | QMD | rel DOM% |
|---|---|---|---|---|---|---|---|---|---|
| # allWTs | 227 | red alder | 15 | 3to20 | 9.5 | 9 | 6.61 | 10.3 | 2.05 |
| all avDiam | 15.6 | redcedar | 19 | 3to60 | 16.1 | 12 | 8.37 | 20.3 | 10.10 |
| all QMD | 18.5 | Douglas-fir | 145 | 3to50 | 18.2 | 15 | 63.88 | 20.8 | 80.93 |
| SrvyYr | 1874 | hemlock | 21 | 5to14 | 9.5 | 9 | 9.25 | 9.8 | 2.60 |
| | | bl maple | 5 | 4to20 | 9.8 | 5 | 2.20 | 11.8 | 0.90 |
| | | madrone | 3 | 4to12 | 7 | 5 | 1.32 | 7.9 | 0.24 |
| | | pine | 17 | 3to30 | 9.8 | 9 | 7.49 | 11.4 | 2.85 |
| | | vine maple | 2 | 6 | 6 | 6 | 0.88 | 6 | 0.09 |

| Twp-Range | 23-4w | species | # WTs | Diam Range | av Diam | Median Diam | rel Freq% | QMD | rel DOM% |
|---|---|---|---|---|---|---|---|---|---|
| # allWTs | 251 | red alder | 4 | 8to10 | 8.8 | 8.5 | 1.59 | 8.8 | 0.46 |
| all avDiam | 13.3 | redcedar | 18 | 5to40 | 15.6 | 10 | 7.14 | 18.6 | 9.18 |
| all QMD | 16.4 | dogwood | 1 | 4 | 4 | 4 | 0.40 | 4 | 0.02 |
| SrvyYr | 1873 | Douglas-fir | 103 | 4to42 | 19 | 18 | 40.87 | 21.4 | 69.50 |
| | | hemlock | 66 | 3to40 | 10.8 | 10 | 26.19 | 12.7 | 15.69 |
| | | bl maple | 5 | 3to18 | 9.2 | 8 | 1.98 | 10.4 | 0.80 |
| | | pine | 3 | 8 | 8 | 8 | 1.19 | 8 | 0.28 |
| | | shittemwoo | 5 | 2to4 | 3 | 3 | 1.98 | 3 | 0.07 |
| | | vine maple | 40 | 3to28 | 5 | 4 | 15.87 | 6.4 | 2.41 |
| | | willow | 6 | 2to28 | 8.3 | 4 | 2.38 | 12.4 | 1.36 |

| Twp-Range | 22-1e | species | # WTs | Diam Range | av Diam | Median Diam | rel Freq% | QMD | rel DOM% |
|---|---|---|---|---|---|---|---|---|---|
| # allWTs | 228 | red alder | 26 | 4to18 | 6.2 | 5 | 11.40 | 7.1 | 1.46 |
| all avDiam | 16.4 | redcedar | 21 | 5to79 | 20.1 | 18 | 9.21 | 25 | 14.64 |
| all QMD | 19.8 | cherry | 1 | 3 | 3 | 3 | 0.44 | 3 | 0.01 |
| SrvyYr | 1857 | Douglas-fir | 143 | 3to84 | 19.8 | 18 | 62.72 | 22.4 | 80.01 |
| | | hemlock | 17 | 4to22 | 11 | 10 | 7.46 | 11.9 | 2.68 |
| | | bl maple | 3 | 3to16 | 7.3 | 3 | 1.32 | 9.6 | 0.31 |
| | | madrone | 1 | 4 | 4 | 4 | 0.44 | 4 | 0.02 |
| | | pine | 11 | 4to12 | 8.4 | 8 | 4.82 | 8.7 | 0.93 |
| | | willow | 5 | 3to4 | 3.2 | 3.2 | 2.19 | 3.2 | 0.06 |

| Twp-Range | 22-2e | species | # WTs | Diam Range | av Diam | Median Diam | rel Freq% | QMD | rel DOM% |
|---|---|---|---|---|---|---|---|---|---|
| # allWTs | 130 | red alder | 11 | 3to10 | 6.1 | 5 | 8.46 | 6.6 | 0.82 |
| all avDiam | 17.9 | redcedar | 17 | 8to48 | 21.1 | 16 | 13.08 | 24.6 | 17.54 |
| all QMD | 21.2 | cherry | 1 | 4 | 4 | 4 | 0.77 | 4 | 0.03 |
| SrvyYr | 1857 | Douglas-fir | 87 | 3to50 | 20.3 | 18 | 66.92 | 23 | 78.47 |
| | | hemlock | 8 | 4to12 | 7.8 | 8.5 | 6.15 | 8.2 | 0.92 |
| | | bl maple | 4 | 6to28 | 15.5 | 14 | 3.08 | 17.4 | 2.06 |
| | | madrone | 1 | 4 | 4 | 4 | 0.77 | 4 | 0.03 |
| | | skunkwood | 1 | 6 | 6 | 6 | 0.77 | 6 | 0.06 |

| Twp-Range | 22-4e | species | # WTs | Diam Range | av Diam | Median Diam | rel Freq% | QMD | rel DOM% |
|---|---|---|---|---|---|---|---|---|---|
| # allWTs | 231 | red alder | 34 | 3to34 | 6.7 | 5 | 14.72 | 9 | 4.68 |
| all avDiam | 11.7 | ash | 23 | 3to18 | 8.8 | 8 | 9.96 | 9.9 | 3.83 |
| all QMD | 16.0 | barberry | 6 | 3to18 | 6.3 | 3 | 2.60 | 8.4 | 0.72 |
| SrvyYr | 1863 | redcedar | 42 | 4to50 | 17 | 12.5 | 18.18 | 19.8 | 27.96 |
| | | crabapple | 5 | 4to8 | 5.8 | 6 | 2.16 | 6 | 0.31 |
| | | cottonwood | 6 | 12to50 | 33.5 | 36 | 2.60 | 35.5 | 12.84 |
| | | dogwood | 4 | 3to4 | 3.3 | 3 | 1.73 | 3.3 | 0.07 |
| | | Douglas-fir | 44 | 3to79 | 18.5 | 13.5 | 19.05 | 23 | 39.53 |
| | | hemlock | 2 | 12to13 | 12.5 | 12.5 | 0.87 | 12.5 | 0.53 |
| | | bl maple | 17 | 3to34 | 9.5 | 4 | 7.36 | 13.2 | 5.03 |
| | | spruce | 1 | 36 | 36 | 36 | 0.43 | 36 | 2.20 |
| | | vine maple | 25 | 3to12 | 5.6 | 5 | 10.82 | 6.2 | 1.63 |
| | | willow | 22 | 3to9 | 4.5 | 3.5 | 9.52 | 4.9 | 0.90 |

| Twp-Range | 22-5e | species | # WTs | Diam Range | av Diam | Median Diam | rel Freq% | QMD | rel DOM% |
|---|---|---|---|---|---|---|---|---|---|
| # allWTs | 261 | red alder | 41 | 3to24 | 10.7 | 10 | 15.71 | 12.2 | 5.32 |
| all avDiam | 16.6 | ash | 5 | 6to30 | 16.8 | 12 | 1.92 | 19 | 1.57 |
| all QMD | 21.0 | barberry | 2 | 8to15 | 11.5 | 11.5 | 0.77 | 12 | 0.25 |
| SrvyYr | 1867 | redcedar | 6 | 8to60 | 24.7 | 16 | 2.30 | 30.6 | 4.90 |
| | | cherry | 4 | 7to30 | 16.3 | 14 | 1.53 | 18.8 | 1.23 |
| | | crabapple | 1 | 4 | 4 | 4 | 0.38 | 4 | 0.01 |
| | | cottonwood | 2 | 36 | 36 | 36 | 0.77 | 36 | 2.26 |
| | | dogwood | 3 | 4to7 | 5 | 4 | 1.15 | 5.2 | 0.07 |
| | | Douglas-fir | 72 | 6to80 | 26.8 | 24 | 27.59 | 30.7 | 59.20 |
| | | hemlock | 86 | 3to50 | 14.5 | 12 | 32.95 | 17 | 21.68 |
| | | bl maple | 19 | 4to30 | 11.1 | 8 | 7.28 | 13.5 | 3.02 |
| | | vine maple | 17 | 3to12 | 4.5 | 4 | 6.51 | 5 | 0.37 |
| | | willow | 3 | 4to14 | 8 | 6 | 1.15 | 9.1 | 0.22 |

| Twp-Range | 22-6e | species | # WTs | Diam Range | av Diam | Median Diam | rel Freq% | QMD | rel DOM% |
|---|---|---|---|---|---|---|---|---|---|
| # allWTs | 289 | red alder | 26 | 3to16 | 8.4 | 8 | 9.00 | 9.1 | 2.18 |
| all avDiam | 13.9 | barberry | 6 | 4to6 | 5.2 | 5 | 2.08 | 5.2 | 0.16 |
| all QMD | 18.5 | redcedar | 5 | 4to50 | 18.8 | 12 | 1.73 | 25.5 | 3.29 |
| SrvyYr | 1880 | cherry | 7 | 5to8 | 6.4 | 6 | 2.42 | 6.5 | 0.30 |
| | | cottonwood | 2 | 14 | 14 | 14 | 0.69 | 14 | 0.40 |
| | | dogwood | 1 | 3 | 3 | 3 | 0.35 | 3 | 0.01 |
| | | Douglas-fir | 143 | 3to60 | 18.8 | 14 | 49.48 | 23.5 | 79.93 |
| | | hemlock | 53 | 3to30 | 10.5 | 8 | 18.34 | 12.7 | 8.65 |
| | | hazel | 1 | 4 | 4 | 4 | 0.35 | 4 | 0.02 |
| | | bl maple | 19 | 3to30 | 12.9 | 12 | 6.57 | 15.3 | 4.50 |
| | | spruce | 1 | 20 | 20 | 20 | 0.35 | 20 | 0.40 |
| | | vine maple | 21 | 3to7 | 4.2 | 4 | 7.27 | 4.4 | 0.41 |
| | | willow | 4 | 3to5 | 3.5 | 3 | 1.38 | 3.6 | 0.05 |

| Twp-Range | 22-7e | species | # WTs | Diam Range | av Diam | Median Diam | rel Freq% | QMD | rel DOM% |
|---|---|---|---|---|---|---|---|---|---|
| # allWTs | 290 | red alder | 7 | 6to20 | 10.4 | 9 | 2.41 | 11.3 | 0.42 |
| all avDiam | 23.0 | redcedar | 41 | 10to72 | 26.2 | 24 | 14.14 | 29.2 | 16.58 |
| all QMD | 27.0 | Douglas-fir | 125 | 3to84 | 26.4 | 28 | 43.10 | 31.3 | 58.09 |
| SrvyYr | 1891 | hemlock | 97 | 3to40 | 19.2 | 20 | 33.45 | 20.9 | 20.10 |
| | | bl maple | 8 | 5to36 | 19.9 | 19 | 2.76 | 22.8 | 1.97 |
| | | spruce | 8 | 10to50 | 23.3 | 18 | 2.76 | 26.9 | 2.75 |
| | | vine maple | 3 | 3to40 | 3.7 | 3 | 1.03 | 3.4 | 0.02 |
| | | willow | 1 | 3 | 3 | 3 | 0.34 | 3 | 0.00 |

| Twp-Range | 22-1w | species | # WTs | Diam Range | av Diam | Median Diam | rel Freq% | QMD | rel DOM% |
|---|---|---|---|---|---|---|---|---|---|
| # allWTs | 221 | red alder | 11 | 3to14 | 5.7 | 4 | 4.98 | 6.6 | 0.44 |
| all avDiam | 19.2 | redcedar | 10 | 10to40 | 22.3 | 24 | 4.52 | 24.5 | 5.50 |
| all QMD | 22.2 | dogwood | 1 | 5 | 5 | 5 | 0.45 | 5 | 0.02 |
| SrvyYr | 1856 | Douglas-fir | 184 | 3to84 | 20.5 | 18 | 83.26 | 23.3 | 91.60 |
| | | hemlock | 2 | 8to16 | 12 | 12 | 0.90 | 12.6 | 0.29 |
| | | bl maple | 4 | 12to30 | 18.5 | 16 | 1.81 | 19.8 | 1.44 |
| | | pine | 9 | 4to12 | 8.2 | 8 | 4.07 | 8.7 | 0.62 |

| Twp-Range | 22-2w | species | # WTs | Diam Range | av Diam | Median Diam | rel Freq% | QMD | rel DOM% |
|---|---|---|---|---|---|---|---|---|---|
| # allWTs | 231 | red alder | 15 | 6to18 | 11.3 | 12 | 6.49 | 12.1 | 2.02 |
| all avDiam | 18.3 | barberry | 1 | 8 | 8 | 8 | 0.43 | 8 | 0.06 |
| all QMD | 21.7 | redcedar | 12 | 12to50 | 21 | 13 | 5.19 | 24.4 | 6.56 |
| SrvyYr | 1861 | dogwood | 2 | 8 | 8 | 8 | 0.87 | 8 | 0.12 |
| | | Douglas-fir | 168 | 4to80 | 20.9 | 19 | 72.73 | 24 | 88.80 |
| | | hemlock | 11 | 8to20 | 12.7 | 12 | 4.76 | 13.3 | 1.79 |
| | | bl maple | 2 | 4to5 | 4.5 | 4.5 | 0.87 | 4.5 | 0.04 |
| | | madrone | 2 | 4to5 | 4.5 | 4.5 | 0.87 | 4.5 | 0.04 |
| | | pine | 15 | 4to16 | 6.3 | 5 | 6.49 | 7 | 0.67 |
| | | vine maple | 2 | 6to10 | 8 | 8 | 0.87 | 8.2 | 0.12 |
| | | yew | 1 | 8 | 8 | 8 | 0.43 | 8 | 0.06 |

| Twp-Range | 22-3w | species | # WTs | Diam Range | av Diam | Median Diam | rel Freq% | QMD | rel DOM% |
|---|---|---|---|---|---|---|---|---|---|
| # allWTs | 215 | red alder | 8 | 5to20 | 10.6 | 10 | 3.72 | 11.8 | 1.38 |
| all avDiam | 16.1 | redcedar | 13 | 6to42 | 20.3 | 20 | 6.05 | 22.6 | 8.25 |
| all QMD | 19.3 | dogwood | 3 | 4to6 | 5.3 | 6 | 1.40 | 5.4 | 0.11 |
| SrvyYr | 1861 | Douglas-fir | 131 | 4to60 | 19.3 | 18 | 60.93 | 22.4 | 81.71 |
| | | hemlock | 18 | 6to25 | 12.2 | 11 | 8.37 | 13.1 | 3.84 |
| | | bl maple | 6 | 5to24 | 15.2 | 16 | 2.79 | 16.3 | 1.98 |
| | | madrone | 8 | 4to12 | 5.5 | 4 | 3.72 | 6.2 | 0.38 |
| | | pine | 16 | 4to18 | 8.4 | 6 | 7.44 | 9.4 | 1.76 |
| | | vine maple | 8 | 4to8 | 5.1 | 4.5 | 3.72 | 5.3 | 0.28 |
| | | willow | 1 | 4 | 4 | 4 | 0.47 | 4 | 0.02 |
| | | yew | 3 | 6to15 | 9.7 | 8 | 1.40 | 10.4 | 0.40 |

| Twp-Range | 22-4w | species | # WTs | Diam Range | av Diam | Median Diam | rel Freq% | QMD | rel DOM% |
|---|---|---|---|---|---|---|---|---|---|
| # allWTs | 241 | red alder | 4 | 4to14 | 7.8 | 6.5 | 1.66 | 8.6 | 0.33 |
| all avDiam | 15.8 | redcedar | 19 | 4to28 | 15.4 | 14 | 7.88 | 16.9 | 6.08 |
| all QMD | 19.2 | Douglas-fir | 150 | 3to60 | 19.7 | 16 | 62.24 | 22.8 | 87.43 |
| SrvyYr | 1873 | hemlock | 19 | 4to20 | 10.3 | 9 | 7.88 | 11.3 | 2.72 |
| | | hazel | 1 | 3 | 3 | 3 | 0.41 | 3 | 0.01 |
| | | bl maple | 7 | 4to14 | 8.9 | 8 | 2.90 | 9.5 | 0.71 |
| | | pine | 22 | 3to24 | 8.6 | 8 | 9.13 | 9.7 | 2.32 |
| | | vine maple | 15 | 3to12 | 4.7 | 4 | 6.22 | 5.2 | 0.45 |
| | | willow | 3 | 2to4 | 3 | 3 | 1.24 | 3.1 | 0.03 |
| | | yew | 1 | 10 | 10 | 10 | 0.41 | 10 | 0.11 |

| Twp-Range | 22-5w | species | # WTs | Diam Range | av Diam | Median Diam | rel Freq% | QMD | rel DOM% |
|---|---|---|---|---|---|---|---|---|---|
| # allWTs | 197 | red alder | 1 | 6 | 6 | 6 | 0.51 | 6 | 0.03 |
| all avDiam | 20.3 | redcedar | 21 | 7to48 | 19.5 | 18 | 10.66 | 21.7 | 9.43 |
| all QMD | 23.1 | cottonwood | 4 | 10to40 | 22.5 | 20 | 2.03 | 25.6 | 2.50 |
| SrvyYr | 1895 | dogwood | 1 | 7 | 7 | 7 | 0.51 | 7 | 0.05 |
| | | Douglas-fir | 119 | 5to60 | 24 | 24 | 60.41 | 25.9 | 76.12 |
| | | grand fir | 1 | 18 | 18 | 18 | 0.51 | 18 | 0.31 |
| | | hemlock | 27 | 4to36 | 16.9 | 16 | 13.71 | 19.6 | 9.89 |
| | | bl maple | 1 | 4 | 4 | 4 | 0.51 | 4 | 0.02 |
| | | pine | 8 | 6to24 | 11.8 | 10 | 4.06 | 12.9 | 1.27 |
| | | vine maple | 14 | 2to7 | 4.2 | 4 | 7.11 | 4.4 | 0.26 |

| Twp-Range | 21-1e | species | # WTs | Diam Range | av Diam | Median Diam | rel Freq% | QMD | rel DOM% |
|---|---|---|---|---|---|---|---|---|---|
| # allWTs | 146 | red alder | 17 | 3to14 | 6.1 | 6 | 11.64 | 6.6 | 1.66 |
| all avDiam | 13.6 | barberry | 2 | 4to12 | 8 | 8 | 1.37 | 8.9 | 0.36 |
| all QMD | 17.5 | redcedar | 22 | 5to60 | 19 | 16 | 15.07 | 22 | 23.90 |
| SrvyYr | 1857 | cherry | 1 | 5 | 5 | 5 | 0.68 | 5 | 0.06 |
| | | cottonwood | 2 | 3to4 | 3.5 | 3.5 | 1.37 | 3.5 | 0.05 |
| | | Douglas-fir | 76 | 3to60 | 17.2 | 14 | 52.05 | 20.5 | 71.70 |
| | | hemlock | 5 | 3to10 | 6.4 | 6 | 3.42 | 6.8 | 0.52 |
| | | bl maple | 4 | 3to8 | 4.5 | 3.5 | 2.74 | 4.9 | 0.22 |
| | | madrone | 10 | 3to6 | 4 | 4 | 6.85 | 4.1 | 0.38 |
| | | willow | 6 | 3to8 | 4.8 | 4.5 | 4.11 | 5.1 | 0.35 |
| | | yew | 1 | 14 | 14 | 14 | 0.68 | 14 | 0.44 |

| Twp-Range | 21-2e | species | # WTs | Diam Range | av Diam | Median Diam | rel Freq% | QMD | rel DOM% |
|---|---|---|---|---|---|---|---|---|---|
| # allWTs | 167 | red alder | 17 | 3to15 | 7.4 | 6 | 10.18 | 8.4 | 2.44 |
| all avDiam | 13.8 | ash? | 2 | 6 | 6 | 6 | 1.20 | 6 | 0.15 |
| all QMD | 17.2 | redcedar | 33 | 4to40 | 18.2 | 16 | 19.76 | 20.9 | 29.31 |
| SrvyYr | 1857 | cherry | 1 | 7 | 7 | 7 | 0.60 | 7 | 0.10 |
| | | dogwood | 1 | 7 | 7 | 7 | 0.60 | 7 | 0.10 |
| | | Douglas-fir | 93 | 3to60 | 15 | 12 | 55.69 | 18.4 | 64.02 |
| | | hemlock | 11 | 3to16 | 8.4 | 12 | 6.59 | 9.4 | 1.98 |
| | | madrone | 5 | 3to28 | 9.8 | 4 | 2.99 | 13.6 | 1.88 |
| | | willow | 4 | 2to6 | 3.5 | 3 | 2.40 | 3.8 | 0.12 |

| Twp-Range | 21-3e | species | # WTs | Diam Range | av Diam | Median Diam | rel Freq% | QMD | rel DOM% |
|---|---|---|---|---|---|---|---|---|---|
| # allWTs | 73 | red alder | 8 | 3to12 | 7.3 | 7.5 | 10.96 | 7.9 | 1.27 |
| all avDiam | 19.1 | aspen | 2 | 4to5 | 4.5 | 4.5 | 2.74 | 4.5 | 0.10 |
| all QMD | 23.2 | redcedar | 10 | 4to24 | 18.2 | 20 | 13.70 | 19.6 | 9.77 |
| SrvyYr | 1873 | Douglas-fir | 43 | 3to57 | 24.2 | 24 | 58.90 | 27.9 | 85.10 |
| | | hemlock | 2 | 8to10 | 9 | 9 | 2.74 | 9.1 | 0.42 |
| | | bl maple | 1 | 4 | 4 | 4 | 1.37 | 4 | 0.04 |
| | | madrone | 1 | 12 | 12 | 12 | 1.37 | 12 | 0.37 |
| | | spruce | 5 | 8to20 | 13.6 | 15 | 6.85 | 14.6 | 2.71 |
| | | willow | 1 | 4 | 4 | 4 | 1.37 | 4 | 0.04 |

| Twp-Range | 21-4e | species | # WTs | Diam Range | av Diam | Median Diam | rel Freq% | QMD | rel DOM% |
|---|---|---|---|---|---|---|---|---|---|
| # allWTs | 271 | red alder | 49 | 2to32 | 6.3 | 5 | 18.08 | 8.2 | 2.41 |
| all avDiam | 17.9 | ash | 7 | 3to18 | 9.1 | 8 | 2.58 | 10.5 | 0.56 |
| all QMD | 22.5 | aspen | 2 | 6to12 | 9 | 9 | 0.74 | 9.5 | 0.13 |
| SrvyYr | 1868 | redcedar | 48 | 4to48 | 19.9 | 15 | 17.71 | 22.6 | 17.92 |
| | | cottonwood | 3 | 30to36 | 32 | 30 | 1.11 | 32.1 | 2.26 |
| | | Douglas-fir | 131 | 3to60 | 23.2 | 20 | 48.34 | 27.4 | 71.91 |
| | | hemlock | 15 | 3to30 | 12.2 | 10 | 5.54 | 13.8 | 2.09 |
| | | bl maple | 2 | 24to40 | 32 | 32 | 0.74 | 33 | 1.59 |
| | | spruce | 8 | 6to20 | 12.5 | 12.5 | 2.95 | 13.6 | 1.08 |
| | | vine maple | 1 | 8 | 8 | 8 | 0.37 | 8 | 0.05 |
| | | willow | 5 | 3to12 | 5.4 | 3 | 1.85 | 6.4 | 0.15 |

| Twp-Range | 21-5e | species | # WTs | Diam Range | av Diam | Median Diam | rel Freq% | QMD | rel DOM% |
|---|---|---|---|---|---|---|---|---|---|
| # allWTs | 244 | red alder | 49 | 3to22 | 9.3 | 8 | 20.08 | 10.6 | 5.09 |
| all avDiam | 16.5 | ash | 1 | 8 | 8 | 8 | 0.41 | 8 | 0.06 |
| all QMD | 21.1 | redcedar | 8 | 8to80 | 32.5 | 30 | 3.28 | 38.5 | 10.96 |
| SrvyYr | 1867 | cherry | 5 | 3to8 | 5.2 | 5 | 2.05 | 5.4 | 0.13 |
| | | cottonwood | 16 | 10to36 | 24.1 | 24 | 6.56 | 25.6 | 9.69 |
| | | dogwood | 3 | 3 | 3 | 3 | 1.23 | 3 | 0.02 |
| | | Douglas-fir | 116 | 3to60 | 20 | 18 | 47.54 | 24.2 | 62.78 |
| | | hemlock | 8 | 6to36 | 18 | 15 | 3.28 | 20.7 | 3.17 |
| | | bl maple | 14 | 4to50 | 16.8 | 16.5 | 5.74 | 20.9 | 5.65 |
| | | spruce | 3 | 18to30 | 26 | 30 | 1.23 | 26.6 | 1.96 |
| | | vine maple | 11 | 3to7 | 4.5 | 4 | 4.51 | 4.7 | 0.22 |
| | | willow | 9 | 3to6 | 4.2 | 4 | 3.69 | 4.3 | 0.15 |
| | | yew | 1 | 8 | 8 | 8 | 0.41 | 8 | 0.06 |

| Twp-Range | 21-6e | species | # WTs | Diam Range | av Diam | Median Diam | rel Freq% | QMD | rel DOM% |
|---|---|---|---|---|---|---|---|---|---|
| # allWTs | 254 | red alder | 40 | 3to20 | 8.7 | 8 | 15.75 | 9.6 | 5.02 |
| all avDiam | 13.0 | ash | 2 | 14to15 | 14.5 | 14.5 | 0.79 | 14.5 | 0.57 |
| all QMD | 17.0 | barberry | 1 | 5 | 5 | 5 | 0.39 | 5 | 0.03 |
| SrvyYr | 1872 | redcedar | 13 | 3to40 | 23.8 | 24 | 5.12 | 25.9 | 11.87 |
| | | cherry | 1 | 4 | 4 | 4 | 0.39 | 4 | 0.02 |
| | | cottonwood | 3 | 3to30 | 14.3 | 10 | 1.18 | 18.3 | 1.37 |
| | | dogwood | 1 | 3 | 3 | 3 | 0.39 | 3 | 0.01 |
| | | Douglas-fir | 129 | 3to80 | 14.9 | 12 | 50.79 | 19.7 | 68.15 |
| | | hemlock | 37 | 3to30 | 13.2 | 12 | 14.57 | 15.3 | 11.79 |
| | | bl maple | 16 | 3to10 | 5.5 | 4.5 | 6.30 | 6 | 0.78 |
| | | vine maple | 11 | 3to80 | 4.6 | 4 | 4.33 | 5 | 0.37 |

| Twp-Range | 21-7e | species | # WTs | Diam Range | av Diam | Median Diam | rel Freq% | QMD | rel DOM% |
|---|---|---|---|---|---|---|---|---|---|
| # allWTs | 287 | red alder | 25 | 5to24 | 10.8 | 10 | 8.71 | 11.8 | 5.52 |
| all avDiam | 11.6 | redcedar | 4 | 8to36 | 17.3 | 12.5 | 1.39 | 20.5 | 2.67 |
| all QMD | 14.8 | cherry | 8 | 3to7 | 5.4 | 5.5 | 2.79 | 5.5 | 0.38 |
| SrvyYr | 1881 | cottonwood | 1 | 12 | 12 | 12 | 0.35 | 12 | 0.23 |
| | | Douglas-fir | 187 | 3to60 | 11.4 | 9 | 65.16 | 15 | 66.72 |
| | | grand fir | 2 | 10to12 | 11 | 11 | 0.70 | 11 | 0.38 |
| | | hemlock | 39 | 4to48 | 14.7 | 10 | 13.59 | 17.7 | 19.38 |
| | | bl maple | 5 | 3to12 | 7 | 5 | 1.74 | 7.8 | 0.48 |
| | | pine | 1 | 5 | 5 | 5 | 0.35 | 5 | 0.04 |
| | | spruce | 4 | 20to24 | 23 | 24 | 1.39 | 23.1 | 3.38 |
| | | vine maple | 8 | 4to10 | 6 | 5.5 | 2.79 | 6.4 | 0.52 |
| | | willow | 3 | 3to7 | 5.3 | 6 | 1.05 | 5.6 | 0.15 |

| Twp-Range | 21-1w | species | # WTs | Diam Range | av Diam | Median Diam | rel Freq% | QMD | rel DOM% |
|---|---|---|---|---|---|---|---|---|---|
| # allWTs | 151 | red alder | 9 | 4to20 | 8.3 | 6 | 5.96 | 9.6 | 1.01 |
| all avDiam | 19.5 | redcedar | 28 | 9to40 | 22.6 | 21.5 | 18.54 | 24 | 19.61 |
| all QMD | 23.3 | cherry | 1 | 6 | 6 | 6 | 0.66 | 6 | 0.04 |
| SrvyYr | 1856 | dogwood | 1 | 8 | 8 | 8 | 0.66 | 8 | 0.08 |
| | | Douglas-fir | 93 | 4to70 | 22.7 | 20 | 61.59 | 26.3 | 78.23 |
| | | hemlock | 6 | 5to20 | 9 | 8 | 3.97 | 10.3 | 0.77 |
| | | madrone | 7 | 3to6 | 3.6 | 3 | 4.64 | 3.7 | 0.12 |
| | | sweetwood | 1 | 3 | 3 | 3 | 0.66 | 3 | 0.01 |
| | | thorn | 3 | 4to12 | 8.7 | 4 | 1.99 | 9.3 | 0.32 |
| | | willow | 1 | 3 | 3 | 3 | 0.66 | 3 | 0.01 |
| | | yew | 1 | 10 | 10 | 10 | 0.66 | 10 | 0.12 |

| Twp-Range | 21-2w | species | # WTs | Diam Range | av Diam | Median Diam | rel Freq% | QMD | rel DOM% |
|---|---|---|---|---|---|---|---|---|---|
| # allWTs | 262 | red alder | 13 | 4to20 | 6.2 | 4 | 4.96 | 7.5 | 0.68 |
| all avDiam | 16.6 | redcedar | 28 | 4to50 | 19.3 | 18 | 10.69 | 21.8 | 12.39 |
| all QMD | 20.2 | Douglas-fir | 180 | 4to80 | 18.8 | 14 | 68.70 | 22.3 | 83.37 |
| SrvyYr | 1858 | hemlock | 16 | 4to20 | 10.8 | 10 | 6.11 | 11.8 | 2.07 |
| | | bl maple | 3 | 5to16 | 9.7 | 8 | 1.15 | 10.7 | 0.32 |
| | | madrone | 8 | 4to25 | 8 | 4 | 3.05 | 10.8 | 0.87 |
| | | pine | 9 | 4to10 | 6.2 | 6 | 3.44 | 6.6 | 0.37 |
| | | willow | 5 | 4to50 | 4.2 | 4 | 1.91 | 4.2 | 0.08 |

| Twp-Range | 21-3w | species | # WTs | Diam Range | av Diam | Median Diam | rel Freq% | QMD | rel DOM% |
|---|---|---|---|---|---|---|---|---|---|
| # allWTs | 281 | red alder | 12 | 4to14 | 7.9 | 7 | 4.27 | 8.8 | 1.48 |
| all avDiam | 13.1 | ash | 1 | 10 | 10 | 10 | 0.36 | 10 | 0.16 |
| all QMD | 14.9 | redcedar | 6 | 6to25 | 13.7 | 14 | 2.14 | 15 | 2.15 |
| SrvyYr | 1857 | crabapple | 3 | 5to20 | 10 | 5 | 1.07 | 12.2 | 0.71 |
| | | dogwood | 1 | 6 | 6 | 6 | 0.36 | 6 | 0.06 |
| | | Douglas-fir | 195 | 4to40 | 15 | 14 | 69.40 | 16.6 | 85.62 |
| | | hemlock | 25 | 6to25 | 12 | 10 | 8.90 | 13 | 6.73 |
| | | madrone | 4 | 4to6 | 4.8 | 4.5 | 1.42 | 4.8 | 0.15 |
| | | pine | 32 | 4to14 | 6.6 | 6 | 11.39 | 7.2 | 2.64 |
| | | willow | 2 | 5to5 | 5.5 | 5.5 | 0.71 | 5.5 | 0.10 |

| Twp-Range | 21-4w | species | # WTs | Diam Range | av Diam | Median Diam | rel Freq% | QMD | rel DOM% |
|---|---|---|---|---|---|---|---|---|---|
| # allWTs | 267 | red alder | 26 | 4to20 | 11.7 | 12 | 9.74 | 12.9 | 3.35 |
| all avDiam | 18.0 | ocean spray | 1 | 4 | 4 | 4 | 0.37 | 4 | 0.01 |
| all QMD | 22.0 | barberry | 1 | 4 | 4 | 4 | 0.37 | 4 | 0.01 |
| SrvyYr | 1861 | redcedar | 10 | 10to80 | 27.2 | 21 | 3.75 | 33.7 | 8.80 |
| | | crabapple | 2 | 5to16 | 10.5 | 10.5 | 0.75 | 11.9 | 0.22 |
| | | cottonwood | 3 | 6to30 | 16 | 12 | 1.12 | 19 | 0.84 |
| | | dogwood | 1 | 6 | 6 | 6 | 0.37 | 6 | 0.03 |
| | | Douglas-fir | 143 | 6to60 | 23.9 | 20 | 53.56 | 26.7 | 78.98 |
| | | hemlock | 9 | 5to20 | 9.3 | 6 | 3.37 | 10.5 | 0.77 |
| | | hazel | 1 | 4 | 4 | 4 | 0.37 | 4 | 0.01 |
| | | bl maple | 9 | 4to36 | 18.4 | 18 | 3.37 | 21.4 | 3.19 |
| | | pine | 39 | 4to20 | 9.2 | 8 | 14.61 | 9.8 | 2.90 |
| | | vine maple | 18 | 4to7 | 5.3 | 5 | 6.74 | 5.3 | 0.39 |
| | | willow | 4 | 4to20 | 8.8 | 5.5 | 1.50 | 10.9 | 0.37 |

| Twp-Range | 21-5w | species | # WTs | Diam Range | av Diam | Median Diam | rel Freq% | QMD | rel DOM% |
|---|---|---|---|---|---|---|---|---|---|
| # allWTs | 278 | red alder | 8 | 5to24 | 12.8 | 12 | 2.88 | 14.1 | 1.47 |
| all avDiam | 16.2 | redcedar | 14 | 4to80 | 25.1 | 19 | 5.04 | 31.4 | 12.76 |
| all QMD | 19.7 | cascara | 1 | 6 | 6 | 6 | 0.36 | 6 | 0.03 |
| SrvyYr | 1875 | crabapple | 2 | 5to8 | 6.5 | 6.5 | 0.72 | 6.7 | 0.08 |
| | | dogwood | 2 | 5to7 | 6 | 6 | 0.72 | 61 | 6.88 |
| | | Douglas-fir | 132 | 3to50 | 21 | 20 | 47.48 | 24.1 | 70.86 |
| | | hemlock | 69 | 4to35 | 11.5 | 10 | 24.82 | 12.9 | 10.61 |
| | | hazel | 2 | 3 | 3 | 3 | 0.72 | 3 | 0.02 |
| | | bl maple | 10 | 3to36 | 14 | 13 | 3.60 | 17.4 | 2.80 |
| | | pine | 10 | 5to20 | 9.3 | 8 | 3.60 | 10.2 | 0.96 |
| | | vine maple | 28 | 2to7 | 4.1 | 4 | 10.07 | 4.2 | 0.46 |

| Twp-Range | 20-1e | species | # WTs | Diam Range | av Diam | Median Diam | rel Freq% | QMD | rel DOM% |
|---|---|---|---|---|---|---|---|---|---|
| # allWTs | 99 | red alder | 10 | 4to14 | 9.2 | 10 | 10.10 | 9.6 | 2.98 |
| all avDiam | 15.3 | redcedar | 16 | 6to40 | 17 | 13 | 16.16 | 19.6 | 19.90 |
| all QMD | 17.7 | Douglas-fir | 70 | 4to40 | 16.1 | 14 | 70.71 | 18.3 | 75.88 |
| SrvyYr | 1853 | madrone | 2 | 10 | 10 | 10 | 2.02 | 10 | 0.65 |
| | | willow | 1 | 8 | 8 | 8 | 1.01 | 8 | 0.21 |

| Twp-Range | 20-2e | species | # WTs | Diam Range | av Diam | Median Diam | rel Freq% | QMD | rel DOM% |
|---|---|---|---|---|---|---|---|---|---|
| # allWTs | 194 | red alder | 7 | 4to12 | 8.1 | 8 | 3.61 | 8.6 | 0.48 |
| all avDiam | 20.2 | redcedar | 13 | 5to40 | 19.5 | 20 | 6.70 | 21.7 | 5.71 |
| all QMD | 23.5 | Douglas-fir | 128 | 3to50 | 22.9 | 24 | 65.98 | 25.7 | 78.83 |
| SrvyYr | 1868 | hemlock | 1 | 12 | 12 | 12 | 0.52 | 12 | 0.13 |
| | | bl maple | 1 | 4 | 4 | 4 | 0.52 | 4 | 0.01 |
| | | oak | 40 | 3to40 | 16.3 | 15 | 20.62 | 19.7 | 14.48 |
| | | willow | 4 | 3to10 | 5.5 | 4.5 | 2.06 | 6.2 | 0.14 |

| Twp-Range | 20-3e | species | # WTs | Diam Range | av Diam | Median Diam | rel Freq% | QMD | rel DOM% |
|---|---|---|---|---|---|---|---|---|---|
| # allWTs | 255 | red alder | 17 | 4to20 | 9.5 | 10 | 6.67 | 10.4 | 1.18 |
| all avDiam | 21.2 | ash | 9 | 10to24 | 15.2 | 12 | 3.53 | 15.9 | 1.47 |
| all QMD | 24.7 | barberry | 1 | 5 | 5 | 5 | 0.39 | 5 | 0.02 |
| SrvyYr | 1866 | redcedar | 16 | 4to50 | 20.5 | 18 | 6.27 | 23.4 | 5.65 |
| | &73 | crabapple | 4 | 4to7 | 5.3 | 5 | 1.57 | 5.4 | 0.08 |
| | | cottonwood | 8 | 3to70 | 19.8 | 9 | 3.14 | 29.7 | 4.55 |
| | | Douglas-fir | 170 | 4to80 | 24.1 | 24 | 66.67 | 26.8 | 78.68 |
| | | hemlock | 4 | 6to18 | 12.8 | 13.5 | 1.57 | 13.5 | 0.47 |
| | | hazel | 2 | 3to15 | 9 | 9 | 0.78 | 10.8 | 0.15 |
| | | bl maple | 1 | 4 | 4 | 4 | 0.39 | 4 | 0.01 |
| | | oak | 12 | 12to36 | 23.8 | 24 | 4.71 | 24.7 | 4.72 |
| | | spruce | 2 | 10to18 | 14 | 14 | 0.78 | 14.6 | 0.27 |
| | | vine maple | 1 | 3 | 3 | 3 | 0.39 | 3 | 0.01 |
| | | willow | 7 | 3to60 | 14.4 | 4 | 2.75 | 24 | 2.60 |
| | | yew | 1 | 7 | 7 | 7 | 0.39 | 7 | 0.03 |

| Twp-Range | 20-4e | species | # WTs | Diam Range | av Diam | Median Diam | rel Freq% | QMD | rel DOM% |
|---|---|---|---|---|---|---|---|---|---|
| # allWTs | 284 | red alder | 51 | 3to30 | 10.3 | 6 | 17.96 | 12.3 | 6.98 |
| all avDiam | 14.9 | ash | 12 | 8to24 | 16.3 | 16.5 | 4.23 | 17.2 | 3.21 |
| ttl QMD | 19.7 | barberry | 1 | 5 | 5 | 5 | 0.35 | 5 | 0.02 |
| SrvyYr | 1864&73 | redcedar | 38 | 3to108 | 20.9 | 18 | 13.38 | 27.9 | 26.77 |
| | | crabapple | 12 | 3to24 | 8.1 | 4 | 4.23 | 10.4 | 1.17 |
| | | cottonwood | 17 | 4to42 | 19.7 | 14 | 5.99 | 24 | 8.86 |
| | | dogwood | 1 | 6 | 6 | 6 | 0.35 | 6 | 0.03 |
| | | Douglas-fir | 72 | 3to54 | 21.8 | 20 | 25.35 | 25.6 | 42.71 |
| | | hemlock | 11 | 6to46 | 13.9 | 12 | 3.87 | 17.7 | 3.12 |
| | | hazel | 3 | 3to7 | 4.3 | 3 | 1.06 | 4.7 | 0.06 |
| | | bl maple | 31 | 3to18 | 7.4 | 7 | 10.92 | 8 | 1.80 |
| | | spruce | 4 | 18to36 | 28.5 | 30 | 1.41 | 29.5 | 3.15 |
| | | vine maple | 3 | 4to7 | 7 | 7 | 1.06 | 7.4 | 0.15 |
| | | willow | 28 | 2to24 | 6.4 | 3 | 9.86 | 8.6 | 1.87 |

| Twp-Range | 20-5e | species | # WTs | Diam Range | av Diam | Median Diam | rel Freq% | QMD | rel DOM% |
|---|---|---|---|---|---|---|---|---|---|
| # allWTs | 258 | red alder | 41 | 3to30 | 10.4 | 8 | 15.89 | 12.5 | 5.01 |
| all avDiam | 16.8 | ash | 10 | 7to24 | 10.7 | 10 | 3.88 | 12 | 1.13 |
| all QMD | 22.3 | barberry | 1 | 5 | 5 | 5 | 0.39 | 5 | 0.02 |
| SrvyYr | 1872 | redcedar | 38 | 5to66 | 25.7 | 22 | 14.73 | 29 | 25.01 |
| | | cherry | 6 | 4to12 | 6.7 | 6 | 2.33 | 7.2 | 0.24 |
| | | crabapple | 4 | 5to12 | 8.3 | 7.5 | 1.55 | 8.4 | 0.22 |
| | | cottonwood | 12 | 6to36 | 15.2 | 12.5 | 4.65 | 18 | 3.04 |
| | | dogwood | 2 | 4to5 | 4.5 | 4.5 | 0.78 | 4.5 | 0.03 |
| | | Douglas-fir | 109 | 4to96 | 20.8 | 16 | 42.25 | 27.1 | 62.65 |
| | | hemlock | 8 | 4to20 | 10.5 | 10 | 3.10 | 11.5 | 0.83 |
| | | bl maple | 9 | 4to30 | 10.4 | 5 | 3.49 | 13.7 | 1.32 |
| | | vine maple | 1 | 7 | 7 | 7 | 0.39 | 7 | 0.04 |
| | | willow | 17 | 2to15 | 5.8 | 5 | 6.59 | 6.7 | 0.60 |

| Twp-Range | 20-6e | species | # WTs | Diam Range | av Diam | Median Diam | rel Freq% | QMD | rel DOM% |
|---|---|---|---|---|---|---|---|---|---|
| # allWTs | 296 | red alder | 71 | 3to22 | 7.8 | 7 | 23.99 | 8.7 | 6.58 |
| all avDiam | 11.0 | ash | 5 | 4to24 | 11.6 | 6 | 1.69 | 14.4 | 1.27 |
| all QMD | 16.6 | aspen | 3 | 4to6 | 5.3 | 6 | 1.01 | 5.4 | 0.11 |
| SrvyYr | 1882 | barberry | 1 | 6 | 6 | 6 | 0.34 | 6 | 0.04 |
| | | redcedar | 23 | 10to66 | 27.5 | 24 | 7.77 | 32.4 | 29.58 |
| | | cherry | 17 | 4to10 | 5.7 | 5 | 5.74 | 5.9 | 0.72 |
| | | cottonwood | 23 | 3to18 | 8 | 8 | 7.77 | 8.9 | 2.23 |
| | | dogwood | 1 | 8 | 8 | 8 | 0.34 | 8 | 0.08 |
| | | Douglas-fir | 110 | 3to96 | 12.2 | 7 | 37.16 | 20.1 | 54.44 |
| | | hemlock | 6 | 3to18 | 10.7 | 9.5 | 2.03 | 11.1 | 0.91 |
| | | bl maple | 12 | 5to24 | 10.2 | 8 | 4.05 | 11.6 | 1.98 |
| | | spruce | 3 | 7to8 | 7.7 | 8 | 1.01 | 7.7 | 0.22 |
| | | willow | 21 | 3to8 | 5.6 | 6 | 7.09 | 5.7 | 0.84 |

| Twp-Range | 20-7e | species | # WTs | Diam Range | av Diam | Median Diam | rel Freq% | QMD | rel DOM% |
|---|---|---|---|---|---|---|---|---|---|
| # allWTs | 291 | red alder | 16 | 3to20 | 6.8 | 5.5 | 5.50 | 7.8 | 1.36 |
| all avDiam | 12.7 | redcedar | 22 | 5to48 | 14.6 | 11 | 7.56 | 15.6 | 7.50 |
| all QMD | 15.7 | cherry | 3 | 3to5 | 4 | 4 | 1.03 | 4.1 | 0.07 |
| SrvyYr | 1881 | Douglas-fir | 59 | 4to36 | 15.4 | 10 | 20.27 | 18.8 | 29.20 |
| | | hemlock | 168 | 3to48 | 12.8 | 10 | 57.73 | 15.2 | 54.35 |
| | | bl maple | 7 | 4to12 | 7.4 | 6 | 2.41 | 8.1 | 0.64 |
| | | spruce | 5 | 4to36 | 19.2 | 18 | 1.72 | 22.2 | 3.45 |
| | | vine maple | 3 | 3to6 | 5 | 6 | 1.03 | 5.2 | 0.11 |
| | | willow | 8 | 3to6 | 4.6 | 4 | 2.75 | 4.8 | 0.26 |

| Twp-Range | 20-1w | species | # WTs | Diam Range | av Diam | Median Diam | rel Freq% | QMD | rel DOM% |
|---|---|---|---|---|---|---|---|---|---|
| # allWTs | 160 | red alder | 3 | 8to12 | 10 | 10 | 1.88 | 10.1 | 0.71 |
| all avDiam | 14.1 | (none) | | | | | 0.00 | | 0.00 |
| all QMD | 16.4 | redcedar | 17 | 6to36 | 15.3 | 18 | 10.63 | 17.1 | 11.54 |
| SrvyYr | 1853 | cherry | 1 | 6 | 6 | 6 | 0.63 | 6 | 0.08 |
| | | dogwood | 1 | 8 | 8 | 8 | 0.63 | 8 | 0.15 |
| | | Douglas-fir | 120 | 2to40 | 14.5 | 12 | 75.00 | 16.9 | 79.55 |
| | | hemlock | 10 | 3to20 | 10.7 | 9 | 6.25 | 12.2 | 3.45 |
| | | bl maple | 4 | 6to24 | 12 | 9 | 2.50 | 13.9 | 1.79 |
| | | madrone | 3 | 10to16 | 16 | 16 | 1.88 | 14.3 | 1.42 |
| | | yew | 1 | 20 | 20 | 20 | 0.63 | 20 | 0.93 |

| Twp-Range | 20-2w | species | # WTs | Diam Range | av Diam | Median Diam | rel Freq% | QMD | rel DOM% |
|---|---|---|---|---|---|---|---|---|---|
| # allWTs | 211 | red alder | 5 | 6to12 | 9.6 | 10 | 2.37 | 9.8 | 0.69 |
| all avDiam | 15.4 | ash | 1 | 24 | 24 | 24 | 0.47 | 24 | 0.83 |
| all QMD | 18.1 | redcedar | 30 | 3to40 | 17.9 | 12 | 14.22 | 20.9 | 18.89 |
| SrvyYr | 1853 | crabapple | 1 | 8 | 8 | 8 | 0.47 | 8 | 0.09 |
| | | dogwood | 1 | 10 | 10 | 10 | 0.47 | 10 | 0.14 |
| | | Douglas-fir | 147 | 4to60 | 15.8 | 12 | 69.67 | 18.6 | 73.29 |
| | | hemlock | 19 | 4to24 | 12.3 | 12 | 9.00 | 13.3 | 4.84 |
| | | bl maple | 1 | 18 | 18 | 18 | 0.47 | 18 | 0.47 |
| | | madrone | 2 | 8to14 | 11 | 11 | 0.95 | 11.4 | 0.37 |
| | | thorn | 1 | 6 | 6 | 6 | 0.47 | 6 | 0.05 |
| | | willow | 2 | 6to10 | 8 | 8 | 0.95 | 8.2 | 0.19 |
| | | yew | 1 | 12 | 12 | 12 | 0.47 | 12 | 0.21 |

| Twp-Range | 20-3w | species | # WTs | Diam Range | av Diam | Median Diam | rel Freq% | QMD | rel DOM% |
|---|---|---|---|---|---|---|---|---|---|
| # allWTs | 254 | red alder | 15 | 4to22 | 10.9 | 10 | 5.91 | 12.2 | 2.80 |
| all avDiam | 15.4 | ash | 2 | 16 | 16 | | 0.79 | 16.5 | 0.68 |
| all QMD | 17.7 | barberry | 1 | 5 | 5 | 5 | 0.39 | 5 | 0.03 |
| SrvyYr | 1855 | redcedar | 43 | 5to35 | 16.3 | 14 | 16.93 | 18.1 | 17.66 |
| | | dogwood | 2 | 6 | 6 | 6 | 0.79 | 6 | 0.09 |
| | | Douglas-fir | 126 | 5to48 | 17.9 | 15 | 49.61 | 20.1 | 63.82 |
| | | hemlock | 42 | 3to50 | 11.7 | 10 | 16.54 | 14 | 10.32 |
| | | bl maple | 5 | 4to22 | 13.6 | 14 | 1.97 | 15 | 1.41 |
| | | madrone | 3 | 8to30 | 16 | 10 | 1.18 | 18.8 | 1.33 |
| | | oak | 1 | 10 | 10 | 10 | 0.39 | 10 | 0.13 |
| | | pine | 14 | 4to15 | 8.7 | 8 | 5.51 | 9.6 | 1.62 |

| Twp-Range | 20-4w | species | # WTs | Diam Range | av Diam | Median Diam | rel Freq% | QMD | rel DOM% |
|---|---|---|---|---|---|---|---|---|---|
| # allWTs | 280 | red alder | 20 | 4to24 | 11 | 12 | 7.14 | 11.9 | 1.90 |
| all avDiam | 18.1 | barberry | 2 | 4to5 | 4.5 | 4.5 | 0.71 | 4.5 | 0.03 |
| all QMD | 23.1 | redcedar | 33 | 5to60 | 23.9 | 20 | 11.79 | 28.8 | 18.39 |
| SrvyYr | 1861 | crabapple | 3 | 6to18 | 11 | 9 | 1.07 | 12.1 | 0.30 |
| | &72 | dogwood | 2 | 6to12 | 9 | 9 | 0.71 | 9.5 | 0.12 |
| | | Douglas-fir | 120 | 4to65 | 25.7 | 24 | 42.86 | 29.6 | 70.66 |
| | | hemlock | 53 | 3to36 | 10.7 | 10 | 18.93 | 12.8 | 5.84 |
| | | hazel | 2 | 4to5 | 4.5 | 4.5 | 0.71 | 4.5 | 0.03 |
| | | bl maple | 8 | 4to36 | 11.5 | 8 | 2.86 | 15 | 1.21 |
| | | oak | 2 | 14to20 | 17 | 17 | 0.71 | 17.3 | 0.40 |
| | | pine | 13 | 4to12 | 8.2 | 8 | 4.64 | 8.5 | 0.63 |
| | | pigeonwood | 1 | 4 | 4 | 4 | 0.36 | 4 | 0.01 |
| | | cascara | 2 | 5 | 5 | 5 | 0.71 | 5 | 0.03 |
| | | vine maple | 17 | 4to65 | 4.7 | 5 | 6.07 | 4.8 | 0.26 |
| | | willow | 2 | 5to60 | 5.5 | 5.5 | 0.71 | 5.5 | 0.04 |

| Twp-Range | 20-5w | species | # WTs | Diam Range | av Diam | Median Diam | rel Freq% | QMD | rel DOM% |
|---|---|---|---|---|---|---|---|---|---|
| # allWTs | 282 | red alder | 8 | 3to14 | 8.8 | 10 | 2.84 | 9.4 | 0.52 |
| all avDiam | 17.7 | redcedar | 51 | 4to50 | 21.6 | 20 | 18.09 | 24.8 | 23.16 |
| all QMD | 21.9 | cherry | 1 | 14 | 14 | 14 | 0.35 | 14 | 0.14 |
| SrvyYr | 1875 | cascara | 7 | 4to8 | 6 | 6 | 2.48 | 6.2 | 0.20 |
| | | crabapple | 1 | 4 | 4 | 4 | 0.35 | 4 | 0.01 |
| | | dogwood | 1 | 4 | 4 | 4 | 0.35 | 4 | 0.01 |
| | | Douglas-fir | 80 | 3to60 | 26.5 | 27 | 28.37 | 29.9 | 52.80 |
| | | hemlock | 113 | 3to84 | 13 | 10 | 40.07 | 16.1 | 21.63 |
| | | bl maple | 2 | 4to8 | 6 | 6 | 0.71 | 6.3 | 0.06 |
| | | pine | 6 | 4to18 | 12.3 | 12 | 2.13 | 13.2 | 0.77 |
| | | spruce | 2 | 10to16 | 13 | 13 | 0.71 | 14.6 | 0.31 |
| | | vine maple | 7 | 3to8 | 5 | 5 | 2.48 | 5.3 | 0.15 |
| | | willow | 3 | 5to10 | 7.7 | 8 | 1.06 | 7.9 | 0.14 |

| Twp-Range | 20-6w | species | # WTs | Diam Range | av Diam | Median Diam | rel Freq% | QMD | rel DOM% |
|---|---|---|---|---|---|---|---|---|---|
| # allWTs | 290 | red alder | 10 | 4to18 | 10.4 | 11 | 3.45 | 11.2 | 0.78 |
| all avDiam | 18.4 | redcedar | 13 | 10to36 | 18.5 | 10 | 4.48 | 19.9 | 3.19 |
| all QMD | 23.6 | cascara | 10 | 3to10 | 5.5 | 5 | 3.45 | 5.9 | 0.22 |
| SrvyYr | 1875 | crabapple | 1 | 10 | 10 | 10 | 0.34 | 10 | 0.06 |
| | | dogwood | 4 | 4to8 | 5.8 | 5.5 | 1.38 | 5.9 | 0.09 |
| | | Douglas-fir | 122 | 3to96 | 27 | 24 | 42.07 | 32.2 | 78.44 |
| | | grand fir | 3 | 6to18 | 11.3 | 10 | 1.03 | 12.4 | 0.29 |
| | | hemlock | 84 | 3to40 | 15 | 12 | 28.97 | 16.8 | 14.70 |
| | | hazel | 1 | 3 | 3 | 3 | 0.34 | 3 | 0.01 |
| | | bl maple | 8 | 4to18 | 7.8 | 6.6 | 2.76 | 8.9 | 0.39 |
| | | pigeonwood | 1 | 7 | 7 | 7 | 0.34 | 7 | 0.03 |
| | | spruce | 5 | 12to30 | 21.4 | 24 | 1.72 | 22.4 | 1.56 |
| | | vine maple | 28 | 3to8 | 5 | 5 | 9.66 | 5.2 | 0.47 |

| Twp-Range | 19-1e | species | # WTs | Diam Range | av Diam | Median Diam | rel Freq% | QMD | rel DOM% |
|---|---|---|---|---|---|---|---|---|---|
| # allWTs | 126 | red alder | 2 | 18to24 | 21 | 21 | 1.59 | 21.2 | 2.28 |
| all avDiam | 15.4 | ash | 3 | 10to30 | 18 | 14 | 2.38 | 20 | 3.04 |
| all QMD | 17.7 | barberry | 1 | 8 | 8 | 8 | 0.79 | 8 | 0.16 |
| SrvyYr | 1853 | redcedar | 9 | 12to30 | 19.6 | 20 | 7.14 | 20.9 | 9.96 |
| | | crabapple | 1 | 10 | 10 | 10 | 0.79 | 10 | 0.25 |
| | | dogwood | 1 | 8 | 8 | 8 | 0.79 | 8 | 0.16 |
| | | Douglas-fir | 78 | 3to40 | 15.6 | 12 | 61.90 | 18 | 64.01 |
| | | hemlock | 4 | 4to16 | 10.5 | 11 | 3.17 | 11.4 | 1.32 |
| | | madrone | 5 | 6to20 | 11.2 | 10 | 3.97 | 12.1 | 1.85 |
| | | oak | 16 | 3to30 | 17.2 | 18 | 12.70 | 19.5 | 15.41 |
| | | pine | 3 | 8to12 | 10 | 10 | 2.38 | 10.2 | 0.79 |
| | | thorn | 1 | 12 | 12 | 12 | 0.79 | 12 | 0.36 |
| | | willow | 2 | 6to10 | 8 | 8 | 1.59 | 8.2 | 0.34 |

| Twp-Range | 19-2e | species | # WTs | Diam Range | av Diam | Median Diam | rel Freq% | QMD | rel DOM% |
|---|---|---|---|---|---|---|---|---|---|
| # allWTs | 188 | cottonwood | 2 | 6to10 | 8 | 8 | 1.06 | 8.2 | 0.13 |
| all avDiam | 21.2 | Douglas-fir | 96 | 6to60 | 20.3 | 18 | 51.06 | 23 | 47.61 |
| all QMD | 23.8 | hemlock | 1 | 22 | 22 | 22 | 0.53 | 22 | 0.45 |
| SrvyYr | 1870 | oak | 87 | 4to48 | 22.5 | 20 | 46.28 | 24.9 | 50.57 |
| | | pine | 1 | 36 | 36 | 36 | 0.53 | 36 | 1.22 |
| | | willow | 1 | 7 | 7 | 7 | 0.53 | 7 | 0.05 |

| Twp-Range | 19-3e | species | # WTs | Diam Range | av Diam | Median Diam | rel Freq% | QMD | rel DOM% |
|---|---|---|---|---|---|---|---|---|---|
| # allWTs | 190 | red alder | 3 | 5to6 | 5.7 | 6 | 1.58 | 5.7 | 0.11 |
| all avDiam | 18.7 | ash | 3 | 10to20 | 16 | 18 | 1.58 | 16.6 | 0.97 |
| all QMD | 21.2 | redcedar | 24 | 6to38 | 18.5 | 14.5 | 12.63 | 20.9 | 12.33 |
| SrvyYr | 1872 | cottonwood | 2 | 7to24 | 15.5 | 15.5 | 1.05 | 17.7 | 0.74 |
| | | Douglas-fir | 121 | 3to48 | 20.2 | 20 | 63.68 | 22.5 | 72.05 |
| | | hemlock | 1 | 16 | 16 | 16 | 0.53 | 16 | 0.30 |
| | | bl maple | 1 | 6 | 6 | 6 | 0.53 | 6 | 0.04 |
| | | oak | 31 | 3to36 | 16.8 | 16 | 16.32 | 19.3 | 13.58 |
| | | vine maple | 2 | 3 | 3 | 3 | 1.05 | 3 | 0.02 |
| | | willow | 2 | 3to10 | 6.5 | 6.5 | 1.05 | 7.4 | 0.13 |

| Twp-Range | 19-4e | species | # WTs | Diam Range | av Diam | Median Diam | rel Freq% | QMD | rel DOM% |
|---|---|---|---|---|---|---|---|---|---|
| # allWTs | 267 | red alder | 10 | 3to30 | 10.8 | 9 | 3.75 | 13.4 | 1.34 |
| all avDiam | 19.0 | ash | 4 | 4to20 | 12 | 12 | 1.50 | 13.9 | 0.58 |
| all QMD | 22.4 | redcedar | 40 | 2to60 | 20.2 | 16 | 14.98 | 23 | 15.77 |
| SrvyYr | 1872 | cherry | 1 | 11 | 11 | 11 | 0.37 | 11 | 0.09 |
| | | crabapple | 1 | 7 | 7 | 7 | 0.37 | 7 | 0.04 |
| | | cottonwood | 2 | 10to12 | 11 | 11 | 0.75 | 11 | 0.18 |
| | | Douglas-fir | 184 | 3to60 | 20.8 | 16 | 68.91 | 24.1 | 79.63 |
| | | hemlock | 16 | 6to22 | 10.3 | 9.5 | 5.99 | 11.2 | 1.50 |
| | | bl maple | 4 | 6to18 | 12.3 | 12.5 | 1.50 | 13 | 0.50 |
| | | vine maple | 3 | 5to6 | 5.7 | 6 | 1.12 | 5.7 | 0.07 |
| | | willow | 2 | 4to6 | 5 | 5 | 0.75 | 5.1 | 0.04 |

| Twp-Range | 19-5e | species | # WTs | Diam Range | av Diam | Median Diam | rel Freq% | QMD | rel DOM% |
|---|---|---|---|---|---|---|---|---|---|
| # allWTs | 262 | red alder | 14 | 5to36 | 13.6 | 10 | 5.32 | 16.8 | 3.46 |
| all avDiam | 17.4 | ash | 6 | 3to14 | 7.3 | 6 | 2.28 | 8.2 | 0.35 |
| all QMD | 20.8 | aspen | 2 | 4to5 | 4.5 | 4.5 | 0.76 | 4.5 | 0.04 |
| SrvyYr | 1873 | barberry | 5 | 3to12 | 6.6 | 5 | 1.90 | 7.3 | 0.23 |
| | | redcedar | 42 | 4to48 | 18.5 | 18 | 15.97 | 20.7 | 15.77 |
| | | cherry | 2 | 5to8 | 6.5 | 6.5 | 0.76 | 6.7 | 0.08 |
| | | cottonwood | 5 | 6to60 | 19.6 | 12 | 1.90 | 28.2 | 3.49 |
| | | dogwood | 2 | 3to4 | 3.5 | 3.5 | 0.76 | 3.5 | 0.02 |
| | | Douglas-fir | 135 | 3to50 | 20.8 | 20 | 51.33 | 24 | 68.15 |
| | | hemlock | 28 | 4to31 | 13.1 | 12 | 10.65 | 14.7 | 5.30 |
| | | bl maple | 10 | 4to24 | 16 | 16.5 | 3.80 | 17.3 | 2.62 |
| | | vine maple | 8 | 4to8 | 5.3 | 5 | 3.04 | 5.4 | 0.20 |
| | | willow | 3 | 4to8 | 6.7 | 8 | 1.14 | 6.9 | 0.13 |

| Twp-Range | 19-6e | species | # WTs | Diam Range | av Diam | Median Diam | rel Freq% | QMD | rel DOM% |
|---|---|---|---|---|---|---|---|---|---|
| # allWTs | 266 | red alder | 15 | 4to18 | 9.4 | 9 | 5.64 | 9.9 | 2.00 |
| all avDiam | 12.3 | ash | 2 | 10to12 | 11 | 11 | 0.75 | 11 | 0.33 |
| all QMD | 16.6 | redcedar | 62 | 4to66 | 18.2 | 11.5 | 23.31 | 23.9 | 48.26 |
| SrvyYr | 1873 | cherry | 3 | 6to10 | 7.7 | 7 | 1.13 | 7.9 | 0.26 |
| | | crabapple | 2 | 4to12 | 8 | 8 | 0.75 | 8.9 | 0.22 |
| | | cottonwood | 2 | 3to4 | 3.5 | 3.5 | 0.75 | 3.5 | 0.03 |
| | | dogwood | 3 | 4to5 | 4 | 4 | 1.13 | 4.1 | 0.07 |
| | | Douglas-fir | 29 | 3to40 | 15.1 | 12 | 10.90 | 18.1 | 12.95 |
| | | grand fir | 1 | 24 | 24 | 24 | 0.38 | 24 | 0.78 |
| | | hemlock | 109 | 3to48 | 9.5 | 8 | 40.98 | 11.8 | 20.68 |
| | | bl maple | 12 | 3to18 | 7.6 | 5.5 | 4.51 | 8.9 | 1.30 |
| | | spruce | 10 | 9to45 | 27.2 | 30 | 3.76 | 30.3 | 12.51 |
| | | vine maple | 15 | 3to6 | 4.1 | 4 | 5.64 | 4.2 | 0.36 |
| | | willow | 1 | 3 | 3 | 3 | 0.38 | 3 | 0.01 |

| Twp-Range | 19-7e | species | # WTs | Diam Range | av Diam | Median Diam | rel Freq% | QMD | rel DOM% |
|---|---|---|---|---|---|---|---|---|---|
| # allWTs | 273 | red alder | 3 | 8to10 | 9.3 | 10 | 1.10 | 9.4 | 0.22 |
| all avDiam | 17.9 | redcedar | 29 | 6to50 | 27.6 | 26 | 10.62 | 30.6 | 23.00 |
| all QMD | 20.8 | cherry | 2 | 6 | 6 | 6 | 0.73 | 6 | 0.06 |
| SrvyYr | 1891 | Douglas-fir | 10 | 6to54 | 26.6 | 23 | 3.66 | 31.1 | 8.19 |
| | | grand fir | 12 | 6to24 | 11.7 | 10 | 4.40 | 12.9 | 1.69 |
| | | hemlock | 207 | 4to48 | 16.9 | 16 | 75.82 | 19.2 | 64.63 |
| | | larch | 2 | 16to20 | 18 | 18 | 0.73 | 18.1 | 0.55 |
| | | bl maple | 1 | 8 | 8 | 8 | 0.37 | 8 | 0.05 |
| | | pine | 1 | 12 | 12 | 12 | 0.37 | 12 | 0.12 |
| | | spruce | 6 | 4to30 | 13.5 | 10 | 2.20 | 16.1 | 1.32 |

| Twp-Range | 19-1w | species | # WTs | Diam Range | av Diam | Median Diam | rel Freq% | QMD | rel DOM% |
|---|---|---|---|---|---|---|---|---|---|
| # allWTs | 203 | red alder | 4 | 8to12 | 9.5 | 9 | 1.97 | 9.6 | 0.48 |
| all avDiam | 16.5 | ash | 6 | 6to36 | 19.3 | 19 | 2.96 | 21.7 | 3.67 |
| all QMD | 19.5 | redcedar | 64 | 2to50 | 18 | 14 | 31.53 | 21.6 | 38.79 |
| SrvyYr | 1853 | crabapple | 1 | 10 | 10 | 10 | 0.49 | 10 | 0.13 |
| | | Douglas-fir | 117 | 6to40 | 15.9 | 12 | 57.64 | 18.6 | 52.58 |
| | | hemlock | 8 | 10to30 | 17.5 | 18 | 3.94 | 18.5 | 3.56 |
| | | bl maple | 1 | 18 | 18 | 18 | 0.49 | 18 | 0.42 |
| | | madrone | 1 | 6 | 6 | 6 | 0.49 | 6 | 0.05 |
| | | pine | 1 | 16 | 16 | 16 | 0.49 | 16 | 0.33 |

| Twp-Range | 19-2w | species | # WTs | Diam Range | av Diam | Median Diam | rel Freq% | QMD | rel DOM% |
|---|---|---|---|---|---|---|---|---|---|
| # allWTs | 165 | red alder | 10 | 4to25 | 10.6 | 10 | 6.06 | 11.9 | 2.64 |
| all avDiam | 15.7 | ash | 2 | 8to12 | 10 | 10 | 1.21 | 10.2 | 0.39 |
| all QMD | 18.0 | redcedar | 55 | 6to40 | 17.4 | 14 | 33.33 | 20.4 | 42.72 |
| SrvyYr | 1853 | cherry | 1 | 10 | 10 | 10 | 0.61 | 10 | 0.19 |
| | | Douglas-fir | 78 | 4to36 | 16.1 | 12 | 47.27 | 18 | 47.17 |
| | | hemlock | 9 | 6to24 | 10.9 | 10 | 5.45 | 12.1 | 2.46 |
| | | bl maple | 8 | 6to24 | 15.8 | 17 | 4.85 | 16.6 | 4.11 |
| | | madrone | 2 | 8 | 8 | 8 | 1.21 | 8 | 0.24 |

| Twp-Range | 19-3w | species | # WTs | Diam Range | av Diam | Median Diam | rel Freq% | QMD | rel DOM% |
|---|---|---|---|---|---|---|---|---|---|
| # allWTs | 239 | red alder | 17 | 2to24 | 9.6 | 8 | 7.11 | 11.3 | 2.51 |
| all avDiam | 16.2 | ash | 6 | 4to22 | 11.2 | 190 | 2.51 | 12.8 | 1.13 |
| all QMD | 19.0 | redcedar | 34 | 4to44 | 17.8 | 15 | 14.23 | 20.1 | 15.85 |
| SrvyYr | 1855 | dogwood | 1 | 10 | 10 | 10 | 0.42 | 10 | 0.12 |
| | | Douglas-fir | 117 | 5to85 | 19.5 | 18 | 48.95 | 22.4 | 67.75 |
| | | grand fir | 1 | 25 | 25 | 25 | 0.42 | 25 | 0.72 |
| | | hemlock | 46 | 4to25 | 10 | 8 | 19.25 | 10.9 | 6.31 |
| | | bl maple | 14 | 4to30 | 16.1 | 16 | 5.86 | 18 | 5.23 |
| | | madrone | 1 | 12 | 12 | 12 | 0.42 | 12 | 0.17 |
| | | oak | 1 | 18 | 18 | 18 | 0.42 | 18 | 0.37 |
| | | willow | 1 | 4 | 4 | 4 | 0.42 | 4 | 0.02 |

| Twp-Range | 19-4W | species | # WTs | Diam Range | av Diam | Median Diam | rel Freq% | QMD | rel DOM% |
|---|---|---|---|---|---|---|---|---|---|
| # allWTs | 286 | red alder | 29 | 4to24 | 12.2 | 12 | 10.14 | 13.4 | 2.73 |
| all avDiam | 19.9 | ash | 5 | 6to24 | 13.6 | 12 | 1.75 | 14.9 | 0.58 |
| all QMD | 25.8 | barberry | 6 | 4to6 | 4.5 | 4 | 2.10 | 4.6 | 0.07 |
| SrvyYr | 1864 | redcedar | 54 | 5to70 | 25 | 20 | 18.88 | 29.6 | 24.78 |
| | | cherry | 1 | 5 | 5 | 5 | 0.35 | 5 | 0.01 |
| | | crabapple | 1 | 12 | 12 | 12 | 0.35 | 12 | 0.08 |
| | | dogwood | 11 | 4to12 | 7.9 | 7 | 3.85 | 8.5 | 0.42 |
| | | Douglas-fir | 91 | 4to80 | 29.2 | 24 | 31.82 | 35.5 | 60.06 |
| | | hemlock | 54 | 4to40 | 13.8 | 12 | 18.88 | 16.1 | 7.33 |
| | | bl maple | 22 | 4to36 | 15.2 | 13 | 7.69 | 18.1 | 3.77 |
| | | pigeonwood | 1 | 5 | 5 | 5 | 0.35 | 5 | 0.01 |
| | | willow | 11 | 4to6 | 4.5 | 4 | 3.85 | 4.6 | 0.12 |

| Twp-Range | 19-5W | species | # WTs | Diam Range | av Diam | Median Diam | rel Freq% | QMD | rel DOM% |
|---|---|---|---|---|---|---|---|---|---|
| # allWTs | 265 | red alder | 5 | 4to18 | 10 | 10 | 1.89 | 11.3 | 0.41 |
| all avDiam | 19.5 | barberry | 1 | 8 | 8 | 8 | 0.38 | 8 | 0.04 |
| all QMD | 24.3 | redcedar | 34 | 3to80 | 36.7 | 36 | 12.83 | 40.8 | 36.19 |
| SrvyYr | 1874 | cherry | 3 | 6to18 | 12.7 | 14 | 1.13 | 13.6 | 0.35 |
| | | Douglas-fir | 45 | 8to60 | 28.3 | 24 | 16.98 | 31.7 | 28.92 |
| | | hemlock | 143 | 3to50 | 15.7 | 12 | 53.96 | 18.4 | 30.96 |
| | | bl maple | 8 | 12to30 | 290 | 20 | 3.02 | 20.8 | 2.21 |
| | | spruce | 3 | 6to12 | 9.3 | 10 | 1.13 | 9.7 | 0.18 |
| | | cascara | 1 | 12 | 12 | 12 | 0.38 | 12 | 0.09 |
| | | vine maple | 20 | 3to80 | 4.8 | 4 | 7.55 | 5.1 | 0.33 |
| | | willow | 2 | 8 | 8 | 8 | 0.75 | 8 | 0.08 |

| Twp-Range | 19-6w | species | # WTs | Diam Range | av Diam | Median Diam | rel Freq% | QMD | rel DOM% |
|---|---|---|---|---|---|---|---|---|---|
| # allWTs | 288 | red alder | 7 | 7to20 | 13.9 | 16 | 2.43 | 14.7 | 0.91 |
| all avDiam | 18.4 | redcedar | 18 | 5to96 | 27.6 | 24 | 6.25 | 34.3 | 12.80 |
| all QMD | 24.0 | cherry | 1 | 3 | 3 | 3 | 0.35 | 3 | 0.01 |
| SrvyYr | 1875 | crabapple | 3 | 3to12 | 6.7 | 5 | 1.04 | 8.8 | 0.14 |
| | | dogwood | 1 | 8 | 8 | 8 | 0.35 | 8 | 0.04 |
| | | Douglas-fir | 83 | 3to96 | 29.6 | 30 | 28.82 | 34.1 | 58.33 |
| | | grand fir | 2 | 12to18 | 15 | 15 | 0.69 | 15.3 | 0.28 |
| | | hemlock | 111 | 2to48 | 16.4 | 16 | 38.54 | 19.1 | 24.47 |
| | | hazel | 1 | 3 | 3 | 3 | 0.35 | 3 | 0.01 |
| | | bl maple | 6 | 3to48 | 20.5 | 19 | 2.08 | 25.7 | 2.40 |
| | | pine | 5 | 3to8 | 5.6 | 6 | 1.74 | 5.9 | 0.11 |
| | | serviceberr | 1 | 3 | 3 | 3 | 0.35 | 3 | 0.01 |
| | | cascara | 2 | 5to6 | 5.6 | 6 | 0.69 | 5.9 | 0.04 |
| | | vine maple | 45 | 3to8 | 4.2 | 4 | 15.63 | 4.4 | 0.53 |
| | | willow | 1 | 3 | 3 | 3 | 0.35 | 3 | 0.01 |
| | | yew | 1 | 12 | 12 | 12 | 0.35 | 12 | 0.09 |

| Twp-Range | 18-1e | species | # WTs | Diam Range | av Diam | Median Diam | rel Freq% | QMD | rel DOM% |
|---|---|---|---|---|---|---|---|---|---|
| # allWTs | 153 | red alder | 6 | 6to12 | 8 | 7 | 3.92 | 8.3 | 0.70 |
| all avDiam | 15.5 | ash | 4 | 6to14 | 10 | 10 | 2.61 | 10.5 | 0.75 |
| all QMD | 19.6 | redcedar | 4 | 5to36 | 18.3 | 16 | 2.61 | 21.6 | 3.16 |
| SrvyYr | 1873 | crabapple | 6 | 6to30 | 12 | 8 | 3.92 | 14.6 | 2.17 |
| | | dogwood | 2 | 7to10 | 8.5 | 8.5 | 1.31 | 8.6 | 0.25 |
| | | Douglas-fir | 90 | 4to96 | 17.8 | 14.5 | 58.82 | 22.5 | 77.22 |
| | | bl maple | 20 | 3to24 | 18.6 | 12 | 13.07 | 12.1 | 4.96 |
| | | oak | 15 | 4to36 | 18.5 | 20 | 9.80 | 20.6 | 10.79 |
| | | cascara | 1 | 6 | 6 | 6 | 0.65 | 6 | 0.06 |
| | | thorn | 2 | 6 | 6 | 6 | 1.31 | 6 | 0.12 |
| | | vine maple | 3 | 4to6 | 5.3 | 6 | 1.96 | 5.4 | 0.15 |

| Twp-Range | 18-2e | species | # WTs | Diam Range | av Diam | Median Diam | rel Freq% | QMD | rel DOM% |
|---|---|---|---|---|---|---|---|---|---|
| # allWTs | 133 | red alder | 1 | 8 | 8 | 8 | 0.75 | 8 | 0.08 |
| all avDiam | 21.4 | ash | 7 | 7to66 | 25 | 20 | 5.26 | 30.9 | 8.38 |
| all QMD | 24.5 | aspen | 1 | 5 | 5 | 5 | 0.75 | 5 | 0.03 |
| SrvyYr | 1871 | barberry | 1 | 5 | 5 | 5 | 0.75 | 5 | 0.03 |
| | | redcedar | 2 | 12to18 | 15 | 15 | 1.50 | 15.3 | 0.59 |
| | | dogwood | 1 | 16 | 16 | 16 | 0.75 | 16 | 0.32 |
| | | Douglas-fir | 62 | 4to72 | 18.9 | 14.5 | 46.62 | 22 | 37.61 |
| | | oak | 38 | 4to48 | 23.6 | 24 | 28.57 | 25.5 | 30.97 |
| | | pine | 17 | 9to54 | 29.8 | 30 | 12.78 | 31.8 | 21.55 |
| | | willow | 3 | 8to10 | 9.3 | 10 | 2.26 | 9.4 | 0.33 |

| Twp-Range | 18-3e | species | # WTs | Diam Range | av Diam | Median Diam | rel Freq% | QMD | rel DOM% |
|---|---|---|---|---|---|---|---|---|---|
| # allWTs | 126 | red alder | 3 | 8to20 | 15.3 | 18 | 2.38 | 16.2 | 1.04 |
| all avDiam | 22.0 | ash | 4 | 6to22 | 14 | 14 | 3.17 | 15.4 | 1.25 |
| all QMD | 24.5 | redcedar | 7 | 9to34 | 23 | 28 | 5.56 | 24.8 | 5.68 |
| SrvyYr | 1871 | crabapple | 1 | 7 | 7 | 7 | 0.79 | 7 | 0.06 |
| | | Douglas-fir | 82 | 3to75 | 22.9 | 20 | 65.08 | 25.7 | 71.42 |
| | | hemlock | 3 | 10to26 | 18 | 18 | 2.38 | 19.1 | 1.44 |
| | | oak | 19 | 12to48 | 23.7 | 24 | 15.08 | 25.4 | 16.16 |
| | | pine | 4 | 18to24 | 21 | 21 | 3.17 | 21.1 | 2.35 |
| | | willow | 3 | 7to14 | 11 | 12 | 2.38 | 11.4 | 0.51 |

| Twp-Range | 18-4e | species | # WTs | Diam Range | av Diam | Median Diam | rel Freq% | QMD | rel DOM% |
|---|---|---|---|---|---|---|---|---|---|
| # allWTs | 263 | red alder | 5 | 5to8 | 6.2 | 6 | 1.90 | 6.3 | 0.13 |
| all avDiam | 21.0 | ash | 1 | 8 | 8 | 8 | 0.38 | 8 | 0.04 |
| all QMD | 24.5 | redcedar | 56 | 6to60 | 21.7 | 16 | 21.29 | 25.9 | 23.80 |
| SrvyYr | 1873 | cherry | 7 | 6to48 | 19.1 | 16 | 2.66 | 22.9 | 2.33 |
| | | Douglas-fir | 166 | 6to60 | 22.6 | 20 | 63.12 | 25.7 | 69.47 |
| | | hemlock | 20 | 4to24 | 15.2 | 17 | 7.60 | 16.5 | 3.45 |
| | | bl maple | 4 | 6to24 | 15 | 15 | 1.52 | 16.4 | 0.68 |
| | | willow | 4 | 3to5 | 4 | 4 | 1.52 | 4.1 | 0.04 |

| Twp-Range | 18-5e | species | # WTs | Diam Range | av Diam | Median Diam | rel Freq% | QMD | rel DOM% |
|---|---|---|---|---|---|---|---|---|---|
| # allWTs | 287 | red alder | 11 | 8to18 | 12.7 | 12 | 3.83 | 13 | 1.07 |
| all avDiam | 20.2 | ash | 2 | 12 | 12 | 12 | 0.70 | 12 | 0.17 |
| all QMD | 24.6 | redcedar | 62 | 3to48 | 18.9 | 16 | 21.60 | 21.6 | 16.63 |
| SrvyYr | 1874 | cherry | 4 | 5to10 | 6.5 | 5.5 | 1.39 | 6.8 | 0.11 |
| | | cottonwood | 7 | 6to18 | 12 | 12 | 2.44 | 12.8 | 0.66 |
| | | Douglas-fir | 105 | 4to60 | 28.6 | 24 | 36.59 | 33.2 | 66.52 |
| | | hemlock | 81 | 5to48 | 14.7 | 12 | 28.22 | 16.8 | 13.14 |
| | | bl maple | 10 | 5to20 | 11.5 | 12 | 3.48 | 12.2 | 0.86 |
| | | spruce | 1 | 36 | 36 | 36 | 0.35 | 36 | 0.74 |
| | | vine maple | 1 | 6 | 6 | 6 | 0.35 | 6 | 0.02 |
| | | willow | 3 | 5to8 | 6.3 | 6 | 1.05 | 6.5 | 0.07 |

| Twp-Range | 18-6e | species | # WTs | Diam Range | av Diam | Median Diam | rel Freq% | QMD | rel DOM% |
|---|---|---|---|---|---|---|---|---|---|
| # allWTs | 289 | red alder | 3 | 4to10 | 7.3 | 7 | 1.04 | 7.7 | 0.20 |
| all avDiam | 13.8 | redcedar | 25 | 2to72 | 18.6 | 12 | 8.65 | 25.1 | 17.80 |
| all QMD | 17.5 | Douglas-fir | 8 | 6to36 | 19.8 | 19 | 2.77 | 21.2 | 4.06 |
| SrvyYr | 1875 | grand fir | 18 | 4to50 | 13.4 | 5.5 | 6.23 | 18.6 | 7.04 |
| | | hemlock | 229 | 4to50 | 13.2 | 10 | 79.24 | 16.4 | 69.60 |
| | | bl maple | 1 | 5 | 5 | 5 | 0.35 | 5 | 0.03 |
| | | spruce | 2 | 20to24 | 22 | 22 | 0.69 | 22.1 | 1.10 |
| | | yew | 3 | 4to20 | 12 | 12 | 1.04 | 12.1 | 0.50 |

| Twp-Range | 18-1w | species | # WTs | Diam Range | av Diam | Median Diam | rel Freq% | QMD | rel DOM% |
|---|---|---|---|---|---|---|---|---|---|
| # allWTs | 230 | red alder | 3 | 8to12 | 10 | 10 | 1.30 | 10.1 | 0.31 |
| all avDiam | 17.7 | ash | 10 | 6to30 | 14 | 10 | 4.35 | 16.2 | 2.63 |
| all QMD | 20.8 | redcedar | 58 | 4to60 | 19.2 | 16 | 25.22 | 23 | 30.75 |
| SrvyYr | 1853 | crabapple | 1 | 4 | 4 | 4 | 0.43 | 4 | 0.02 |
| | | Douglas-fir | 121 | 4to60 | 17.1 | 12 | 52.61 | 20 | 48.51 |
| | | hemlock | 5 | 12to24 | 18.8 | 20 | 2.17 | 19.5 | 1.91 |
| | | bl maple | 3 | 8to20 | 12.7 | 10 | 1.30 | 13.7 | 0.56 |
| | | oak | 18 | 6to40 | 25.9 | 24 | 7.83 | 27.5 | 13.64 |
| | | pine | 8 | 6to20 | 11.3 | 9 | 3.48 | 12.5 | 1.25 |
| | | spruce | 1 | 12 | 12 | 12 | 0.43 | 12 | 0.14 |
| | | willow | 2 | 4to8 | 6 | 6 | 0.87 | 6.3 | 0.08 |

| Twp-Range | 18-2w | species | # WTs | Diam Range | av Diam | Median Diam | rel Freq% | QMD | rel DOM% |
|---|---|---|---|---|---|---|---|---|---|
| # allWTs | 250 | red alder | 18 | 4to12 | 8.1 | 8 | 7.20 | 8.5 | 1.27 |
| all avDiam | 17.1 | ash | 2 | 12to14 | 13 | 13 | 0.80 | 13 | 0.33 |
| all QMD | 20.2 | redcedar | 70 | 4to60 | 20.3 | 16 | 28.00 | 24.1 | 39.82 |
| SrvyYr | 1853 | dogwood | 1 | 10 | 10 | 10 | 0.40 | 10 | 0.10 |
| | | Douglas-fir | 110 | 4to50 | 18.2 | 14 | 44.00 | 21.1 | 47.96 |
| | | hemlock | 24 | 6to24 | 12.8 | 12 | 9.60 | 13.6 | 4.35 |
| | | bl maple | 18 | 12to36 | 17.1 | 16 | 7.20 | 18 | 5.71 |
| | | oak | 1 | 16 | 16 | 16 | 0.40 | 16 | 0.25 |
| | | pine | 1 | 12 | 12 | 12 | 0.40 | 12 | 0.14 |
| | | spruce | 3 | 6to12 | 8.7 | 8 | 1.20 | 9 | 0.24 |
| | | cascara | 1 | 4 | 4 | 4 | 0.40 | 4 | 0.02 |
| | | vine maple | 1 | 4 | 4 | 4 | 0.40 | 4 | 0.02 |

| Twp-Range | 18-3w | species | # WTs | Diam Range | av Diam | Median Diam | rel Freq% | QMD | rel DOM% |
|---|---|---|---|---|---|---|---|---|---|
| # allWTs | 276 | red alder | 8 | 4to16 | 9.1 | 7.5 | 2.90 | 10 | 0.71 |
| all avDiam | 16.6 | ash | 1 | 3 | 3 | 3 | 0.36 | 3 | 0.01 |
| all QMD | 20.2 | redcedar | 57 | 6to70 | 20.5 | 16 | 20.65 | 25.3 | 32.39 |
| SrvyYr | 1855 | Douglas-fir | 132 | 3to60 | 18.7 | 16 | 47.83 | 21.5 | 54.17 |
| | &75 | grand fir | 1 | 30 | 30 | 30 | 0.36 | 30 | 0.80 |
| | | hemlock | 60 | 3to40 | 11.6 | 10 | 21.74 | 13.8 | 10.14 |
| | | bl maple | 15 | 3to20 | 8.7 | 7 | 5.43 | 10.1 | 1.36 |
| | | oak | 1 | 18 | 18 | 18 | 0.36 | 18 | 0.29 |
| | | vine maple | 1 | 3 | 3 | 3 | 0.36 | 3 | 0.01 |

| Twp-Range | 18-4w | species | # WTs | Diam Range | av Diam | Median Diam | rel Freq% | QMD | rel DOM% |
|---|---|---|---|---|---|---|---|---|---|
| # allWTs | 288 | red alder | 4 | 6to18 | 13.5 | 15 | 1.39 | 14.2 | 0.71 |
| all avDiam | 15.5 | redcedar | 26 | 6to60 | 24.8 | 19 | 9.03 | 29.5 | 19.98 |
| all QMD | 19.8 | Douglas-fir | 47 | 6to84 | 23.9 | 18 | 16.32 | 28.7 | 34.19 |
| SrvyYr | 1872 | hemlock | 165 | 4to40 | 13.6 | 10 | 57.29 | 16 | 37.31 |
| | | bl maple | 12 | 5to20 | 13.9 | 14 | 4.17 | 14.6 | 2.26 |
| | | spruce | 1 | 72 | 72 | 72 | 0.35 | 72 | 4.58 |
| | | cascara | 2 | 5to8 | 6.5 | 6.5 | 0.69 | 6.7 | 0.08 |
| | | vine maple | 31 | 4to8 | 4.7 | 4 | 10.76 | 4.8 | 0.63 |

| Twp-Range | 17-1e | species | # WTs | Diam Range | av Diam | Median Diam | rel Freq% | QMD | rel DOM% |
|---|---|---|---|---|---|---|---|---|---|
| # allWTs | 148 | red alder | 2 | 12to20 | 16 | 16 | 1.35 | 16.5 | 1.30 |
| all avDiam | 14.4 | ash | 3 | 3to12 | 9 | 12 | 2.03 | 9.9 | 0.70 |
| all QMD | 16.8 | redcedar | 25 | 8to40 | 15.7 | 12 | 16.89 | 18.2 | 19.84 |
| SrvyYr | 1853 | crabapple | 3 | 4to6 | 5.3 | 6 | 2.03 | 5.4 | 0.21 |
| | | Douglas-fir | 89 | 5to48 | 14.1 | 12 | 60.14 | 16.2 | 55.97 |
| | | hemlock | 5 | 4to18 | 10 | 10 | 3.38 | 11.1 | 1.48 |
| | | bl maple | 2 | 10to18 | 14 | 14 | 1.35 | 14.6 | 1.02 |
| | | oak | 15 | 6to36 | 17.9 | 14 | 10.14 | 20.2 | 14.67 |
| | | spruce | 1 | 40 | 40 | 40 | 0.68 | 40 | 3.83 |
| | | thorn | 1 | 15 | 15 | 15 | 0.68 | 15 | 0.54 |
| | | vine maple | 1 | 4 | 4 | 4 | 0.68 | 4 | 0.04 |
| | | willow | 1 | 4 | 4 | 4 | 0.68 | 4 | 0.04 |

| Twp-Range | 17-2e | species | # WTs | Diam Range | av Diam | Median Diam | rel Freq% | QMD | rel DOM% |
|---|---|---|---|---|---|---|---|---|---|
| # allWTs | 215 | red alder | 5 | 10to12 | 11.2 | 12 | 2.33 | 11.2 | 0.52 |
| all avDiam | 21.3 | ash | 5 | 7to24 | 12.6 | 10 | 2.33 | 14 | 0.81 |
| all QMD | 23.8 | redcedar | 32 | 8to72 | 20.3 | 20 | 14.88 | 23.2 | 14.18 |
| SrvyYr | 1870 | cottonwood | 4 | 4to15 | 11.3 | 13 | 1.86 | 12.1 | 0.48 |
| | | dogwood | 2 | 11 | 11 | 11 | 0.93 | 11 | 0.20 |
| | | Douglas-fir | 151 | 5to54 | 22.4 | 20 | 70.23 | 24.4 | 74.04 |
| | | hemlock | 2 | 12 | 12 | 12 | 0.93 | 12 | 0.24 |
| | | bl maple | 2 | 12 | 12 | 12 | 0.93 | 12 | 0.24 |
| | | oak | 10 | 10to48 | 27.4 | 25 | 4.65 | 29.8 | 7.31 |
| | | pine | 1 | 48 | 48 | 48 | 0.47 | 48 | 1.90 |
| | | yew | 1 | 12 | 12 | 12 | 0.47 | 12 | 0.12 |

| Twp-Range | 17-3e | species | # WTs | Diam Range | av Diam | Median Diam | rel Freq% | QMD | rel DOM% |
|---|---|---|---|---|---|---|---|---|---|
| # allWTs | 283 | red alder | 11 | 5to10 | 7.2 | 8 | 3.89 | 7.4 | 0.62 |
| all avDiam | 15.6 | ash | 10 | 7to20 | 10.2 | 10 | 3.53 | 10.9 | 1.22 |
| all QMD | 18.6 | redcedar | 52 | 7to48 | 17.3 | 14 | 18.37 | 19.6 | 20.44 |
| SrvyYr | 1873 | crabapple | 1 | 16 | 16 | 16 | 0.35 | 16 | 0.26 |
| | | cottonwood | 10 | 4to24 | 9.5 | 8 | 3.53 | 11 | 1.24 |
| | | Douglas-fir | 172 | 5to72 | 16.9 | 14 | 60.78 | 20.1 | 71.09 |
| | | hemlock | 13 | 6to18 | 12.2 | 12 | 4.59 | 12.8 | 2.18 |
| | | bl maple | 6 | 5to18 | 9.7 | 7 | 2.12 | 11 | 0.74 |
| | | oak | 3 | 24to30 | 28 | 30 | 1.06 | 28.1 | 2.42 |
| | | willow | 5 | 4to8 | 5.2 | 5 | 1.77 | 5.4 | 0.15 |

| Twp-Range | 17-4e | species | # Wit | Diam iamRang | av meter | Median Median | rel FREQ% | QMD | rel relDOM% |
|---|---|---|---|---|---|---|---|---|---|
| # allWTs | 278 | red alder | 18 | 4to12 | 6.8 | 6 | 6.47 | 7.3 | 0.64 |
| all avDiam | 19.5 | ash | 8 | 8to15 | 10.6 | 10 | 2.88 | 8.6 | 0.39 |
| all QMD | 23.2 | redcedar | 79 | 6to48 | 19.5 | 16 | 28.42 | 22.1 | 25.71 |
| SrvyYr | 1873 | cherry | 1 | 6 | 6 | 6 | 0.36 | 6 | 0.02 |
| | | Douglas-fir | 127 | 4to60 | 24.8 | 24 | 45.68 | 28.3 | 67.78 |
| | | hemlock | 25 | 4to24 | 12.3 | 10 | 8.99 | 13.5 | 3.04 |
| | | bl maple | 12 | 4to24 | 11.1 | 12 | 4.32 | 12.3 | 1.21 |
| | | oak | 1 | 30 | 30 | 30 | 0.36 | 30 | 0.60 |
| | | spruce | 2 | 5to16 | 10.5 | 10.5 | 0.72 | 11.9 | 0.19 |
| | | willow | 5 | 3to5 | 3.8 | 4 | 1.80 | 3.9 | 0.05 |

| Twp-Range | 17-5e | species | # WTs | Diam Range | av Diam | Median Diam | rel Freq% | QMD | rel DOM% |
|---|---|---|---|---|---|---|---|---|---|
| # allWTs | 280 | red alder | 4 | 4to18 | 9.8 | 8.5 | 1.43 | 11.1 | 0.17 |
| all avDiam | 27.2 | redcedar | 48 | 7to120 | 32.5 | 30 | 17.14 | 38.1 | 24.12 |
| all QMD | 32.1 | Douglas-fir | 70 | 5to70 | 37.2 | 36 | 25.00 | 40.9 | 40.54 |
| SrvyYr | 1890 | grand fir | 10 | 6to56 | 24.4 | 24 | 3.57 | 27.6 | 2.64 |
| | | hemlock | 143 | 4to55 | 21.3 | 17 | 51.07 | 25 | 30.94 |
| | | bl maple | 4 | 6to24 | 12.5 | 10 | 1.43 | 14.2 | 0.28 |
| | | spruce | 1 | 60 | 60 | 60 | 0.36 | 60 | 1.25 |

| Twp-Range | 17-1w | species | # WTs | Diam Range | av Diam | Median Diam | rel Freq% | QMD | rel DOM% |
|---|---|---|---|---|---|---|---|---|---|
| # allWTs | 224 | red alder | 13 | 4to20 | 9.3 | 10 | 5.80 | 10.1 | 1.45 |
| all avDiam | 16.8 | ash | 3 | 6to24 | 13.3 | 10 | 1.34 | 15.4 | 0.78 |
| all QMD | 20.2 | redcedar | 43 | 5to72 | 20 | 16 | 19.20 | 24 | 27.03 |
| SrvyYr | 1853 | Douglas-fir | 128 | 3to72 | 17.1 | 14 | 57.14 | 20.3 | 57.55 |
| | | hemlock | 6 | 6to30 | 11.7 | 8 | 2.68 | 14.3 | 1.34 |
| | | bl maple | 13 | 4to24 | 11.3 | 8 | 5.80 | 13.1 | 2.43 |
| | | oak | 15 | 10to40 | 21.5 | 18 | 6.70 | 23.8 | 9.27 |
| | | spruce | 1 | 12 | 12 | 12 | 0.45 | 12 | 0.16 |
| | | vine maple | 2 | 5 | 5 | 5 | 0.89 | 5 | 0.05 |

| Twp-Range | 17-2w | species | # WTs | Diam Range | av Diam | Median Diam | rel Freq% | QMD | rel DOM% |
|---|---|---|---|---|---|---|---|---|---|
| # allWTs | 248 | red alder | 32 | 3to20 | 9.1 | 8 | 12.90 | 9.8 | 2.83 |
| all avDiam | 16.9 | ash | 9 | 8to14 | 10.9 | 12 | 3.63 | 11.1 | 1.02 |
| all QMD | 20.9 | redcedar | 57 | 4to50 | 23.1 | 18 | 22.98 | 27 | 38.21 |
| SrvyYr | 1855 | dogwood | 1 | 6 | 6 | 6 | 0.40 | 6 | 0.03 |
| | | Douglas-fir | 110 | 3to60 | 19.4 | 16 | 44.35 | 23 | 53.50 |
| | | hemlock | 11 | 5to30 | 12.6 | 8 | 4.44 | 15.9 | 2.56 |
| | | bl maple | 3 | 8to14 | 11.3 | 12 | 1.21 | 11.6 | 0.37 |
| | | pine | 13 | 4to12 | 7.7 | 8 | 5.24 | 8 | 0.76 |
| | | spruce | 1 | 8 | 8 | 8 | 0.40 | 8 | 0.06 |
| | | thorn | 2 | 2to10 | 6 | 6 | 0.81 | 7.2 | 0.10 |
| | | vine maple | 1 | 6 | 6 | 6 | 0.40 | 6 | 0.03 |
| | | willow | 8 | 4to20 | 6.8 | 5 | 3.23 | 8.4 | 0.52 |

| Twp-Range | 17-3w | species | # WTs | Diam Range | av Diam | Median Diam | rel Freq% | QMD | rel DOM% |
|---|---|---|---|---|---|---|---|---|---|
| # allWTs | 259 | red alder | 28 | 5to30 | 12.4 | 10.5 | 10.81 | 14.1 | 4.38 |
| all avDiam | 17.9 | ash | 1 | 6 | 6 | 6 | 0.39 | 6 | 0.03 |
| all QMD | 22.2 | redcedar | 41 | 4to44 | 18.8 | 16 | 15.83 | 21.1 | 14.35 |
| SrvyYr | 1855 | dogwood | 11 | 5to15 | 9.5 | 9 | 4.25 | 10 | 0.86 |
| | | Douglas-fir | 104 | 3to84 | 23 | 17.5 | 40.15 | 28.3 | 65.47 |
| | | hemlock | 54 | 5to36 | 14.1 | 11.5 | 20.85 | 15.9 | 10.73 |
| | | bl maple | 3 | 15to34 | 26 | 27 | 1.16 | 26.5 | 1.66 |
| | | pine | 4 | 6to40 | 25.3 | 20 | 1.54 | 24.9 | 1.95 |
| | | spruce | 1 | 8 | 8 | 8 | 0.39 | 8 | 0.05 |
| | | vine maple | 9 | 4to6 | 4.7 | 5 | 3.47 | 4.7 | 0.16 |
| | | willow | 1 | 5 | 5 | 5 | 0.39 | 5 | 0.02 |
| | | yew | 2 | 10to14 | 12 | 12 | 0.77 | 12.2 | 0.23 |

| Twp-Range | 16-1e | species | # WTs | Diam Range | av Diam | Median Diam | rel Freq% | QMD | rel DOM% |
|---|---|---|---|---|---|---|---|---|---|
| # allWTs | 217 | red alder | 7 | 7to20 | 12.1 | 12 | 3.23 | 12.9 | 0.80 |
| all avDiam | 23.1 | ash | 8 | 10to20 | 15.1 | 15.5 | 3.69 | 15.5 | 1.32 |
| all QMD | 25.9 | redcedar | 47 | 7to48 | 21.9 | 20 | 21.66 | 24.8 | 19.85 |
| SrvyYr | 1855 | Douglas-fir | 120 | 8to68 | 26.8 | 24 | 55.30 | 29.2 | 70.24 |
| | | grand fir | 2 | 14to36 | 25 | 25 | 0.92 | 27.3 | 1.02 |
| | | hemlock | 17 | 6to34 | 16.2 | 15 | 7.83 | 17.7 | 3.66 |
| | | bl maple | 10 | 6to24 | 13.1 | 12 | 4.61 | 14.2 | 1.38 |
| | | oak | 4 | 10to30 | 22 | 24 | 1.84 | 23.6 | 1.53 |
| | | pine | 1 | 10 | 10 | 10 | 0.46 | 10 | 0.07 |
| | | willow | 1 | 4 | 4 | 4 | 0.46 | 4 | 0.01 |

| Twp-Range | 16-2e | species | # WTs | Diam Range | av Diam | Median Diam | rel Freq% | QMD | rel DOM% |
|---|---|---|---|---|---|---|---|---|---|
| # allWTs | 277 | red alder | 20 | 5to12 | 7.5 | 7.5 | 7.22 | 7.7 | 1.68 |
| all avDiam | 13.3 | ash | 10 | 5to14 | 7.4 | 5 | 3.61 | 8.1 | 0.93 |
| all QMD | 15.9 | redcedar | 33 | 4to48 | 15.9 | 18 | 11.91 | 18.4 | 15.86 |
| SrvyYr | 1873 | cottonwood | 2 | 10to12 | 11 | 11 | 0.72 | 11 | 0.34 |
| | | dogwood | 6 | 3to10 | 5.8 | 5 | 2.17 | 6.3 | 0.34 |
| | | Douglas-fir | 164 | 3to60 | 15.5 | 14 | 59.21 | 18 | 75.42 |
| | | hemlock | 6 | 5to12 | 8 | 8 | 2.17 | 8.4 | 0.60 |
| | | bl maple | 8 | 3to14 | 6.8 | 6 | 2.89 | 7.4 | 0.62 |
| | | madrone | 3 | 8to24 | 14.7 | 12 | 1.08 | 16.2 | 1.12 |
| | | oak | 6 | 3to28 | 12.3 | 7.5 | 2.17 | 15.9 | 2.15 |
| | | pine | 10 | 4to15 | 7.9 | 7 | 3.61 | 8.6 | 1.05 |
| | | cascara | 2 | 2to3 | 2.5 | 2.5 | 0.72 | 2.5 | 0.02 |
| | | vine maple | 1 | 6 | 6 | 6 | 0.36 | 6 | 0.05 |
| | | willow | 6 | 3to5 | 4.2 | 4 | 2.17 | 4.6 | 0.18 |

| Twp-Range | 16-3e | species | # WTs | Diam Range | av Diam | Median Diam | rel Freq% | QMD | rel DOM% |
|---|---|---|---|---|---|---|---|---|---|
| ttl trees | 277 | red alder | 16 | 4to16 | 8.1 | 8 | 5.78 | 8.6 | 1.55 |
| av diam | 13.8 | ash | 10 | 4to12 | 8.3 | 8 | 3.61 | 8.7 | 0.99 |
| ttlQMD | 16.6 | redcedar | 49 | 6to72 | 17.3 | 12 | 17.69 | 22 | 31.09 |
| SrvyYr | 1873 | crabapple | 2 | 5to8 | 6.6 | 6.6 | 0.72 | 6.7 | 0.12 |
| | | cottonwood | 3 | 6to30 | 17.3 | 16 | 1.08 | 19.9 | 1.56 |
| | | dogwood | 1 | 8 | 8 | 8 | 0.36 | 8 | 0.08 |
| | | Douglas-fir | 170 | 3to48 | 14.5 | 12 | 61.37 | 16.6 | 61.41 |
| | | hemlock | 8 | 8to18 | 10.9 | 10 | 2.89 | 11.3 | 1.34 |
| | | bl maple | 15 | 4to20 | 8.1 | 7 | 5.42 | 8.9 | 1.56 |
| | | madrone | 1 | 18 | 18 | 18 | 0.36 | 18 | 0.42 |
| | | willow | 2 | 5 | 5 | 5 | 0.72 | 5 | 0.07 |

| Twp-Range | 16-4e | species | # WTs | Diam Range | av Diam | Median Diam | rel Freq% | QMD | rel DOM% |
|---|---|---|---|---|---|---|---|---|---|
| # allWTs | 269 | red alder | 15 | 3to16 | 6.7 | 5 | 5.58 | 7.8 | 0.81 |
| all avDiam | 16.6 | ash | 1 | 15 | 15 | 15 | 0.37 | 15 | 0.20 |
| all QMD | 20.5 | redcedar | 67 | 3to40 | 17.1 | 12 | 24.91 | 19.5 | 22.58 |
| SrvyYr | 1893 | cherry | 1 | 4 | 4 | 4 | 0.37 | 4 | 0.01 |
| | | cottonwood | 1 | 6 | 6 | 6 | 0.37 | 6 | 0.03 |
| | | dogwood | 1 | 5 | 5 | 5 | 0.37 | 5 | 0.02 |
| | | Douglas-fir | 122 | 4to60 | 21 | 15.5 | 45.35 | 25.2 | 68.66 |
| | | grand fir | 1 | 30 | 30 | 30 | 0.37 | 30 | 0.80 |
| | | hemlock | 38 | 5to26 | 10.6 | 9.5 | 14.13 | 12 | 4.85 |
| | | bl maple | 13 | 5to24 | 10 | 10 | 4.83 | 11.2 | 1.45 |
| | | madrone | 3 | 4to12 | 7.3 | 6 | 1.12 | 8.1 | 0.17 |
| | | vine maple | 2 | 4to6 | 5 | 5 | 0.74 | 5.1 | 0.05 |
| | | willow | 4 | 3to8 | 4.5 | 3.5 | 1.49 | 4.9 | 0.09 |

| Twp-Range | 16-1w | species | # WTs | Diam Range | av Diam | Median Diam | rel Freq% | QMD | rel DOM% |
|---|---|---|---|---|---|---|---|---|---|
| # allWTs | 267 | red alder | 13 | 6to18 | 10.2 | 10 | 4.87 | 10.7 | 1.22 |
| all avDiam | 18.8 | ash | 6 | 4to20 | 10.5 | 11 | 2.25 | 11.8 | 0.68 |
| all QMD | 21.4 | redcedar | 84 | 5to105 | 19.3 | 18 | 31.46 | 21 | 30.35 |
| SrvyYr | | cherry | 1 | 7 | 7 | 7 | 0.37 | 7 | 0.04 |
| | | Douglas-fir | 121 | 5to60 | 22.3 | 20 | 45.32 | 24.9 | 61.47 |
| | | hemlock | 15 | 6to16 | 10.2 | 9 | 5.62 | 10.6 | 1.38 |
| | | bl maple | 11 | 6to24 | 12.7 | 10 | 4.12 | 14 | 1.77 |
| | | oak | 11 | 10to24 | 17.9 | 18 | 4.12 | 18.7 | 3.15 |
| | | vine maple | 1 | 4 | 4 | 4 | 0.37 | 4 | 0.01 |
| | | willow | 4 | 4to7 | 5.5 | 5.5 | 1.50 | 5.7 | 0.11 |

| Twp-Range | 16-2w | species | # WTs | Diam Range | av Diam | Median Diam | rel Freq% | QMD | rel DOM% |
|---|---|---|---|---|---|---|---|---|---|
| # allWTs | 220 | red alder | 30 | 4to24 | 10.3 | 10 | 13.64 | 11.1 | 3.54 |
| all avDiam | 18.0 | ash | 8 | 4to14 | 9.4 | 9.5 | 3.64 | 10.1 | 0.78 |
| all QMD | 21.8 | redcedar | 48 | 5to44 | 21.1 | 20 | 21.82 | 23.6 | 25.58 |
| SrvyYr | 1855 | crabapple | 1 | 5 | 5 | 5 | 0.45 | 5 | 0.02 |
| | | dogwood | 2 | 6to9 | 7.5 | 7.5 | 0.91 | 7.6 | 0.11 |
| | | Douglas-fir | 99 | 5to60 | 21.4 | 16 | 45.00 | 25.6 | 62.08 |
| | | hemlock | 8 | 6to20 | 9.5 | 8 | 3.64 | 10.5 | 0.84 |
| | | bl maple | 8 | 3to30 | 11.6 | 12 | 3.64 | 14.2 | 1.54 |
| | | oak | 12 | 10to30 | 20.4 | 22 | 5.45 | 21.7 | 5.41 |
| | | thorn | 1 | 4 | 4 | 4 | 0.45 | 4 | 0.02 |
| | | willow | 3 | 3to4 | 3.7 | 4 | 1.36 | 3.7 | 0.04 |

| Twp-Range | 16-3w | species | # WTs | Diam Range | av Diam | Median Diam | rel Freq% | QMD | rel DOM% |
|---|---|---|---|---|---|---|---|---|---|
| # allWTs | 207 | red alder | 3 | 6to20 | 14.7 | 18 | 1.45 | 15.9 | 0.57 |
| all avDiam | 21.3 | ash | 16 | 5to20 | 10.3 | 9 | 7.73 | 11.6 | 1.62 |
| all QMD | 25.3 | redcedar | 50 | 6to96 | 23.6 | 22 | 24.15 | 27.5 | 28.52 |
| SrvyYr | 1871 | crabapple | 2 | 4to6 | 5 | 5 | 0.97 | 5.1 | 0.04 |
| | | dogwood | 8 | 6to20 | 11.6 | 10 | 3.86 | 12.5 | 0.94 |
| | | Douglas-fir | 92 | 5to72 | 25.2 | 24 | 44.44 | 29 | 58.36 |
| | | hemlock | 14 | 6to30 | 15.7 | 16 | 6.76 | 16.7 | 2.95 |
| | | bl maple | 6 | 10to30 | 22.7 | 24 | 2.90 | 23.7 | 2.54 |
| | | oak | 10 | 10to40 | 19.9 | 14.5 | 4.83 | 22.3 | 3.75 |
| | | pine | 1 | 10 | 10 | 10 | 0.48 | 10 | 0.08 |
| | | willow | 2 | 4 | 4 | 4 | 0.97 | 4 | 0.02 |
| | | yellowwooc | 2 | 4to6 | 5 | 5 | 0.97 | 5.1 | 0.04 |
| | | yew | 1 | 23 | 23 | 23 | 0.48 | 23 | 0.40 |

| Twp-Range | North Whidbey | species | # WTs | Diam Range | av Diam | Median Diam | rel Freq% | QMD | rel DOM% |
|---|---|---|---|---|---|---|---|---|---|
| # allWTs | 662 | alder | 75 | 4to16 | 5.56 | 5 | 11.33 | 5.99 | 0.88 |
| all avDiam | 17.4 | redcedar | 16 | 8to40 | 21.81 | 20 | 2.42 | 24.4 | 3.11 |
| all QMD | 21.5 | cherry | 1 | 4 | 4 | 4 | 0.15 | 4 | 0.01 |
| SrvyYr | 1855 | cottonw | 3 | 5to6 | 5.3 | 5 | 0.45 | 5.4 | 0.03 |
| | &59 | Douglas-fir | 422 | 4to16 | 22.09 | 20 | 63.75 | 25.5 | 89.10 |
| | | hemlock | 46 | 4to32 | 12.17 | 9 | 6.95 | 13.8 | 2.86 |
| | | bl maple | 2 | 12to14 | 13 | 13 | 0.30 | 13 | 0.11 |
| | | oak | 3 | 14to40 | 23.3 | 16 | 0.45 | 26.2 | 0.67 |
| | | pine | 1 | 8 | 8 | 8 | 0.15 | 8 | 0.02 |
| | | spruce | 23 | 4to40 | 11.91 | 9 | 3.47 | 14.8 | 1.64 |
| | | willow | 69 | 4to24 | 6.9 | 5 | 10.42 | 8.03 | 1.45 |
| | | yew | 1 | 10 | 10 | 10 | 0.15 | 10 | 0.03 |

[illegible]

| Twp-Range | S Whidbey &Camano | species | # WTs | Diam Range | av Diam | Median Diam | rel Freq% | QMD | rel DOM% |
|---|---|---|---|---|---|---|---|---|---|
| # allWTs | 876 | alder | 51 | 3to18 | 6.6 | 5 | 5.82 | 7.45 | 0.69 |
| all avDiam | 17.3 | redcedar | 233 | 4to72 | 21.4 | 18 | 26.60 | 25.4 | 36.89 |
| all QMD | 21.6 | cherry | 1 | 4 | 4 | 4 | 0.11 | 4 | 0.00 |
| SrvyYr | 1855 | crabapple | 2 | 4to6 | 5 | 5 | 0.23 | 5.1 | 0.01 |
| | &59 | cottonw | 1 | 15 | 15 | 15 | 0.11 | 15 | 0.06 |
| | | dogw | 2 | 6to8 | 7 | 7 | 0.23 | 7.07 | 0.02 |
| | | Douglas-fir | 271 | 2to60 | 23.14 | 20 | 30.94 | 27.2 | 48.99 |
| | | grand fir | 6 | 8to30 | 16.83 | 16.5 | 0.68 | 18.3 | 0.49 |
| | | hemlock | 261 | 2to42 | 11.41 | 10 | 29.79 | 13.6 | 11.72 |
| | | bl maple | 13 | 8to16 | 13.2 | 14.5 | 1.48 | 13.4 | 0.57 |
| | | pine | 10 | 3to18 | 7.1 | 6 | 1.14 | 8.11 | 0.16 |
| | | spruce | 7 | 6to30 | 11.7 | 8 | 0.80 | 14.1 | 0.34 |
| | | willow | 14 | 3to20 | 5.64 | 3.5 | 1.60 | 7.23 | 0.18 |
| | | yew | 4 | 6to8 | 6.75 | 6.5 | 0.46 | 6.8 | 0.05 |

| Twp-Range | Bainbrid | species | # WTs | Diam Range | av Diam | Median Diam | rel Freq% | QMD | rel DOM% |
|---|---|---|---|---|---|---|---|---|---|
| # allWTs | 216 | red alder | 10 | 5to12 | 8.6 | 7.5 | 4.63 | 9.59 | 0.64 |
| all avDiam | 21.8 | redcedar | 85 | 3to60 | 21.59 | 20 | 39.35 | 24 | 34.10 |
| all QMD | 25.7 | Douglas-fir | 86 | 2to70 | 26.78 | 25.5 | 39.81 | 31.4 | 59.21 |
| SrvyYr | 1857 | hemlock | 24 | 5to28 | 13 | 11 | 11.11 | 14.3 | 3.41 |
| | | madrone | 2 | 4to5 | 4.5 | 4.5 | 0.93 | 4.5 | 0.03 |
| | | bl maple | 5 | 12to30 | 18 | 18 | 2.31 | 19.2 | 1.28 |
| | | grand fir | 1 | 40 | 40 | 40 | 0.46 | 40 | 1.12 |
| | | willow | 2 | 5 | 5 | 5 | 0.93 | 5 | 0.03 |
| | | yew | 1 | 14 | 14 | 14 | 0.46 | 14 | 0.14 |

| Twp-Range | Vashon | species | # WTs | Diam Range | av Diam | Median Diam | rel Freq% | QMD | rel DOM% |
|---|---|---|---|---|---|---|---|---|---|
| # allWTs | 263 | red alder | 19 | 4to18 | 9.11 | 8 | 7.22 | 10.3 | 1.22 |
| all avDiam | 20.5 | redcedar | 60 | 3to60 | 25.33 | 24 | 22.81 | 29.2 | 30.85 |
| all QMD | 25.1 | Douglas-fir | 136 | 2to84 | 22.68 | 18 | 51.71 | 27.4 | 61.44 |
| SrvyYr | 1856 | hemlock | 36 | 3to30 | 12.22 | 10 | 13.69 | 14.4 | 4.47 |
| | | madrone | 5 | 8to40 | 16.6 | 12 | 1.90 | 20.5 | 1.27 |
| | | bl maple | 2 | 14to28 | 21 | 21 | 0.76 | 22.1 | 0.59 |
| | | grand fir | 1 | 14 | 14 | 14 | 0.38 | 14 | 0.12 |
| | | willow | 4 | 2ro4 | 2.75 | 2.5 | 1.52 | 2.87 | 0.02 |

| Twp-Range | SanJuan County | species | # WTs | Diam Range | av Diam | Median Diam | rel Freq% | QMD | rel DOM% |
|---|---|---|---|---|---|---|---|---|---|
| # allWTs | 1378 | red alder | 219 | 3to36 | 8.288 | 8 | 15.89 | 9.22 | 3.55 |
| all avDiam | 15.1 | ash | 1 | 18 | 18 | 18 | 0.07 | 18 | 0.06 |
| all QMD | 19.5 | redcedar | 54 | 4to48 | 18.19 | 18 | 3.92 | 20.4 | 4.29 |
| SrvyYr | 1874 | cherry | 2 | 3.5to8 | 5.75 | 5 | 0.15 | 6.17 | 0.01 |
| | | cottonwood | 2 | 4to12 | 8 | 8 | 0.15 | 8.94 | 0.03 |
| | | crabapple | 4 | 4to6 | 5 | 5 | 0.29 | 5.1 | 0.02 |
| | | Douglas-fir | 809 | 2to60 | 19.4 | 18 | 58.71 | 23.8 | 87.10 |
| | | grand fir | 5 | 6to30 | 18.8 | 24 | 0.36 | 20.9 | 0.42 |
| | | hemlock | 26 | 3to24 | 8.73 | 7.5 | 1.89 | 10 | 0.50 |
| | | bl maple | 9 | 12to30 | 18 | 18 | 0.65 | 18.7 | 0.60 |
| | | madrone | 20 | 3to10 | 5.6 | 5 | 1.45 | 5.98 | 0.14 |
| | | oak | 8 | 5to32 | 20.38 | 19 | 0.58 | 22.4 | 0.76 |
| | | pine | 126 | 3to20 | 6.9 | 6 | 9.14 | 7.66 | 1.41 |
| | | spruce | 11 | 6to30 | 13.27 | 12 | 0.80 | 14.7 | 0.45 |
| | | willow | 80 | 3to14 | 5.969 | 6 | 5.81 | 6.37 | 0.62 |
| | | yew | 2 | 10 | 10 | 10 | 0.15 | 10 | 0.04 |

| Twp-Range | NonSJCo "SJ"Isls | species | # WTs | Diam Range | av Diam | Median Diam | rel Freq% | QMD | rel DOM% |
|---|---|---|---|---|---|---|---|---|---|
| # allWTs | 361 | red alder | 54 | 2to24 | 8.1 | 7 | 14.96 | 9.2 | 2.77 |
| all avDiam | 17.3 | redcedar | 55 | 4to60 | 21.7 | 18 | 15.24 | 25.5 | 21.70 |
| all QMD | 21.4 | cherry | 4 | 7to12 | 9 | 8.5 | 1.11 | 9.2 | 0.21 |
| SrvyYr | 1871 | Douglas-fir | 183 | 4to72 | 20.6 | 18 | 50.69 | 24.6 | 67.20 |
| | &73 | grand fir | 8 | 10to30 | 20.3 | 21 | 2.22 | 21.1 | 2.16 |
| | | hemlock | 22 | 5to24 | 12.9 | 12.5 | 6.09 | 13.8 | 2.54 |
| | | juniper | 2 | 12to15 | 13.5 | 13.5 | 0.55 | 13.6 | 0.22 |
| | | bl maple | 9 | 3to20 | 10.4 | 10 | 2.49 | 11.7 | 0.75 |
| | | madrone | 1 | 7 | 7 | 7 | 0.28 | 7 | 0.03 |
| | | pine | 4 | 6to18 | 14.3 | 18 | 1.11 | 15.1 | 0.55 |
| | | spruce | 1 | 42 | 42 | 42 | 0.28 | 42 | 1.07 |
| | | vine maple | 2 | 3to4 | 3.5 | 3.5 | 0.55 | 3.5 | 0.01 |
| | | willow | 16 | 5to24 | 7.6 | 6 | 4.43 | 8.8 | 0.75 |
